# Orthogonal CRISPR screens to identify transcriptional and epigenetic regulators of human CD8 T cell function

**DOI:** 10.1101/2023.05.01.538906

**Authors:** Sean R. McCutcheon, Adam M. Swartz, Michael C. Brown, Alejandro Barrera, Christian McRoberts Amador, Keith Siklenka, Lucas Humayun, James M. Isaacs, Timothy E. Reddy, Smita Nair, Scott Antonia, Charles A. Gersbach

**Affiliations:** Department of Biomedical Engineering, Duke University, Durham, NC 27708, USA; Center for Advanced Genomic Technologies, Duke University, Durham, NC 27708, USA; Department of Surgery, Duke University Medical Center, Durham, NC 27710, USA; Department of Neurosurgery, Duke University School of Medicine, Durham, NC 27710, USA; Department of Biostatistics and Bioinformatics, Duke University Medical Center, Durham, NC 27710, USA; Department of Pharmacology and Cancer Biology, Durham, NC 27710, USA; Duke Cancer Institute Center for Cancer Immunotherapy, Duke University School of Medicine, Durham, NC 27710, USA; Department of Pathology, Duke University School of Medicine, Durham, NC 27710, USA

## Abstract

The clinical response to adoptive T cell therapies is strongly associated with transcriptional and epigenetic state. Thus, technologies to discover regulators of T cell gene networks and their corresponding phenotypes have great potential to improve the efficacy of T cell therapies. We developed pooled CRISPR screening approaches with compact epigenome editors to systematically profile the effects of activation and repression of 120 transcription factors and epigenetic modifiers on human CD8+ T cell state. These screens nominated known and novel regulators of T cell phenotypes with BATF3 emerging as a high confidence gene in both screens. We found that BATF3 overexpression promoted specific features of memory T cells such as increased IL7R expression and glycolytic capacity, while attenuating gene programs associated with cytotoxicity, regulatory T cell function, and T cell exhaustion. In the context of chronic antigen stimulation, BATF3 overexpression countered phenotypic and epigenetic signatures of T cell exhaustion. CAR T cells overexpressing BATF3 significantly outperformed control CAR T cells in both in vitro and in vivo tumor models. Moreover, we found that BATF3 programmed a transcriptional profile that correlated with positive clinical response to adoptive T cell therapy. Finally, we performed CRISPR knockout screens with and without BATF3 overexpression to define co-factors and downstream factors of BATF3, as well as other therapeutic targets. These screens pointed to a model where BATF3 interacts with JUNB and IRF4 to regulate gene expression and illuminated several other novel targets for further investigation.

## Introduction

Adoptive T cell therapy (ACT) holds tremendous potential for cancer treatment by redirecting T cells to cancer cells via expression of engineered receptors that recognize and bind to tumor-associated antigens. Receptor-antigen interactions can initiate complex transcriptional networks that drive multipotent T cell response and lead to cancer cell death. The potency and duration of T cell response are associated with defined T cell subsets, and cell products enriched in stem or memory T cells provide superior tumor control in animal models and in the clinic^1–5^. Given the association between defined T cell subsets and clinical outcomes, precise regulation or programming of T cell state is a promising approach to improve the therapeutic potential of ACT.

T cell state and function are largely regulated by specific transcription factors (TFs) and epigenetic modifiers that process intrinsic and extrinsic signals into complex and tightly controlled gene expression programs. For example, TOX^6–10^ and NFAT^11^ program CD8+ T cell exhaustion in the context of chronic antigen exposure. Conversely, T cell function can be enhanced by rewiring transcriptional networks through either enforced expression or genetic deletion of specific TFs and epigenetic modifiers. Ectopic overexpression of specific TFs such as c-JUN^12^, BATF^13^, and RUNX3^14^ or genetic deletion of NR4A^15^, FLI1^16^, members of the BAF chromatin remodeling complex^17, 18^, and regulators of DNA methylation^19, 20^ can alter T cell state and improve T cell function through diverse mechanisms.

Large-scale CRISPR knockout (CRISPRko)^21–23^ and open reading frame (ORF) overexpression^24^ screens have further accelerated gene discovery and defined the effects of individual genes on T cell proliferation and cytokine production. Compared to other screening modes, it has been more challenging to conduct gene activation and repression screens via epigenome editing in primary human T cells^25^. One study optimized lentiviral production to overcome limitations of delivering large CRISPR-based epigenome editors and then conducted proof-of-concept gene silencing and activation screens to define regulators of cytokine production^25^. Indeed, CRISPR-based and ORF genetic screens in primary human T cells use proliferation or cytokine production as the primary readout. Although these are important phenotypic indicators of T cell function, these readouts are also susceptible to missing genes that impact T cell state without significantly changing T cell survival, proliferation, or cytokine production. For example, several key regulators of memory T cells such as TCF7, MYB, and FOXO1 have not emerged from proliferation-based screens. Additionally, combinatorial perturbations and dissection of gene interactions that control human T cell phenotypes have not been extensively explored.

In this study, we developed compact *Staphylococcus aureus* Cas9 (SaCas9)-based epigenome editors for targeted gene silencing and activation in primary human T cells. We leveraged these tools to profile the effects of 120 genes with complementary CRISPR interference (CRISPRi) and activation (CRISPRa) screens on human CD8+ T cell state. These screens and subsequent validation revealed that BATF3 overexpression could be harnessed to support specific features of memory T cells, counter T cell exhaustion, and improve tumor control. By conducting parallel pooled CRISPRko screens of all human transcription factor genes (TFome) with or without BATF3 overexpression, we defined co-factors and downstream targets of BATF3. More generally, we developed orthogonal CRISPR-based screening approaches to systematically discover regulators of complex T cell phenotypes, which should accelerate efforts to engineer T cells with enhanced durability and therapeutic potential.

## Results

### Development and characterization of compact and efficient dSaCas9-based epigenome editors for targeted gene regulation in primary human T cells

SaCas9 has been extensively used for genome editing in vivo as its compact size (3,159 bp) enables packaging into adeno-associated virus (AAV)^26–28^. However, SaCas9 has been used sparingly as an epigenome editor for targeted gene regulation^29, 30^ and has not been used in the context of an epigenome editing screen. First, we evaluated dSaCas9 for targeted gene silencing in primary human T cells by conducting two high-throughput promoter tiling CRISPRi screens. Previous CRISPRi/a screens with dSpCas9 performed serial transductions with one lentivirus encoding the dCas9-effector and another lentivirus encoding for the gRNA-library^25^. To minimize the number of transduction events, we constructed an all-in-one CRISPRi lentiviral plasmid encoding for dSaCas9 fused to the KRAB repressor domain and a gRNA cassette (Figure 1A).

**Figure 1.**
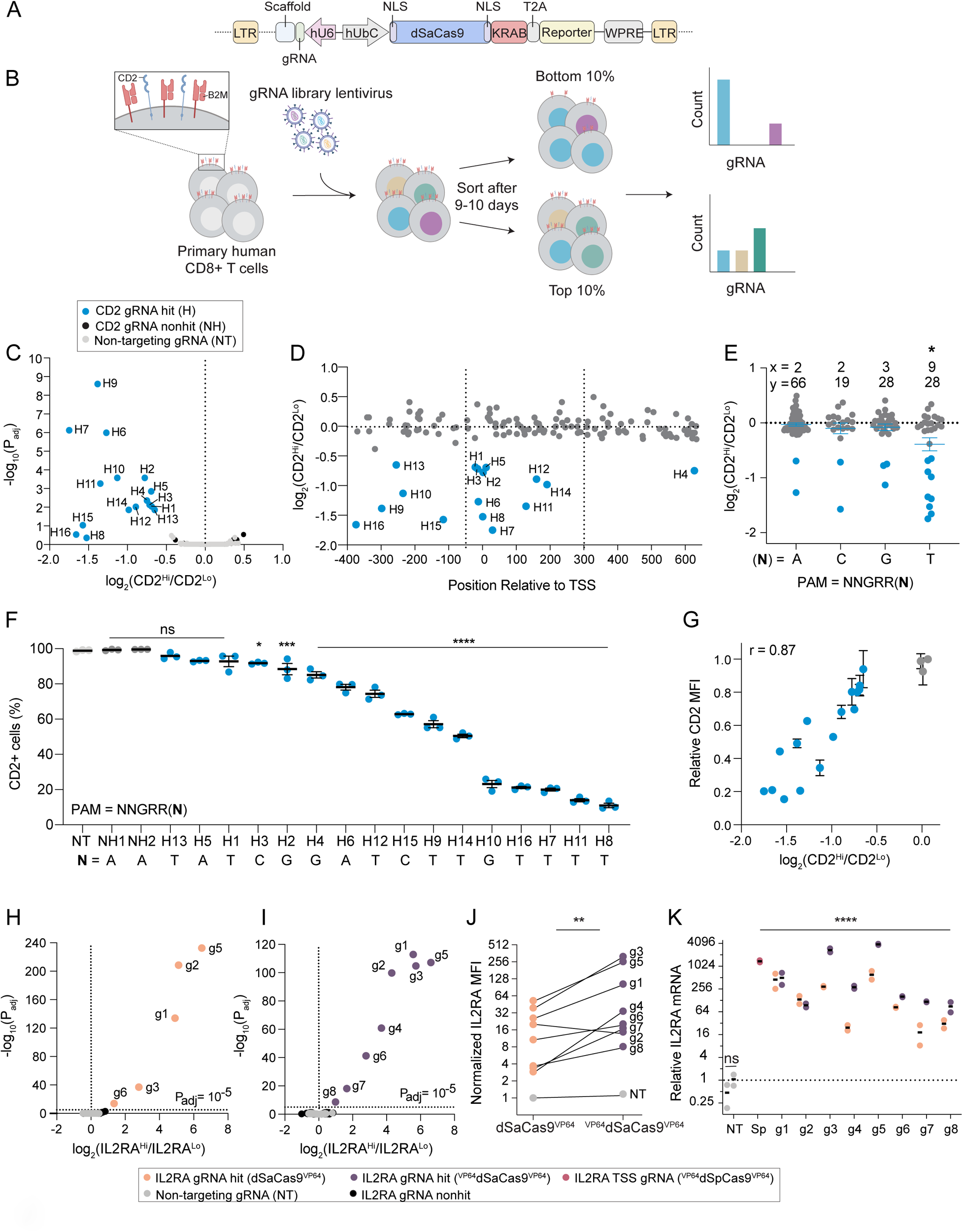
Compact and efficient dSaCas9 epigenome editors for targeted gene silencing and activation. **(A)** Schematic of all-in-one lentiviral plasmid encoding for dSaCas9^KRAB^ and a gRNA cassette. **(B)** Schematic of CD2 and B2M promoter tiling CRISPRi screens in human CD8+ T cells. **(C)** Volcano plot of significance (P_adj_) versus fold change in gRNA abundance between CD2-high and CD2-low populations for the CD2 CRISPRi screen. Blue data points indicate CD2 gRNA hits with a P_adj_ < 0.05 or log_2_(fc) < -1. Black data points indicate non-significant CD2 gRNAs and gray data points indicate the 250 non-targeting (NT) gRNAs. **(D)** CD2 gRNA fold change versus gRNA position relative to the transcriptional start site (TSS). Dashed lines represent the previously defined optimal window^32^ (-50 to +300 bp of TSS) for CRISPRi. **(E)** CD2 gRNA fold change as a function of the final base pair of the PAM (5’-NNGRRN-3’). x represents the number of gRNA hits and y represents the total number of gRNAs in the library for each PAM variant. A one-way ANOVA with Dunnett’s post hoc test was used to compare the average fold change of gRNAs for each PAM variant to NNGRRT (* < 0.05 denotes that the fold change of gRNAs targeting NNGRRT PAMs was significantly different than all other PAM variants). **(F)** Validation of CD2 gRNA hits. Percentage of CD2 positive cells on day 9 post-transduction plotted in rank order based on the mean gRNA activity (n = 3 replicates of CD8+ T cells from pooled PBMC donors, error bars represent SEM). A one-way ANOVA with Dunnett’s post hoc test was used to compare the mean percentage of CD2 positive cells for each gRNA to NT. The final base pair of the PAM for each gRNA is indicated beneath the gRNA label. **(G)** Relationship between CD2 gRNA activity and fold enrichment in screen. Relative CD2 mean fluorescent intensity (MFI) was calculating by normalizing the MFI of each gRNA to the MFI of the NT population. Pearson’s correlation coefficient (r) is indicated in the upper left. Volcano plots of significance (P_adj_) versus fold change in gRNA abundance between IL2RA-high and IL2RA-low populations for the IL2RA CRISPRa Jurkat screens (n = 3 replicates) with **(H)** dSaCas9^VP64^ and **(I)** ^VP64^dSaCas9^VP64^. Yellow and purple data points indicate respective gRNA hits (P_adj_ < 10^-5^). Black data points indicate non-significant IL2RA gRNAs and gray data points indicate the 94 NT gRNAs. **(J)** Normalized IL2RA MFI of dSaCas9^VP64^ and ^VP64^dSaCas9^VP64^ Jurkat lines transduced with indicated gRNAs (n = 2 replicates). Each gRNA was normalized to the IL2RA MFI of the dSaCas9^VP64^ Jurkat line transduced with NT. A paired ratio t-test was used to compare gRNA activity between dSaCas9^VP64^ and ^VP64^dSaCas9^VP64^ Jurkat lines. **(K)** Relative IL2RA mRNA expression of Jurkat CRISPRa lines transduced with indicated gRNA on day 9 post-transduction (n = 2, error bars represent SEM). A one-way ANOVA with Dunnett’s post hoc test was used to compare each gRNA to the NT.

Next, we considered the protospacer adjacent motif (PAM) requirement for SaCas9. In the context of nuclease activity, SaCas9 is more active when targeting genomic regions upstream of the PAM (5’ – NNGRRT-3’) compared to the more relaxed PAM (5’ – NNGRRV – 3’; where V = A, C, or G)^26^. However, several gRNA design tools do not require a thymine in the final position of the PAM^31, 32^. Moreover, the PAM preference for dSaCas9-based epigenetic effectors has not been rigorously characterized. To systematically evaluate this feature, we designed two independent gRNA libraries with the relaxed PAM variant (5’-NNGRRN-3’) and tiled ∼1,000 bp windows around the promoters of *CD2* and *B2M*. We chose *CD2* and *B2M* as gene targets because both are ubiquitously and high expressed genes encoding for surface markers and thus readily compatible with cell sorting-based screens. The CD2 and B2M gRNA libraries contained 141 and 217 targeting gRNAs, respectively, and 250 non-targeting gRNAs.

For each CRISPRi screen, we transduced primary human CD8+ T cells with the respective gRNA library and expanded the cells for 9-10 days before staining and sorting transduced cells in the lower and upper 10% tails of CD2 or B2M expression (Figure 1B). We recovered 16 and 5 targeting gRNAs enriched in the CD2 low and B2M low populations, respectively (Figure 1C and S1A). Many enriched gRNAs were within an optimal window relative to the transcriptional start site (TSS) for gene silencing^32^ (Figure 1D and S1B). Although only a small fraction of the targeting gRNAs (11% of CD2 gRNAs and 2% of B2M gRNAs) were hits for each gene target, the gRNA hit rates (32% of CD2 gRNAs and 16% of B2M gRNAs) were significantly higher for gRNAs targeting the strict PAM (5’-NNGRRT-3’). This is consistent with previous PAM characterization of SaCas9 for nuclease activity^26^ and suggests that the thymine base in the final position of the PAM facilitates more efficient recognition and binding between dSaCas9 and the target DNA sequence (Figure 1E and S1C).

Subsequent validation of CD2 and B2M gRNA screen hits revealed marked gene silencing and a wide range of activity across gRNAs, underscoring the unique capability of CRISPRi to tune gene expression levels (Figure 1F, S1D-E, and S2A-B). For example, the percentage of CD2 silenced cells varied from 7% to 89% depending on the gRNA (Figure 1F and S2A). The mean expression of CD2 in silenced cells was highly correlated with the percentage of silenced cells, indicating that the effect of a gRNA across a cell population is coupled to the magnitude of gRNA activity at a single cell level (Figure S2C). The most potent CD2 and B2M gRNAs targeted genomic sites adjacent to 5’-NNGRRT-‘3 PAMs (Figure 1F and S1D). As previously observed^33, 34^, individual gRNA activity was strongly correlated with its fold-enrichment in the screen (Figure 1G and S2D). Finally, we adapted this CRISPRi system for multiplex gene silencing by using a lentiviral plasmid with orthogonal mouse and human U6 promoters. We verified this system using the most potent CD2 and B2M gRNAs and only detected dual silenced cells when both CD2 and B2M gRNAs were delivered (Figure S2E-G).

Next, we developed efficient and compact dSaCas9-based activators using the small transactivation domain VP64. Using polyclonal Jurkat cell lines constitutively expressing dSaCas9 fused to either one copy of VP64 (dSaCas9^VP64^) or two copies of VP64 (^VP64^dSaCas9^VP64^), we conducted parallel CRISPRa screens with a 400 gRNA library (306 *IL2RA* gRNAs and 94 non-targeting gRNAs) tiling a 5,000 bp window around the TSS of the transcriptionally silenced *IL2RA* gene (Figure S3A). Interestingly, there were three more gRNA hits in the ^VP64^dSaCas9^VP64^ CRISPRa screen along with a shared set of five gRNA hits (Figure 1H-I). All gRNA hits targeted sites within a prominent open chromatin peak within 350 bp of the TSS with the majority located upstream of the TSS (Figure S3B). As with gene silencing, there was a marked preference for 5’-NNGRRT-3’ PAMs for gene activation with 75% of gRNA hits targeting this PAM variant (Figure S3C). Together, the relative gRNA position and PAM sequence were major predictors of gRNA efficacy, similar to the CRISPRi screen, as 24% (6/25) of 5’-NNGRRT-3’ targeting IL2RA gRNAs within a ∼1,000 bp window around the TSS were hits. Individual validation of all eight gRNA hits in both cell lines showed a significant increase in IL2RA expression with ^VP64^dSaCas9^VP64^ consistently more potent than dSaCas9^VP64^ (Figure 1J-K and S3D). Moreover, the most potent ^VP64^dSaCas9^VP64^ gRNAs achieved equivalent levels of IL2RA gene activation as ^VP64^dSpCas9^VP64^ paired with the best IL2RA gRNA from a published CRISPRa screen tiling the *IL2RA* locus in Jurkats^35^ (Figure 1K).

Given the robust activity of ^VP64^dSaCas9^VP64^ in Jurkat cells, we constructed an all-in-one CRISPRa lentiviral vector encoding for ^VP64^dSaCas9^VP64^ and gRNA cassette for assays in primary human T cells. We hypothesized that the smaller size of dSaCas9 would lead to higher titer lentivirus than S. *pyogenes* Cas9 (SpCas9), thus reducing the quantity of T cells and reagents required to perform CRISPR-based screens with equivalent coverage. We tested this by transducing CD8+ T cells from two donors with serial titrations of all-in-one ^VP64^dSaCas9^VP64^ and ^VP64^dSpCas9^VP64^ lentiviruses. ^VP64^dSaCas9^VP64^ produced nearly two-fold higher lentiviral titers than the equivalent ^VP64^dSpCas9^VP64^ construct, thus requiring half the lentiviral volume to achieve the same transduction rate (Figure S4A-D). We then verified that ^VP64^dSaCas9^VP64^ could potently activate endogenous gene expression of a transcriptionally silenced gene (*EGFR*) in primary human T cells (Figure S4E-F).

### CRISPR interference and activation screens identify transcriptional and epigenetic regulators of human CD8 T cell state

Transcription factors (TFs) and epigenetic modifiers function to establish and maintain cell-type specific gene expression programs, mediate response to internal and external stimuli, and ultimately dictate cell fate and function. We therefore sought to interrogate this important class of genes using high-throughput CRISPRi and CRISPRa screens in primary human CD8+ T cells. We compiled a curated list of 110 TFs associated with T cell state and function based on motif enrichment in differentially accessible chromatin across T cell subsets^4,36–38^ and manually appended the following 11 transcriptional and epigenetic regulators: *BACH2, TOX, TOX2, PRDM1, KLF2, BMI1, DNMT1, DNMT3A, DNMT3B, TET1,* and *TET2* for a total of 121 candidate genes (Supplementary Table 2). Based on our characterization of dSaCas9-based epigenome editors, we generated a gRNA library containing all specific, 5’-NNGRRT-3’ PAM targeting gRNAs within a 1,000 bp window centered around the TSS of each gene. All genes were represented by at least 7 gRNAs with an average of 16 gRNAs per gene, except for PBX2 which did not have any gRNAs (Figure S5A-B). We added 120 non-targeting gRNAs as negative controls, bringing the final gRNA library to 2,099 gRNAs (Supplementary Table 2). We cloned the gRNA library into both all-in-one CRISPRi and CRISPRa lentiviral plasmids. Subsequent lentiviral titrations revealed a dose-dependent response to lentiviral volume with both CRISPRi and CRISPRa constructs eclipsing 90% transduction rates (Figure S5C).

We selected CCR7 as the readout for our screens for several reasons. First, CCR7 is a well-characterized T cell marker and is highly expressed in specific T cell subsets such as naïve, stem-cell memory, and central memory T cells^39^. Second, we hypothesized it would enable us to capture more subtle changes in T cell state than other phenotypic readouts such as proliferation or cytokine production. To start with a homogenous T cell population for our screens, we sorted CD8^+^CCR7^+^ T cells from 2-3 donors and transduced each donor with CRISPRi and CRISPRa gRNA libraries at a low multiplicity of infection (MOI) to ensure that most cells only received a single gRNA (Figure S6A-B). We expanded the cells for 10 days post-transduction to allow enough time for both perturbation of the target gene and any downstream effects on gene regulatory networks, and then sorted transduced cells based on expression of CCR7 (Figure 2A and Figure S6C-D).

**Figure 2.**
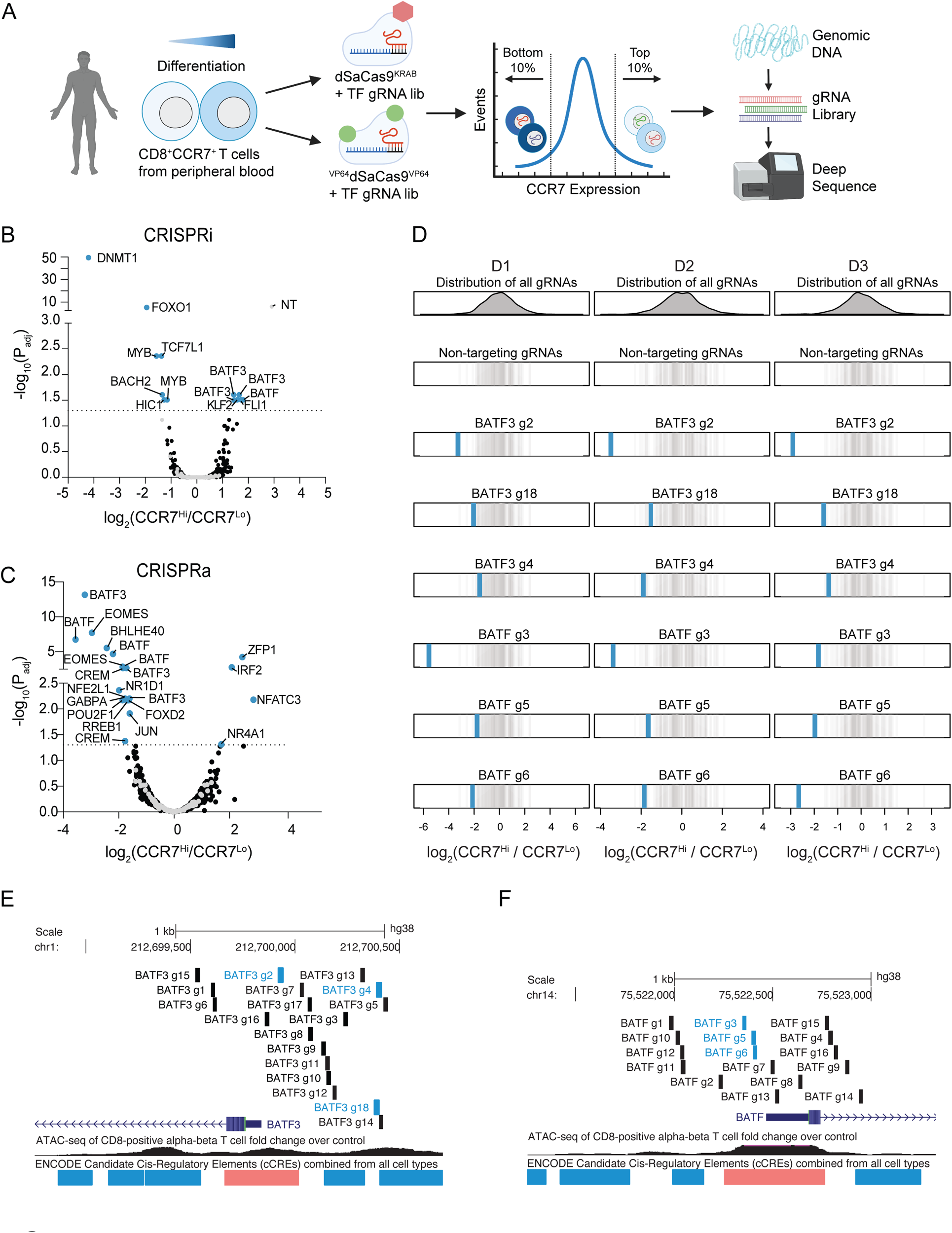
CRISPR interference and activation gene screens identify transcriptional and epigenetic regulators of human CD8+ T cell state. **(A)** Schematic of CRISPRi/a TF screens. Volcano plots of significance (P_adj_) versus fold change in gRNA abundance between CCR7-high and CCR7-low populations for the **(B)** CRISPRi and **(C)** CRISPRa screens. Blue data points indicate gRNA hits (P_adj_ < 0.05) and are annotated with their target gene. Black and gray data points represent non-significant gRNAs and non-targeting gRNAs. **(D)** Fold change of BATF3 and BATF CRISPRa gRNA hits for each donor. Blue vertical lines represent BATF3 or BATF gRNAs and gray vertical lines represent the distribution of 120 non-targeting gRNAs. All **(E)** BATF3 and **(F)** BATF CRISPRa gRNAs in gRNA library relative to TSS, chromatin accessibility, and cCREs. Blue and black vertical lines represent gRNA hits and non-significant gRNAs, respectively.

The CRISPRi screen recovered many canonical regulators of memory T cells including *FOXO1*^40^, *MYB*^41^, and *BACH2*^42^ – all of which when silenced led to reduced expression of CCR7, indicative of T cell differentiation towards effector T cells (Figure 2B and Figure S7A). Interestingly, the most significant hit from the CRISPRi screen was *DNMT1*, which encodes for a DNA methyltransferase that maintains DNA methylation across cell divisions via recognition of hemi-methylated DNA. Genetic disruption of both *TET2* and *DNMT3A*, which encode for proteins that regulate DNA methylation in opposite directions, can improve the therapeutic potential of T cells^19, 20^. There was a single non-targeting gRNA (1/120) hit in the CRISPRi screen. The same non-targeting gRNA emerged as a hit in multiple screens using CCR7 as the readout, suggesting a real off-target effect.

The CRISPRa screen identified several transcription factors that have been implicated in CD8+ T cell differentiation and function such as *EOMES*^43^, *BATF*^13^, and *JUN*^12^ (Figure 2C). Importantly, gRNA enrichment was consistent across the three donors (Figure 2D and Figure S7B). Multiple gRNAs targeting basic leucine zipper ATF-like transcription factors *BATF* and *BATF3* were enriched in reciprocal directions across CRISPRi and CRISPRa screens, highlighting the power of coupling loss- and gain-of-function perturbations, and this was not related to the number of library gRNAs targeting these genes (Figure S7C-D). We noticed that *BATF* and *BATF3* gRNA hits in the CRISPRa screen generally co-localized to regions upstream of the promoter and near the summits of accessible chromatin (Figure 2E-F). This observation is consistent with our CRISPRa IL2RA gRNA tiling screens and recent recommendations for designing gRNAs for *cis*-regulatory elements.

### Single cell RNA-seq characterization of transcriptional and epigenetic regulators of T cell state

We next characterized the transcriptomic effects of each candidate gene identified from our CRISPRi and CRISPRa screens using single cell RNA-seq (scRNA-seq). We adapted the ECCITE-seq protocol^44^ for SaCas9 by designing a reverse transcription primer complementary to the constant scaffold region of the SaCas9 gRNA. This enabled simultaneous capture of both non-polyadenylated gRNA transcripts and mRNA transcripts from individual cells. We cloned the union set of gRNA hits across CRISPRi/a screens (32 gRNAs) and 8 non-targeting gRNAs into both CRISPRi and CRISPRa plasmids. We then followed the same workflow as the sort-based screens, but instead of sorting the cells based on CCR7 expression, we profiled the transcriptomes and gRNA identity of ∼60,000 cells across three donors for each screen. After filtering for high-quality, gRNA-assigned cells, we aggregated the cells and compared the transcriptional profile of cells with the same gRNA to non-perturbed cells (cells with only non-targeting gRNAs). To assess the quality and statistical power of our scRNA-seq data, we compared the quantity and magnitude of effects between targeting and non-targeting gRNAs. Targeting gRNAs were associated with significantly more differentially expressed genes (DEGs) and these gRNA-to-gene links had larger effect sizes than non-targeting gRNAs (Figure S8A-D). We therefore proceeded to evaluate the transcriptomic effects of silencing or activating each candidate gene.

First, we focused on CCR7 expression across gRNAs to validate the results from our CRISPRi/a sort-based screens (Figure 3A-B). The scRNA-seq data revealed that half of the gRNA hits reproducibly affected CCR7 expression with a similar rank order as predicted by the sort-based screens. For example, both assays informed that targeted silencing of DNMT1 or FOXO1 drastically reduced CCR7 expression levels, which was further confirmed through individual gRNA validations (Figure S9A-B). We noticed that gRNA hits that failed to validate in the scRNA-seq characterization were represented by fewer cells than gRNAs that did successfully validate, reaffirming that higher gRNA coverage helps to resolve more subtle changes in gene expression^44^ (Figure S9C). For example, the CRISPRa CREM-targeting gRNA that validated was represented by 635 cells, whereas the other CREM-targeting gRNA was represented only 142 cells and narrowly missed the significance threshold. Nevertheless, both gRNAs upregulated CREM and similarly affected gene expression programs (Figure S9D). Several BATF-targeting gRNAs were also underrepresented. To evaluate CREM and BATF gRNAs, we individually assayed a pair of CRISPRa gRNA hits targeting each gene and validated that each gRNA regulated CCR7 expression as predicted by the screen (Figure S9A-B). We suspect that several gRNAs were underrepresented in the scRNA-seq experiment due to gRNA-intrinsic features that either interfered with reverse transcription, oligo capture by the beads, or subsequent amplification. Underrepresented gRNAs were not depleted in the initial gRNA plasmid pool nor did we observe any fitness defects in individual validations. The same gRNAs were underrepresented in both CRISPRi and CRISPRa assays, further pointing to gene-independent effects. In addition to the scRNA-seq data confirming predicted gRNA effects on CCR7 expression, the true negative rates were high for both CRISPRi (96%) and CRISPRa (82%), demonstrating the specificity of sort-based screens (Figure 3A-B).

**Figure 3.**
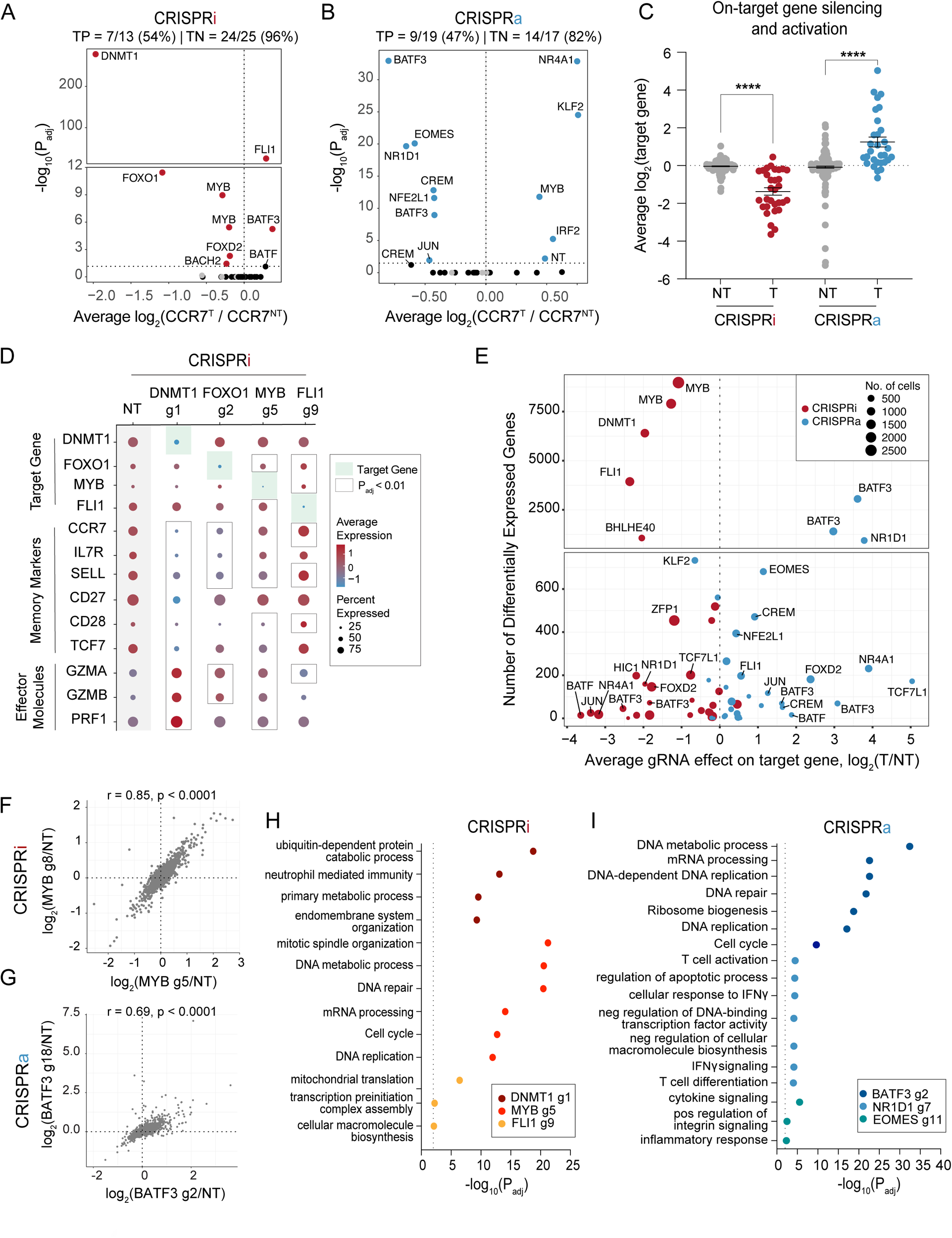
Single cell RNA-sequencing characterization of gene candidates. Volcano plots of significance (P_adj_) versus average fold change of CCR7 expression for each gRNA compared to non-perturbed cells for **(A)** CRISPRi and **(B)** CRISPRa perturbations. Red and blue data points indicate gRNA hits (P_adj_ < 0.05). Gray data points indicate NT gRNAs. True positive and negative rates are displayed above each volcano plot. **(C)** Average fold change in target gene expression for non-targeting gRNAs and targeting gRNAs across CRISPRi and CRISPRa perturbations. A two-way ANOVA with Tukey’s post hoc test was used to compare the fold change in target gene expression between groups. **(D)** Dot plot depicting the average expression and percent of cells expressing target genes, memory markers, and effector molecules for the indicated CRISPRi perturbations. **(E)** Scatter plot of the number of differentially expressed genes (DEGs defined as P_adj_ < 0.01) associated with each gRNA versus the gRNA effect on the target gene for both CRISPRi and CRISPRa perturbations. **(F)** Correlation of the union set of DEGs between the top CRISPRi MYB gRNAs. **(G)** Correlation of the union set of DEGs between the top CRISPRa BATF3 gRNAs. Representative enriched pathways for the top three **(H)** CRISPRi and **(I)** CRISPRa gRNAs.

We next measured on-target gene silencing or activation to confirm downstream-mediated effects, such as changes in CCR7 expression, were driven by each candidate gene. Of CRISPRi and CRISPRa gRNAs assigned to at least 5 cells, 56/61 gRNAs (92%) silenced or activated their gene target (Figure 3C). Given that CCR7 was selected as a surrogate marker for a memory T cell phenotype, we expected some perturbations to regulate subset-defining gene expression programs. Indeed, scRNA-seq revealed that silencing the top predicted positive regulators of memory (*DNMT1*, *FOXO1*, *MYB*) led to decreased expression of CCR7 and other memory-associated genes (such as *IL7R*, *SELL*, *CD27*, *CD28*, *TCF7*) and increased expression of effector-associated genes (*GZMA*, *GZMB*, *PRF1*) (Figure 3D). Conversely, silencing *FLI1* led to increased expression of several memory-associated genes. Finally, we examined all DEGs associated with each perturbation to gain an unbiased view of the transcriptomic effects of silencing and activating each gene. Endogenous regulation of several TFs and epigenetic-modifying proteins had widespread transcriptional effects with 6 gene perturbations (4 CRISPRi gene perturbations and 2 CRISPRa gene perturbations) altering expression of >1,000 genes (Figure 3E). These widespread transcriptional changes were not attributed to impaired cell fitness as these gRNAs were well represented after 10 days of T cell expansion. Unsurprisingly, silencing *DNMT1* – a global epigenetic modifier – massively altered the transcriptome with 6,401 DEGs and affected general biological processes such as metabolism, endomembrane system organization, and mitotic spindle organization (Figure 3H).

Interestingly, *MYB* repression with two unique gRNAs resulted in widespread and concordant gene expression changes with 8,976 and 7,899 DEGs (Figure 3E-F). Mouse models of acute and chronic infection have implicated MYB as an essential positive regulator of stem-like memory CD8+ T cells^41^ and a small and distinct CD62L+ precursor of exhausted T cell population^45^. In both contexts, MYB-deficient CD8+ T cells lacked therapeutic potential due to either impaired recall response or the inability to respond to checkpoint blockage. An important and lingering question has been whether MYB plays a similar role in human CD8+ T cells. Our scRNA-seq data revealed that MYB does indeed regulate human CD8+ T cell stemness with MYB silencing driving CD8+ T cells towards terminal effector T cells. MYB silencing led to downregulation of memory-associated TFs (*TCF7*, *KLF2*), lymph homing molecules (*CCR7*, *CD62L*, *S1PR1*), and cell-cycle inhibitors (*CDKN1B*). In addition, MYB-silenced cells had increased expression of effector-associated TFs (*TBX21*, *PRMD1*, *ZNF683*), effector molecules (*GZMB*, *PRF1*), inflammatory cytokines (*IFNG*, *TNF*), and positive cell-cycle regulators (*E2F1*, *CDC6*, *SKP2*, *CDC25A* and *KIF14*) (Figure S10A-B). The two MYB CRISPRi gRNAs were represented by the first and third most cells across both CRISPRi and CRISPRa screens, suggesting that MYB silencing promoted T cell proliferation (Figure 3E).

Endogenous activation of several TFs including *NR1D1*, *EOMES*, and *BATF3* had large effects on T cell state. Perturbation-driven single cell clustering revealed a distinct cluster with NR1D1 activation (Figure S11A). *NR1D1* encodes a nuclear receptor subfamily 1 transcription factor and negatively regulates expression of core clock proteins that govern cyclical gene expression patterns. Integrative analysis of bulk ATAC-seq data across 12 independent studies of CD8 T cell dysfunction in cancer and infection found that the NR1D1 motif was enriched in open chromatin of exhausted T cells^38^. The causal role of NR1D1 in CD8 T cells, however, has not been studied. NR1D1 activation resulted in 646 upregulated and 293 downregulated genes (Figure S11B). In agreement with NR1D1 motif enrichment^38^, a large set of effector and exhaustion-associated genes were markedly upregulated with NR1D1 activation. To better understand the magnitude of exhaustion induction by NR1D1, we calculated an exhaustion gene signature score using a defined set of 82 exhaustion-specific genes^46^. *NR1D1*-perturbed cells had a significantly higher exhaustion gene signature score than non-perturbed cells (Figure S11C). Many memory-associated surface markers (*IL7R*, *CCR7*, *SELL*, *CD5*) and TFs (*TCF7*, *LEF1*) were downregulated, suggesting NR1D1 activation synthetically programs a transcriptional profile with features of T cell exhaustion.

Endogenous activation of *EOMES*, a regulator of effector T cells, drove markers associated with cytokine signaling and inflammatory response, but did not lead to an increase in exhaustion-related genes (Figure 3I). The top two BATF3 gRNA hits from our cell sorting CRISPRa screen had strong and concordant effects with 3,056 and 1,402 DEGs (Figure 3E, G). Gene ontology analyses revealed that BATF3-induced genes were enriched for DNA and mRNA metabolic processing, ribosomal biogenesis, and cell-cycle pathways, suggesting that BATF3 improves T cell fitness (Figure 3I).

### BATF3 overexpression promotes features of memory T cells and counters signatures of T cell exhaustion

BATF3 has been shown to promote survival and memory formation in mouse CD8+ T cells, however, the molecular and phenotypic effects of BATF3 in human CD8+ T cells has not been well defined^47^. Moreover, it is not known whether manipulating BATF3 expression in CD8+ T cells can improve T cell-mediated control of infection or cancer. To better understand the kinetics of BATF3 expression in human CD8+ cells, we performed a time course experiment where we transduced CD8+ T cells with control lentiviral vector (GFP or CRISPRa + NT gRNA), CRISPRa + BATF3 gRNA, or BATF3 open reading frame (ORF) and measured BATF3 mRNA expression at five different time points (Figure S12A). Consistent with other studies^47, 48^, BATF3 expression levels spiked after T cell activation and tapered back to baseline levels by day 10 post-transduction. Both endogenous activation and ectopic BATF3 expression increased BATF3 levels relative to the controls, however, ectopic expression led to significantly higher levels of BATF3. Given the higher expression and compact size of BATF3’s ORF (only 381 bp), we decided to use ectopic BATF3 expression for all subsequent assays.

First, we found that BATF3 overexpression (OE) markedly increased expression of IL7R, a surface marker associated with T cell survival, long-term persistence, and positive clinical response to ACT^49^ (Figure 4A-B and S12B). Next, we performed RNA-seq across CD8+ T cells from five donors to gain an unbiased view of the transcriptomic changes induced by BATF3 OE. Compared to control cells, there were over 1,100 DEGs distributed almost equally between upregulated and downregulated genes (Figure 4C). Gene ontology analyses revealed that BATF3 OE increased expression of genes involved in metabolic pathways such as glycolysis and gluconeogenesis, T cell proliferation (DNA replication), and translation (Figure 4D and Supplemental Table 4). Metabolic fitness is strongly associated with the potency of T cell responses. For example, central memory CD8+ T cells require glycolysis to mount rapid-recall responses after secondary antigen exposure. TCF1 supports these bioenergetic demands by preprogramming the mobilization of glycolytic enzymes^50^. Interestingly, BATF3 OE increased expression of the transcription factor ID3 (downstream of TCF1), which can activate glycolysis and partially rescue secondary response in the absence of TCF1^50^.

**Figure 4.**
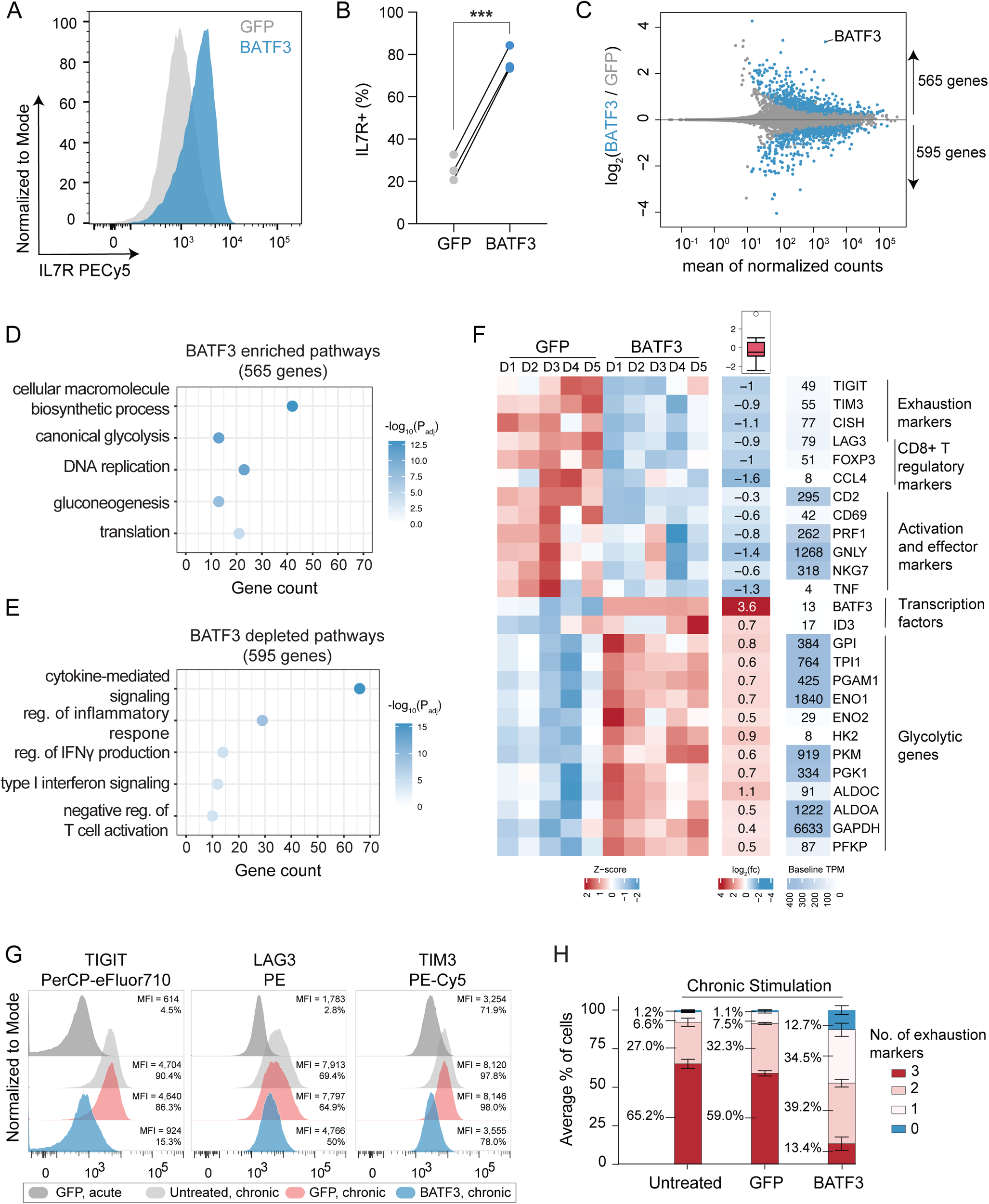
BATF3 overexpression promotes specific features of memory T cells and counters exhaustion and cytotoxic gene signatures. **(A)** Representative histogram of IL7R expression in CD8+ T cells with or without BATF3 overexpression on day 8 post transduction. **(B)** Summary statistics of IL7R expression with or without BATF3 overexpression (n = 3 individual donors, paired t test was used to compare IL7R expression between groups, lines connect the same donor). **(C)** Differential gene expression analysis between CD8+ T cells with or without BATF3 overexpression on day 10 post transduction. Blue data points indicate differentially expressed genes (DEGs, P_adj_ < 0.01, n = 5 donors). **(D)** Selected enriched and **(E)** depleted biological processes from BATF3 overexpression. **(F)** Heatmap of DEGs related to T cell exhaustion, regulatory function, cytotoxicity, transcriptional activity, and glycolysis. (G) Representative histograms of exhaustion markers (TIGIT, LAG3, and TIM3) on day 12 after acute or chronic stimulation across groups. **(H)** Stacked bar chart with average percentage of CD8+ T cells positive for 0, 1, 2, or 3 exhaustion markers (TIGIT, LAG3, TIM3) on day 12 after chronic stimulation across groups (n = 3 independent donors, error bars represent SEM).

In contrast, BATF3 OE dampened T cell effector programs with downregulation of activation markers (*CD69*, *CD2*), inflammatory cytokines and cytotoxic molecules (*TNF*, *PRF1*, *GNLY*, *NKG7*) (Figure 4E-F). Additionally, BATF3 OE reduced expression of several markers associated with regulatory T cells, which have recently emerged as a predictive cell type for clinical response to ACT. In a cohort of refractory B cell lymphoma patients treated with CD19 CAR T cell therapy, the infused T cell product of non-responders were enriched for T_reg_ cells and FOXP3^+^ cells (across all CAR^+^ cells) compared to responders^49^. Albeit less characterized than CD4+ T_regs_, a subset of CD8^+^FOXP3^+^LAG3^+^ T_regs_ suppress T cell activity by secreting CC chemokine ligand 4 (CCL4)^51^. Interestingly, our RNA-seq data revealed that BATF3 OE reduced expression of FOXP3, LAG3, and CCL4 in CD8^+^ T cells (Figure 4F and S12C). BATF3 has previously been shown to silence FOXP3 expression in CD4+ T_regs_ by directly binding to regulatory regions within the *FOXP3* locus^52, 53^.

In addition to LAG3, BATF3 silenced several other canonical markers of T cell exhaustion including TIGIT, TIM3, and CISH (Figure 4F and S12C). We speculated these effects might be amplified in the context of chronic antigen stimulation. To evaluate this, we acutely and chronically stimulated control and BATF3 OE T cells with CD3/CD28 antibody-coated beads and measured expression of exhaustion-associated surface markers (PD1, TIGIT, LAG3, and TIM3) (Figure S13A). As previously observed^54^, PD1 expression peaked after the initial stimulation and then tapered off over time, whereas TIGIT, LAG3, and TIM3 expression were maintained or increased after each subsequent round of stimulation (Figure S13B-C). Notably, BATF3 OE attenuated PD1 induction and restricted TIGIT, LAG3, and TIM3 expression to closely resemble that of acutely stimulated cells despite three additional rounds of TCR stimulation (Figure 4G and S13B-C). As terminally exhausted T cells often co-express multiple exhaustion-associated markers, we quantified the proportion of cells expressing each combination of TIGIT, LAG3, and TIM3. Only 13% of BATF3 OE T cells co-expressed all three markers compared to 65% and 59% of untreated and GFP T cells (Figure 4H).

### BATF3 overexpression remodels the epigenetic landscape of CD8+ T Cells under acute and chronic stimulation

As an orthogonal method of inducing T cell exhaustion, we armed T cells with a human epidermal growth factor 2 (HER2) CAR with or without BATF3 OE and acutely and chronically stimulated the CAR T cells with human HER2+ cancer cells. Using assay for transposase-accessible chromatin with sequencing (ATAC-seq), we profiled the epigenetic landscape of T cells in each group. As expected, chronic antigen stimulation induced widespread changes in chromatin accessibility with 23,322 differentially accessible regions between acutely and chronically stimulated control cells. Many of these regions were proximal to memory and effector/exhaustion-genes (Figure S14A).

Next, we assessed chromatin remodeling in response to BATF3 OE under acute stimulation. There was extensive chromatin remodeling with 5,104 differentially accessible regions between the groups (Figure S15A). Of these regions, roughly 60% were more accessible with BATF3 OE. Most of these changes were in intronic or intergenic regions consistent with *cis*-regulatory or enhancer elements (Figure S15B). To better understand whether changes in chromatin accessibility corresponded to changes in gene expression, we jointly analyzed our ATAC-seq and RNA-seq data. We assigned each differential region to its closest gene to estimate genes that could be regulated in *cis* by these elements. We then quantified how many differential regions proximal to DEGs gained or lost accessibility. There was an enrichment of regions with increased or decreased accessibility proximal to upregulated and downregulated genes, respectively, indicative that BATF3-driven epigenetic changes affected transcription (Figure S15C). Approximately 25% of the genes that changed expression were associated with a corresponding differentially accessible region (297 out of 1,160 genes). For example, BATF3 OE extensively remodeled the chromatin landscape at *IL7R* and *TIGIT* (Figure S15D-E). BATF3 OE increased accessibility at the *IL7R* promoter, intronic, 3’-UTR, and intergenic regions and decreased accessibility at distal intergenic, 5’-UTR, and exonic regions of *TIGIT*.

Finally, we compared the epigenetic landscapes of chronically stimulated T cells with or without BATF3 OE. There were 22,201 differentially accessible regions between control and BATF3 OE T cells with most regions in intronic and intergenic regions (Figure S16A). Interestingly, we observed increased accessibility at regions near both memory (*TCF7*, *MYB*, *IL7R*, *CCR7*, *SELL*) and effector-associated genes (*EOMES*, *TBX21*) (Figure S16C-D). This may represent a hybrid T cell phenotype or the presence of heterogenous subpopulations of memory and effector T cells. Consistent with RNA-seq and FACS data, we observed reduced accessibility at exhaustion loci such as *TIGIT*, *CTLA4*, *LAG3* with BATF3 OE.

### BATF3 overexpression enhances tumor control and programs a transcriptional signature associated with clinical response

Given that BATF3 OE induced widespread changes in gene expression and chromatin accessibility, we hypothesized that BATF3 OE might improve CD8+ T cell function. To test the antitumor capacity of BATF3 OE T cells, we used an in vitro co-culture model with T cells engineered to express a HER2-CAR and human HER2+ cancer cells. We verified that cancer cell death was dependent on the presence of CAR+ T cells. (Figure S17A-B). At sub-curative doses of control CAR T cells, CAR T cells co-expressing BATF3 were more potent tumor killers than control CAR T cells across donors and multiple effector: target (E:T) ratios (Figure 5A and S17B).

**Figure 5.**
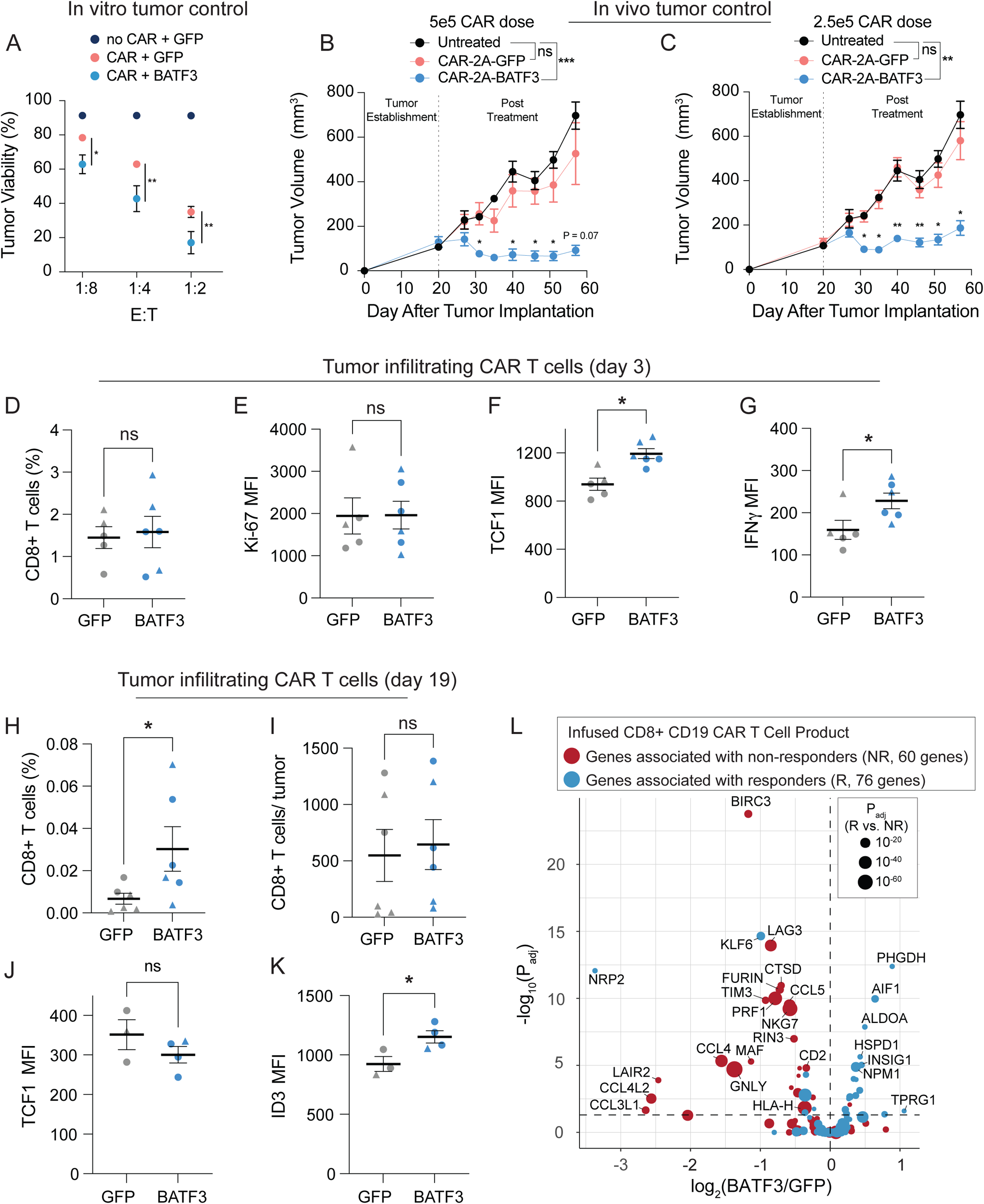
BATF3 OE enhances tumor control in vivo and programs a transcriptional signature associated with clinical response to ACT. **(A**) Tumor viability after 24 hours of co-culture with GFP CAR^null^, GFP CAR+, and BATF3 OE CAR+ CD8 T cells at indicated effector to target (E:T) cell ratios (n = 3 individual donors, error bars represent SEM). A two-way ANOVA with Dunnett’s post hoc test was used to compare tumor viability between GFP+CAR+ and BATF3+CAR+ T cells at each E:T ratio. Tumor volume over time for untreated mice and mice treated with **(B)** 5 x 10^5^ or **(C)** 2.5 x 10^5^ CAR T cells with or without BATF3 overexpression (n = 1 donor, 4-5 mice per treatment, error bars represent SEM). A two-way ANOVA was used to compare the tumor volumes at each time point across treatments. Tumor volumes were not statistically different between untreated and control CAR groups at any time point. Tumor volumes were significantly different between untreated and BATF3 OE CAR groups from day 31 onward. The asterisks above the blue lines indicate significant differences in tumor volumes between mice treated with control and BATF3 OE CAR T cells at each time point. **(D)** Average percentage of CD8+ T cells within each resected, dissociated tumor on day 3 post-treatment (n = 2 donors, 2-3 mice per donor, error bars represent SEM). A Mann-Whitney test was used to compare the percentage of CD8+ cells between groups. **(E-G)** Ki-67, TCF1, and IFNγ MFI of tumor infiltrating CAR T cells on day 3 across groups (n = 2 donors, 2-3 mice per donor, error bars represent SEM). Unpaired t tests were used to compare MFI between groups. **(H)** Average percentage and **(I)** total number of CD8+ T cells within each resected, dissociated tumor on day 19 post-treatment across groups. A Mann-Whitney test was used to compare the percentage and total number of CD8+ cells between the two groups. **(J-K)** TCF1 and ID3 MFI of tumor infiltrating CAR T cells on day 19 across groups (n = 2 donors, 1-3 mice per donor, error bars represent SEM). Unpaired t tests were used to compare MFI between groups. **(L)** Volcano plot of significance (P_adj_) versus fold change between BATF3 OE and control CD8+ T cells for a subset of 144 genes that were negatively (red data points) or positively (blue data points) associated with clinical outcome to CD19 CAR T cell treatment. The size of each data point corresponds to the strength of association between gene expression and clinical response.

Next, we evaluated whether BATF3 OE could improve in vivo control of solid tumors, given the known role of T cell exhaustion in limiting ACT efficacy in the solid tumor setting^55, 56^. To simplify delivery of the CAR and BATF3 transgenes, we constructed all-in-one lentiviral vectors encoding a HER2 CAR coupled to either GFP or BATF3 via a 2A polypeptide skipping sequence. Using an orthotopic human breast cancer HER2+ tumor model in immunodeficient NSG mice, we measured tumor volumes over time as a function of control (GFP) CAR T cell doses (Figure S17C). Tumor control was partial in the cohort of mice treated with 5 x 10^5^ CAR T cells and completely lost in the cohort treated with 10^5^ CAR T cells. We proceeded to test whether BATF3 OE could improve the therapeutic potential of CAR T cells with several sub-curative doses. Strikingly, CAR T cells co-expressing BATF3 markedly enhanced tumor control at two sub-curative doses (2.5 x 10^5^ and 5 x 10^5^ CAR+ cells) compared to control CAR T cells (Figure 5B-C and S17F). Notably, the tumor growth of mice treated with the lower dose of 2.5 x 10^5^ control CAR T cells was completely unrestrained, mimicking that of untreated mice (Figure 5C). In stark contrast, there was clear regression and delay of tumor growth with the matched dose of BATF3 OE CAR T cells.

To explore the mechanism driving superior tumor control with BATF3 OE, we repeated the in vivo experiment with T cells from two different donors and phenotypically characterized the CAR T cells before treatment and after collecting tumor infiltrating CAR T cells on day 3 and day 19 post-treatment (Figure 5D-K, S18-19). Across both sets of experiments, there were no differences in CAR transduction rates (>70% for all groups) or the total number of CAR+ T cells before intravenous injections between CAR constructs (Figure S17D-E). Again, we observed superior tumor control with BATF3 OE CAR T cells across both donors (Figure S18A-B). Although there were no statistically significant differences between the input CAR T cells given the small sample size, BATF3 OE cells tended to express lower levels of several exhaustion markers including LAG3, TIGIT, and TIM3 (Figure S18C-E).

More striking differences between the two groups emerged at the day 3 post-treatment timepoint. We detected equivalent proportions of CD8+ T cells within the tumor and circulating in peripheral blood, indicating that BATF3 OE was not improving tumor control by merely increasing T cell proliferation or tumor trafficking (Figure 5D and S18F). Corroborating this, expression of the proliferative marker Ki-67 was equivalent between the groups (Figure 5E). Rather, tumor infiltrating CAR T cells with BATF3 OE expressed higher levels of both TCF1 and IFNγ (Figure 5F-G). BATF3 OE did not increase expression of TCF7 (which encodes for TCF1) under acute stimulation *in vitro* (Figure S12C). However, there were seven differentially accessible sites near the TCF7 locus between control and BATF3 OE CAR T cells after chronic stimulation (Figure S16C, E). Notably, 5/7 sites were more accessible in BATF3 OE cells including all three intragenic regions, while four distal intergenic regions were split evenly between the two groups (Figure S16E). These data suggest that BATF3 OE can partially counter heterochromatinization of the TCF7 locus during chronic antigen stimulation and retain higher levels of TCF1 expression.

As reflected in the tumor growth curves, we detected a higher proportion of tumor infiltrating CAR T cells in the BATF3 OE group at the final day 19 timepoint, likely due to smaller tumor sizes, as the absolute number of T cells were similar between the two groups (Figure 5H-I). We did not detect any CAR T cells in peripheral blood for either group. To gain further insight into transcriptional regulation, we stained the tumor infiltrating CAR T cells for the following TFs: TCF1, TBET, EOMES, GATA3, ID2, ID3, and IRF4. Interestingly, TCF1 was no longer differentially expressed, but ID3 (a downstream TF of TCF1) was upregulated in the BATF3 OE group (Figure 5J-K). Therefore BATF3 OE T cells may have gradually transitioned from transcriptional programs driven by TCF1 to ID3.

Given the enhanced tumor control conferred by BATF3 OE in CD8+ T cells, we were curious whether BATF3 OE programmed a transcriptional signature associated with clinical response to ACT. Suggestive of this, in a recent clinical trial, non-responders to CD19-targeting CAR T cell therapy had a significantly higher proportion of CD8+ T cells in a cytotoxic or exhausted phenotype than responders^49^. This prompted us to systematically identify DEGs between the infused CD8+ CD19 CAR T cell product of responders and non-responders (Figure S20A). There were 147 DEGs between CD8+ T cells of responders and non-responders in this dataset. We then subset our bulk RNA-seq data with BATF3 OE to query the expression of these genes. Of the 147 DEGs, 144 genes were detected in our RNA-seq data. Strikingly, BATF3 OE silenced 35% (23/65) of genes associated with nonresponse and activated 20% (16/79) of genes associated with response (Figure 5L). Seven of the ten genes most strongly associated with clinical outcome were regulated in a favorable direction. Conversely, only 4.9% (7/144) of genes were regulated in a direction opposing positive clinical response, providing further evidence that BATF3 OE drives a transcriptional program associated with positive clinical outcomes.

### CRISPR knockout screens reveal co-factors of BATF3 and novel targets for cancer immunotherapy

BATF3 is a member of the AP-1 TF family, which regulates diverse biological processes across many cell types through complex and highly specific transcriptional control. This transcriptional specificity is enabled by combinatorial interactions between AP-1 TFs, which form cell-type specific homo- or hetero-dimers to regulate distinct gene expression programs. Several AP-1 complexes such as BATF-JUN heterodimers can interact with interferon-regulatory factors (IRF) at AP-1-IRF consensus elements, providing further flexibility in gene regulation^57^. BATF3 is a compact AP-1 TF with only a basic DNA binding domain and a leucine zipper motif. Unlike other AP-1 TFs, BATF3 lacks additional protein domains such as a transactivation domain for gene activation^57^. We therefore speculated that BATF3 was interacting with other TFs to impact gene expression and chromatin accessibility. Additionally, we reasoned that other TFs might compete with or inhibit BATF3 and that deleting these factors would further amplify BATF3’s effects.

To identify cooperative TFs, downstream factors, and barriers to T cell reprogramming, we conducted parallel CRISPR knockout (CRISPRko) screens with gRNA libraries targeting all human transcription factors genes (TFome), with or without BATF3 OE. We selected IL7R expression as the readout for these screens for two reasons. First, IL7R is expressed in 20-50% of CD8+ T cells at baseline, making it feasible to recover gene hits in both directions, unlike ubiquitously silenced and highly expressed genes. Second, BATF3 OE profoundly increases IL7R expression (Figure 4A-B), thus providing a proxy for BATF3 activity. We expected that IL7R induction by BATF3 would be attenuated if cooperative or downstream TFs were deleted. We designed the TFome gRNA library by subsetting a genome wide knockout library^31^ (4 gRNAs per gene) for 1,612 TFs^58^. We included four IL7R-targeting gRNAs as positive controls and 550 non-targeting gRNAs as negative controls. We cloned the 7,000 gRNA library into two lentiviral plasmids encoding for either mCherry or BATF3 and transduced CD8+ T cells from two donors in parallel with each library. The following day, we electroporated Cas9 protein to facilitate gene editing and then expanded the edited cells^21^. After nine days of expansion, we sorted the cells into the lower and upper 10% tails of IL7R expression and sequenced the gRNA libraries from each population (Figure 6A).

**Figure 6.**
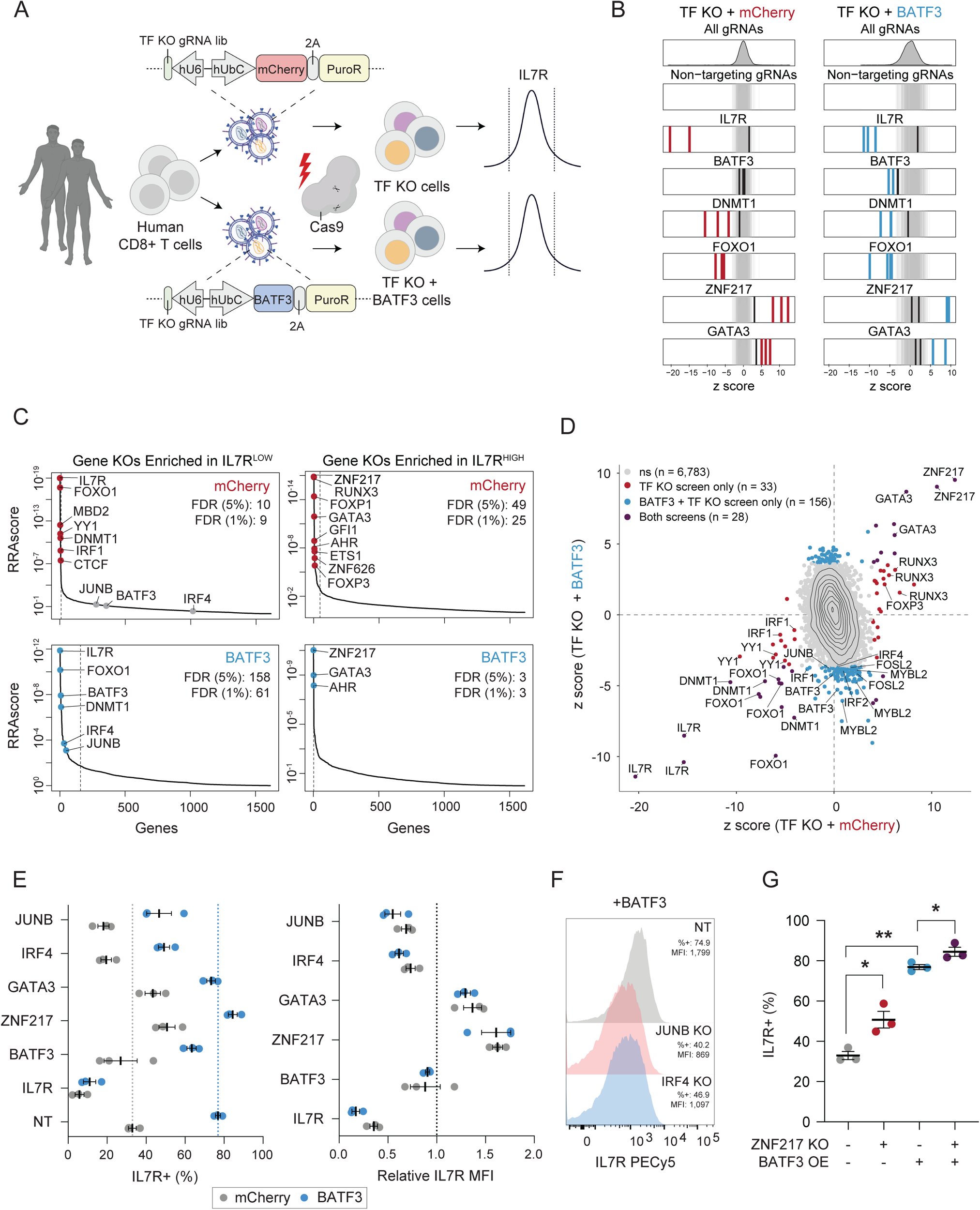
CRISPRko screens reveal co-factors of BATF3 and novel targets for cancer immunotherapy. **(B)** Schematic of CRISPRko screens. **(C)** z scores of gRNAs for selected genes in mCherry (left) and BATF3 (right) screens. Enriched gRNAs (P_adj_ < 0.01) are labeled in red or blue for each screen. Non-targeting gRNAs are labeled in gray. **(D)** Each gene target in the mCherry (top) and BATF3 (bottom) screens ranked based on the MAGeCK robust ranking aggregation (RRA) score in both IL7R^LOW^ (left) and IL7R^HIGH^ (right) populations. Red and blue data points represent enriched genes in each screen. Dashed lines indicate an FDR < 0.05 cutoff. **(E)** Scatter plot of z scores for each gRNA in CRISPR-ko screens with mCherry and BATF3 with enriched gRNAs (P_adj_ < 0.01) colored blue, red, or dark purple. **(F)** Average percentage IL7R+ (left) and relative IL7R MFI (right) in CD8+ T cells with mCherry or BATF3 across gRNAs. Relative IL7R MFI was calculated by dividing the IL7R MFI of each targeting gRNA by the IL7R MFI of the non-targeting gRNA for each donor within the treatment group (n = 3 donors, error bars represent SEM). **(G)** Representative histograms of IL7R expression in CD8+ T cells with BATF3 overexpression in combination with JUNB or IRF4 gene knockouts. **(H)** Individual and combined effects of ZNF217 knockout and BATF3 overexpression on the percentage of IL7R+ cells (n =3 donors, error bars represent SEM). A one-way, paired ANOVA test with Tukey’s post hoc test was used to compare the mean percentage of IL7R+ cells between groups.

As expected, multiple IL7R gRNAs were the most enriched gRNAs in the IL7R low population across both screens (Figure 6B). Notably, BATF3 gRNAs only emerged in the screen with BATF3 OE as BATF3 is lowly expressed at baseline (Figure 6B). BATF3 gRNAs indiscriminately target endogenous and exogenous BATF3, indicating that knocking out exogenous BATF3 nullified its effects. Many DNMT1 and FOXO1 gRNAs were strongly enriched in the IL7R low population, corroborating findings from our CRISPRi cell-sorting screens and subsequent scRNA-seq characterization of DNMT1 and FOXO1 gene silencing. Additionally, we recovered multiple gRNA hits for many genes with all four gRNAs emerging for several genes (e.g FOXO1, FOXP1, and RUNX3) (Figure S21A). The baseline expression of target gene hits was significantly higher than that of non-hit genes, as knockout screens can only capture the effects of expressed genes (Figure S21B).

We then compared gRNA and gene-level enrichment between the CRISPRko screens with or without BATF3 OE (Figure 6C-D). This enabled us to classify genes that regulate IL7R expression in a BATF3-independent or BATF3-dependent manner. For example, FOXO1 and DNMT1 were among the strongest gene hits in the IL7R low population across both screens, indicating BATF3-independent effects. Because BATF3 OE increased the dynamic range of IL7R expression, we were also able to capture unique genes enriched in IL7R low population in the CRISPRko screen with BATF3-OE. These genes represent potential co-factors or downstream actuators of BATF3-mediated effects. Given the known interaction between AP-1 and IRF TFs, we were particularly interested in members of these families that were exclusively enriched in the IL7R low population with BATF3 OE. BATF3, JUNB, and IRF4 were the top genes meeting these criteria, suggesting that BATF3 interacts with JUNB and IRF4 to mediate transcriptional control in CD8+ T cells (Figure 6C and S21C-D).

Both screens also revealed candidate gene targets for further improving ACT as ablating these TFs led to higher levels of IL7R expression (Figure 6C). The most enriched genes in the TF-KO only screen included ZNF217, RUNX3, FOXP1, GATA3, GFI1, AHR, ETS1, ZNF626, and FOXP3. Because BATF3 OE induces IL7R expression, it was more challenging to detect genes enriched in the IL7R high population in the screen with BATF3 OE. In addition, we speculated that some TFs whose effects were lost with BATF3 OE might be downstream targets of BATF3. Indeed, our RNA-seq analysis (Figure 4, S12) revealed that several of the top TFs including FOXP1, ETS1, and FOXP3 were all downregulated by BATF3 OE. This indicates that further reducing the expression levels of these TFs did not affect IL7R expression.

Interestingly, there were three overlapping hits in the IL7R high population between screens: ZNF217, GATA3, and AHR, suggesting that knocking out these genes increased IL7R expression individually and in combination with BATF3 OE. ZNF217 was the top hit across both screens and has not previously been characterized in the context of T cell biology. GATA3 has been shown to promote CD8+ T cell dysfunction with features reminiscent of a regulatory T cell phenotype and targeted deletion of GATA3 improves tumor control^59^. Moreover, both GATA3 and AHR can activate FOXP3 expression in regulatory T cells, providing further evidence of a link between T cell dysfunction and T cell regulatory activity^60–62^.

We individually validated the effects of knocking out IL7R, BATF3, JUNB, IRF4, ZNF217, and GATA3 with and without BATF3 OE (Figure 6E-G). Consistent with previous findings, BATF3 increased IL7R expression by >40% in control CD8+ T cells (∼33% to 77% IL7R+) with a non-targeting gRNA (Figure 6E). Ablating BATF3 led to partial restoration back to control IL7R levels, presumably due to incomplete nuclease activity across ectopic lentiviral copies of BATF3 in all cells. The effects of BATF3 OE were profoundly negated with either JUNB and IRF4 knockouts with both reducing IL7R expression similarly (∼30% decrease for JUNB and ∼28% decrease for IRF4) (Figure 6E-F). Conversely, genetic disruption of GATA3 and ZNF217 increased baseline IL7R levels by ∼10% and ∼18%, respectively (Figure 6E). Combined with BATF3 OE, the fraction of IL7R+ cells did not increase with GATA3 deletion, however the relative fluorescent intensity increased by 36%. Finally, the combined effect of BATF3 OE and ZNF217 knockout led to a significant proportion of IL7R+ T cells (>84%) (Figure 6G and S21E).

## Discussion

In this study, we developed and characterized compact and efficient dSaCas9-based epigenome editors to systematically map transcriptional and epigenetic regulators of primary human CD8+ T cell state through complementary loss-of-function and gain-of-function CRISPRi/a screens. Although we assayed the effects of 120 genes in our CRISPRi/a screens, this technology could readily be scaled to profile all catalogued human genes for their coordination of complex T cell phenotypes. Nevertheless, our CRISPRi/a screens recovered many known and novel regulators of CD8+ T cell state with a striking convergence on BATF3. A prominent effect of BATF3 overexpression was activation of *IL7R*, which encodes for the IL-7 receptor. A primary reason for lymphodepleting regimens before CAR T cell infusion in clinical protocols is to maximize the availability of homeostatic cytokines (IL-2, IL-7, and IL-15) by eliminating competing immune cells. Increased IL7R expression on engineered T cells therefore might increase their sensitivity to IL-7 signaling and enable lower doses of conditioning lymphodepletion agents, which increase the risk of infection and have other associated toxicities.

BATF3 overexpression markedly enhanced the ability of CD8+ T cells to control tumor growth in vitro and in vivo. The compact size of BATF3 could seamlessly integrate into current manufacturing processes of FDA-approved ACTs, which all use lentivirus to deliver the CAR or TCR to donor T cells. Before translating promising gene modules such as BATF3 overexpression into the clinic, it will be important to carefully assess the safety of engineered T cells. Although the progeny of a single TET2^null^ CAR T cell clone cured an advanced refractory chronic lymphocytic leukemia (CLL) patient^20^, a recent study highlighted that biallelic deletion of TET2 in combination with sustained expression of BATF3 can lead to antigen-independent clonal T cell expansion^63^. BATF3 OE alone does not induce adverse effects in T cells^64^, but the BATF-IRF axis can be oncogenic in the context of other genetic and epigenetic aberrations such as mutations, deletions, translocations, and duplications^65–69^. We did not detect increased levels of MYC or Ki-67 expression in our RNA-seq data nor did we detect elevated numbers of CD8+CAR+ T cells with BATF3 OE after 9 days of in vitro expansion and nearly three weeks of in vivo surveillance in tumor-bearing mice. Nevertheless, future work could focus on alternative delivery strategies such as transient delivery of mRNA or self-amplifying mRNA encoding for the transgene, tuning transgene expression through regulatory elements or genetic circuits, or suicide switches to control the activity of T cells in vivo.

To our knowledge, this work is the first example that combines TF overexpression with a TFome knockout screen to dissect co-factors and downstream factors and highlights the power of this approach. These screens provided insight into the mechanism by which BATF3 programs transcriptional changes. Specifically, our data combined with existing data from other cell types supports a model where BATF3 heterodimerizes with JUNB and interacts with IRF4 to drive transcriptional programs in CD8+ T cells. The dynamic and combinatorial interactions between AP-1 TFs have repeatedly been shown to control biological processes that dictate T cell state and function and are promising therapeutic candidates for ACT. This investigation also identified novel factors, such as ZNF217, which have not previously been associated with controlling T cell state or AP-1 gene regulation, which will be worthy targets of additional study. Overall, this work expands the toolkit of epigenome editors and our understanding of regulators of CD8+ T cell state and function. This catalogue of genes could serve as a basis for engineering the next generation of cancer immunotherapies.

## Methods

### Plasmids

All plasmids used were cloned using Gibson assembly (NEB). The all-in-one HER2 CAR constructs used for in vivo tumor control studies were cloned by digesting an empty lentiviral vector for constitutive gene expression (Addgene 79121) with MluI and amplifying the HER2-CAR^70^ and 2A-GFP or 2A-BATF3 (gblock, IDT) fragments with appropriate overhangs for Gibson assembly. The following plasmids were deposited to Addgene: pLV hU6-gRNA hUbC-dSaCas9-KRAB-T2A-Thy1.1 (Addgene 194278) and pLV hU6-gRNA hUbC-VP64-dSaCas9-VP64-T2A-Thy1.1 (Addgene 194279).

### Cell Lines

HEK293Ts and SKBR3s were maintained in DMEM GlutaMAX supplemented with 10% fetal bovine serum (FBS), 1 mM sodium pyruvate, 1x MEM non-essential amino acids (NEAA), 10 mM HEPES, 100 U mL^-1^ of penicillin, and 100 μg mL^-1^ streptomycin. Jurkats lines were maintained in RPMI supplemented with 10% FBS, 100 U mL^-1^ of penicillin, and 100 μg mL^-1^ streptomycin. HCC1954s were maintained in DMEM/F12 supplemented with 10% FBS, 100 U mL^-1^ of penicillin, and 100 μg mL^-1^ streptomycin.

### Isolation and Culture of Primary Human T Cells

Human CD8+ T cells were obtained from either pooled PBMC donors (ZenBio) using negative selection human CD8 isolation kits (StemCell Technologies) or directly from vials containing isolated CD8+ T cells from individual donors (StemCell Technologies). For all technology development experiments, T cells were cultured in Advanced RPMI (Thermo Fisher) supplemented with 10% FBS, 100 U mL^-1^ of penicillin and 100 μg mL^-1^ streptomycin. For all T cell reprogramming experiments, T cells were cultured in PRIME-XV T cell Expansion XSFM (FujiFilm) supplemented with 5% human platelet lysate (Compass Biomed), 100 U mL^-1^ of penicillin and 100 μg mL^-1^ streptomycin. All media was supplemented with 100 U mL^-1^ human IL-2 (Peprotech). T cells were activated with a 3:1 ratio of CD3/CD28 dynabeads to T cells and split or expanded every 2 days to maintain T cells at a concentration of 1-2 x 10^6^ per mL unless otherwise indicated.

### Lentivirus Generation and Transduction of Primary Human T Cells

For all technology development experiments, lentivirus was produced as previously described^34^. For all T cell reprogramming experiments, a recently optimized protocol for high-titer lentivirus was used^25^. Briefly, 1.2 x 10^6^ or 7 x 10^6^ HEK293T cells were plated in a 6 well plate or 10 cm dish in the afternoon with 2 mL or 12 mL of complete opti-MEM (Opti-MEM™ I Reduced Serum Medium supplemented with 1x Glutamax, 5% FBS, 1 mM Sodium Pyruvate, and 1x MEM Non-Essential Amino Acids). The next morning, HEK293T cells were transfected with 0.5 μg pMD2.G, 1.5 μg psPAX2, and 0.5 μg transgene for 6 well transfections or 3.25 μg pMD2.G, 9.75 μg psPAX2, and 4.3 μg transgene for 10 cm dishes using Lipofectamine 3000. Media was exchanged 6 hours after transfection and lentiviral supernatant was collected and pooled at 24 hours and 48 hours after transfection. Lentiviral supernatant was centrifuged at 600xg for 10 min to remove cellular debris and concentrated to 50-100x the initial concentration using Lenti-X Concentrator (Takara Bio). T cells were transduced at 5-10% v/v of concentrated lentivirus at 24 hours post-activation. For dual transduction experiments, T cells were serially transduced at 24 hours and 48 hours post activation.

### Design of CD2, B2M, and IL2RA gRNA Libraries

Saturation CD2 and B2M CRISPRi gRNA libraries were designed to tile a 1,050 bp window (-400 bp to 650 bp) around the TSS of each target gene using CRISPick^32^. The IL2RA CRISPRa gRNA library was designed to tile a 5kb bp window (-4,000 bp to 1000 bp) around the TSS of IL2RA using ChopChop^71^. Any gRNA that aligned to another genomic site with fewer than four mismatches was removed from the library. Each gRNA library was designed to target dSaCas9’s relaxed PAM variant: 5’-NNGRRN-3’. Non-targeting gRNAs were generated for each library to match the nucleotide composition of the targeting gRNAs. CD2, B2M, and Il2RA gRNA libraries can be found in Supplementary Table 1.

### gRNA Library Cloning

Oligonucleotide gRNA pools containing variable protospacer sequences and constant regions for PCR amplification were synthesized by Twist Bioscience. 2-4 ng of each oligonucleotide pool was input into a 7-cycle PCR with 2x Q5 mastermix and 10 μM of each amplification primer with the following cycling conditions: 98C for 10s, 65C for 30s, and 72C for 15s. The gRNA amplicon was gel extracted and then PCR purified. The purified gRNA amplicon was input into a 20 μL Gibson reaction at a 5:1 insert to backbone molar ratio with 200 ng of either all-in-one CRISPRi or CRISPRa backbones digested with Esp3I, dephosphorylated using QuickCIP, and 1x SPRI-selected. The Gibson reactions were ethanol precipitated overnight and transformed into Lucigen’s Endura ElectroCompetent Cells. Cloned gRNA libraries were purified for lentivirus production by midi-prepping 100 mL of bacterial culture.

### CD2 and B2M CRISPRi Screens in Primary Human T Cells

CD8+ T cells from pooled PBMC donors were transduced with all-in-one lentivirus encoding for dSaCas9-KRAB-2A-GFP and either CD2 (n = 2 replicates) or B2M (n = 3 replicates) gRNA libraries. Cells were expanded for 9 days and then stained for the target gene (CD2 or B2M). Transduced GFP+ T cell in the lower and upper 10% tails of target gene expression were sorted for subsequent gRNA library construction and sequencing. All replicates were maintained and sorted at a minimum of 350x coverage.

### Construction of CRISPRa Jurkat Lines and IL2RA CRISPRa Screens in Jurkats

Polyclonal dSaCas9^VP64^ and ^VP64^dSaCas9^VP64^ Jurkat cell lines were generated by transducing 2 x 10^6^ Jurkats with 2% v/v of 50x lentivirus encoding for either dSaCas9^VP64^-2A-PuroR or ^VP64^dSaCas9^VP64^-2A-PuroR. Cells were selected for five days (days 3-7 post-transduction) using 0.5ug/mL of puromycin. After puromycin selection, 1 x 10^6^ dSaCas9^VP64^ and ^VP64^dSaCas9^VP64^ Jurkat cells were plated and transduced in triplicate with the IL2RA gRNA library lentivirus at a multiplicity of infection (MOI = 0.4). Cells were expanded for 10 days, selected for Thy1.1 using a CD90.1 Positive Selection Kit (StemCell Technologies), and then stained for Thy1.1 and IL2RA. Transduced Thy1.1+ Jurkats in the lower and upper 10% tails of IL2RA expression were sorted for subsequent gRNA library construction and sequencing. All replicates were maintained and sorted at a minimum of 500x coverage.

### TF and Epi-Modifier CRISPRi/a gRNA Library Construction

Genes were selected based on motif enrichment in differentially accessible chromatin across T cell subsets^4, 36, 37^ and a unified atlas of over 300 ATAC-seq and RNA-seq experiments from 12 studies of CD8 T cells in cancer and chronic infection^38^. The following transcriptional and epigenetic regulators: BACH2, TOX, TOX2, PRDM1, KLF2, BMI1, DNMT1, DNMT3A, DNMT3B, TET1, and TET2 were manually added to the gene list. The complete 121 member gene list can be found in Supplementary Table 2. The TSS for each gene was extracted using CRISPick and 1,000 bp windows were constructed around each TSS (-500 to +500 bp). After establishing an SaCas9 gRNA database with the strict PAM variant (NNGRRT) using guideScan^72^, the genomic windows were input into the guidescan_guidequery function to generate the gRNA library. Any gRNA that aligned to another genomic site with fewer than four mismatches was removed from the library. The final gRNA library contained at least seven gRNAs targeting 120/121 target gene (there were no PBX2-targeting gRNAs) with an average of 16 gRNAs per gene. 120 non-targeting gRNAs were included in the library for a total of 2,099 gRNAs (Supplementary Table 2).

### TF and Epi-Modifier CRISPRi/a gRNA Screens

CD8+CCR7+ T cells were sorted and transduced with either CRISPRi (n = 2 donors) or CRISPRa (n = 3 donors) TF + epi-modifier gRNA libs. Cells were expanded for 10 days and then stained for Thy1.1 (a marker to identify transduced cells) and CCR7 (a marker associated with T cell state). Transduced Thy1.1+ T cells in the lower and upper 10% tails of CCR7 expression were sorted for subsequent gRNA library construction and sequencing. All replicates were maintained and sorted at a minimum of 300x coverage.

### Genomic DNA Isolation, gRNA PCR, and Sequencing gRNA Libraries

Genomic DNA was isolated from sorted cells using Qiagen’s DNeasy Blood and Tissue Kit. All genomic DNA was split across 100 μL PCR reactions with Q5 2X Master Mix, up to 1 μg of genomic DNA per reaction, and forward and reverse primers. After initial amplicon denaturation at 98C for 30s, gRNA libraries were amplified through 25 PCR cycles at 98C for 10s, 60C for 30s, and 72C for 20s, followed by a final extension at 72C for 20s. PCRs were pooled together for each sample and purified using double-sided SPRI selection at 0.6x and 1.8x to remove gDNA and primer dimer. Libraries were run on a High Sensitivity D1000 tape (Agilent) to confirm the expected amplicon size and quantified using Qubit’s dsDNA High Sensitivity assay. Libraries were individually diluted to 2 nM, pooled together at equal volumes, and sequenced using Illumina’s MiSeq Reagent Kit v2 (50 cycles) according to manufacturer’s recommendations. Read 1 was 22 cycles to sequence the 21 bp protospacers and index read 1 was 6 cycles to sequence the sample barcodes. Primers used in this study can be found in Supplementary Table 5.

### Processing gRNA Sequencing and Enrichment Analysis for FACS-based Screens

FASTQ files were aligned to custom indexes for each gRNA library (generated from the bowtie2-build function) using Bowtie 2^73^. Counts for each gRNA were extracted and used for further analysis. All enrichment analysis was done with R. Individual gRNA enrichment was determined using the DESeq2^74^ package to compare gRNA abundance between high and low conditions for each screen. gRNAs were selected as hits if they met a specific statistical significance threshold (defined in figured legends). DESeq2 results for each cell sorting based screen in this study can be found in Supplementary Tables 1 and 2.

### Individual gRNA Validation Using Flow Cytometry

For CD2 and B2M gRNA validations, CD8 T cells were transduced in triplicate with each individual gRNA and followed the same timeline as the CRISPRi screens. On day 9, cells were stained with either a CD2 or B2M antibody and measured using flow cytometry. For IL2RA gRNA validations, dSaCas9^VP64^ and ^VP64^dSaCas9^VP64^ Jurkat lines were transduced with each gRNA hit and followed the same timeline as the CRISPRa screen. On day 9, cells were stained with a IL2RA antibody and measured using flow cytometry. The percentage of cells expressing the target gene or the mean fluorescence intensity (MFI) of the target gene were reported for flow cytometry data.

### Flow Cytometry and Surface Marker Staining

An SH800 FACS Cell Sorter (Sony Biotechnology) was used for cell sorting and analysis unless otherwise indicated. For antibody staining of all surface markers except CCR7, cells were harvested, spun down at 300xg for 5 min, resuspended in flow buffer (1x PBS, 2 mM EDTA, 0.5% BSA) with the appropriate antibody dilutions and incubated for 30 min at 4C on a rocker. Antibody staining of CCR7 was carried out for 30 min at 37C. Cells were then washed with 1 mL of flow buffer, spun down at 300xg for 5 min, and resuspended in flow buffer for cell sorting or analysis. Antibody details can be found in Supplementary Table 5. FMO controls were used to set appropriate gates for all flow panels.

### Quantitative RT-qPCR

mRNA was isolated from transduced primary human CD8+ T cells or Jurkats using Norgen’s Total RNA Purification Plus Kit. Reverse transcription was carried out by inputting an equal mass of mRNA for each sample into a 10 μL SuperScript Vilo cDNA Synthesis reaction. 2.0 μL of cDNA was used per PCR reaction with Perfecta SYBR Green Fastmix (Quanta BioSciences, 95072) using the CFX96 Real-Time PCR Detection System (Bio-Rad). All primers were designed to be highly specific using NCBI’s primer blast tool and amplicon products were verified by melt curve analysis. All qRT-qPCR are presented as log_2_ fold change in RNA normalized to GAPDH expression unless otherwise indicated. Primers used in this study can be found in Supplementary Table 5.

### Characterization of TF Hits Using scRNA-seq

All 32 gRNA hits (as defined by a P_adj_ < 0.05) from the CRISPRi/a screens and 8 non-targeting gRNAs were selected for scRNA-seq characterization. This 40-gRNA library (Supplementary Table 3) was cloned into the all-in-one CRISPRi and CRISPRa lentiviral plasmids. The experimental timeline for the scRNA-seq screens was identical to the cell sorting-based screens. CD8+CCR7+ T cells from three donors were transduced with CRISPRi and CRISPRa mini-TF gRNA libraries. T cells were expanded for 10 days and then stained and sorted for Thy1.1+ cells. Sorted cells were loaded into the Chromium X for a targeted recovery of 2 x 10^4^ cells per donor and treatment according to the Single Cell 5’-High-Throughput (HT) Reagent Kit v2 protocol (10x Genomics). SaCas9 gRNA sequences were captured by spiking in 2 μM of a custom primer into the reverse transcription master mix, as previously done for SpCas9 gRNA capture^44^. The custom primer was designed to bind to the constant region of SaCas9’s gRNA scaffold. 5’-Gene Expression (GEX) and gRNA libraries were separated using double-sided SPRI selection in the initial cDNA clean up step. 5’-GEX libraries were constructed according to manufacturer’s protocol. gRNA libraries were constructed using two sequential PCRs (PCR 1: 10 cycles, PCR 2: 25 cycles). The PCR 1 product was purified using double-sided SPRI selection at 0.6x and 2x. 20% of the purified PCR 1 product was input into PCR 2. The PCR2 product was purified using double-sided SPRI selection at 0.6x and 1x. All libraries were run on a High Sensitivity D1000 tape to measure the average amplicon size and quantified using Qubit’s dsDNA High Sensitivity assay. Libraries were individually diluted to 20 nM, pooled together at desired ratios, and sequenced on an Illumina NovaSeq S4 Full Flow Cell (200 cycles) with the following read allocation: Read 1 = 26, i7 index = 10, Read 2 = 90. All oligos used in this study can be found in Supplementary Table 5.

### Processing and Analyzing scRNA-seq

CellRanger v6.0.1 was used to process, demultiplex, and generate UMI counts for each transcript and gRNA per cell barcode. UMI counts tables were extracted and used for subsequent analyses in R using the Seurat^75^ v4.1.0 package. Low quality cells with < 200 detected genes, > 20% mitochondrial reads, or < 5% ribosomal reads were discarded. DoubletFinder^76^ was used to identify and remove predicted doublets. All remaining high-quality cells across donors for each treatment (CRISPRi or CRISPRa) were aggregated for further analyses. gRNAs were assigned to cells if they met the threshold (gRNA UMI > 4). Cells were then grouped based on gRNA identity. For differential gene expression analysis, we compared the transcriptomic profiles of cells sharing a gRNA to cells with only non-targeting gRNAs using Seurat’s FindMarkers function to test for differentially expressed genes (DEGs) with the hurdle model implemented in MAST. All significant gRNA-to-gene links can be found in Supplementary Table 3. Upregulated DEGs were input into EnrichR’s GO Biological Process 2021 database^77^ for functional annotation.

### RNA-sequencing with BATF3 Overexpression

CD8+ T cells were transduced with lentivirus encoding for BATF3-2A-GFP or GFP and expanded for 10 days. On day 10, 4 x 10^5^ GFP+ T cells were sorted for subsequent RNA isolation using Norgen’s Total RNA Purification Plus Kit. RNA was submitted to Azenta (formerly Genewiz) for standard RNA-seq with polyA selection. Reads were first trimmed using Trimmomatic^78^ v0.32 to remove adapters and then aligned to GRCh38 using STAR v2.4.1a aligner. Gene counts were obtained with featureCounts^79^ from the subread package (version 1.4.6-p4) using the comprehensive gene annotation in Gencode v22. Differential expression analysis was determined with DESeq2^74^ where gene counts are fitted into a negative binomial generalized linear model (GLM) and a Wald test determines significant DEGs (P_adj_ < 0.01). All DEGs can be found in Supplementary Table 4. Upregulated and downregulated DEGs were input into EnrichR’s GO Biological Processes 2021 database for functional annotation.

### Single cell RNA-seq analysis of CD19 CAR T cell infusion product for responders and non-responders

scRNA-seq data of the infused CD19 CAR T cell products from patients treated with tisagenlecleucel^49^ were downloaded from GEO:GSE197268. Patient data in MarketMatrix format were classified as responders (R) and non-responders (NR) and processed with Seurat^80^ 4.2.0. For each patient, cells with fewer than 20% mitochondrial UMI counts, more than 20 gene expression (GEX) UMI counts, and in the bottom 95th percentile of GEX UMI counts were selected. GEX UMI counts were log-normalized for further analysis. Individual patient data were merged (merge function in Seurat) into a combined Seurat object, preserving the group identity in the cellular barcodes. GEX UMI counts were linearly scaled and centered (ScaleData function with default parameters) before finding the most differentially expressed genes (Seurat FindVariableFeatures) using principal component analysis (PCA). Clustering was performed using the first 10 principal components to identify and select CD8+ T cells for subsequent analyses. MAST was used to identify differentially expressed genes between CD8+ T cells from responders and non-responders. All DEGs between responders and non-responders can be found in Supplementary Table 4.

### ATAC-seq

5 x 10^4^ transduced CD8+ T cells were sorted for Omni ATAC-seq as previously described^81^. Libraries were sequenced on an Illumina NextSeq 2000 with paired-end 50bp reads. Read quality was assessed with FastQC and adapters were trimmed with Trimmomatic^78^. Trimmed reads were aligned to the Hg38 reference genome using Bowtie^82^ (v1.0.0) using parameters -v 2 --best --strata -m 1. Reads mapping to the ENCODE hg38 blacklisted regions were removed using bedtools2^83^ intersect (v2.25.0). Duplicate reads were excluded using Picard MarkDuplicates (v1.130; http://broadinstitute.github.io/picard/). Count per million normalized bigWig files were generated for visualization using deeptools bamCoverage^84^ (v3.0.1). Peak calling was performed using MACS2 narrowPeak^85^ and filtered for P_adj_ ≤ 0.001. Peak calls were merged across samples to make a union-peak set. A count matrix containing the number of reads in peaks for each sample was generated using featureCounts^79^ (subread v1.4.6) and used for differential analysis in DESeq2^74^ (v.1.36). ChIPSeeker^86^ was used to annotate the genomic regions and retrieve the nearest gene around each peak.

### In Vitro Tumor Killing Assay

CD8+ T cells were transduced with lentiviruses encoding for a HER2-CAR-mCherry at 24 hours post-activation and BATF3-2A-GFP or GFP at 48 hours post-activation. After 12 days of expansion, CAR+GFP+ T cells were sorted and counted for the co-culture assay. Four hours before starting the co-culture, 2 x 10^5^ HER2+ SKBR3s were plated in a 24 well plate with cDMEM to allow the SKBR3s to adhere to the plate. After four hours, cDMEM was discarded and mCherry+GFP+ T cells in cPRIME media were added at the indicated effector to target (E:T) cell ratios. After 24 hours of co-culture, the cells were harvested by collecting the supernatant (containing T cells and dead tumor cells) and adherent cells (which were detached from the plate using trypsin). Cells were spun down at 600xg for 5 min and then stained with a fixable viability dye (FVD) and Annexin V to label dead and apoptotic cells according to manufacturer’s protocol. Stained cells were analyzed using flow cytometry. The percentage of viable tumor cells was quantified using the following strict gating strategy. First, T cells were excluded based on cell size and GFP signal. Next, a gate was set around the double negative (FVD-, Annexin V-) fraction containing viable tumor cells and cellular debris. Visualizing these events on SSC vs. FSC, a gate was set to encompass events located in the bottom left quadrant. This gate was then inverted to exclude debris from the viability calculation and moved immediately beneath the T cell exclusion gate on the gating hierarchy. Tumor viability was reported using the percentage of tumor cells in the final double negative (FVD-, Annexin V-) gate.

### CD3/CD28 and Tumor Repeat Stimulations

For repeated rounds of CD3/CD28 dynabead stimulation, CD3/CD28 beads were removed, cells were counted, replated at 1-2.5 x 10^5^ T cells, and restimulated with new CD3/CD28 beads at a 3:1 bead to cell ratio in a 24 well plate every 3 days. On day 12, cells were stained and analyzed for expression of exhaustion-associated markers using flow cytometry. For repeated rounds of tumor stimulation, 1 x 10^5^ HER2 CAR T cells were transferred to a new 24 well plate with 2 x 10^5^ SKBR3s for a 1:2 E:T ratio every 3 days. T cells were recovered without antigen stimulation for two days after the final round of tumor stimulation before ATAC-seq on day 14. For both modes of chronic stimulation, T cells were restimulated on days 3, 6, and 9.

### Mice

All experiments involving animals were conducted with strict adherence to the guidelines for the care and use of laboratory animals of the National Institutes of Health (NIH). All experiments were approved by the Institutional Animal Care and Use Committee (IACUC) at Duke University (protocol number A130-22-07). 6–8-week-old female immunodeficient NOD/SCID gamma (NSG) mice were obtained from Jackson Laboratory and then housed and handled in pathogen-free conditions.

### In Vivo Tumor Model

2.5 x 10^6^ HCC1954 cells were implanted orthotopically into the mammary fat pad of NSG mice in 100 μL 50:50 (v:v) PBS:Matrigel. T cells were expanded for 9-11 days post-transduction before treatment. Transduction rates were measured on the day of treatment using flow cytometry. For all in vivo experiments, transduction rates exceeded 70% for both HER2-CAR-2A-GFP and HER2-CAR-2A-BATF3 constructs. T cells were resuspended at 50 x 10^6^ CAR+ cells mL^-1^ in 1x PBS and serially diluted to the appropriate cell concentrations for 200 μL injections of either 10 x 10^6^, 2 x 10^6^, 5 x 10^5^, 2.5 x 10^5^, or 1 x 10^5^ HER2 CAR+ T cells. 20-21 days after tumor implantation, and immediately prior to CAR T cell injections, mice were randomized into groups and tumors measured. Tumor volumes were calculated based on caliper measurements using the formula volume: = ½ (Length × Width^2^). CAR T cells were injected intravenously by tail vein injection. Tumors were measured every 4-6 days.

### Flow cytometry analysis of input and tumor infiltrating CAR T cells

Mice bearing HCC1954 tumors were euthanized at days 3 and 19 post CAR T cell delivery under deep isoflurane anesthesia via exsanguination, from which blood was collected. Blood was processed via RBC lysis buffer (Sigma) treatment followed by washing in PBS. Tumors were resected, minced, and incubated in RPMI-1640 medium (Gibco) for 45 minutes in 100µg/ml Liberase-TM (Sigma-Aldrich) and 10µg/ml DNAse I (Roche). Single cell suspensions for blood and tumor were filtered through a 70mm cell strainer (Olympus Plastics), washed in PBS (Gibco), stained with Zombie NIR (1:250, Biolegend), washed in FACs buffer [2% FBS (Sigma) + PBS], and treated with 1:50 Mouse Tru-stain Fc block (Biolegend). Cells were then stained for cell surface markers followed by intracellular staining using the Transcription Factor Staining Buffer Set (Invitrogen) per manufacturer’s instructions. Fluorophore conjugated antibodies against the following antigens were used for input and day 3 cells (All Biolegend unless otherwise noted): panel 1: myc-APC (Cell Signaling Technologies), CD3-BUV737 and CD8-BUV395 (BD Biosciences), TIGIT-BV605, LAG3-BV786, CD127-PERCPCy5.5, PD1-BV711, Tim3-PECy5, GranzymeB-PECy7, TCF1-BV421, Ki67-BV510, and IFN-γ; panel 2: myc-APC, CD3-BUV737, CD8-BUV395, CD39-PECF594, CD56-BV605, CD45RO-BV786, CD45RA-PEcy5, CD28-PECy7, CCR7-BV711, CD62L-BV510, CTLA4-BV421, Tbet-PERCPCy5.5, EOMEs-PE. For day 19 post CAR T cell delivery analyses anti-human CD45-FITC (Biolegend) staining was added to the above panels to increase sensitivity of CAR T cell detection, as we anticipated reduction in numbers, and the following additional panel was added against the following antigens: CD45-FITC, myc-APC, CD3-BUV737, CD8-BUV395, LAG3-BV786, TIM3-PECy5, CXCR3-BV711, CD4-BV510, TNF-BV605, ID2-PECy7, GATA3-BV421, IRF4-PERCPCy5.5, ID3-PE. All data were collected on a Fortessa X 20 (Duke Cancer Institute Flow Cytometry Core) and analyzed using Flow Jo V10.8.1. Blood/tumor from sham infused mice and fluorescence minus one controls were used to guide gating for CAR T cells and to confirm appropriate compensation, respectively.

### TFome CRISPRko gRNA library construction

The Brunello genome wide knockout^31^ library was subset for 1,612 TFs^58^ and IL7R. 550 non-targeting gRNAs were included in the library for a total of 7,000 gRNAs (Supplementary Table S6). This gRNA library was cloned into SpCas9 gRNA lentiviral plasmids with either mCherry or BATF3.

### TFome CRISPRko screens and validations

20 x 10^6^ CD8+ T cells from two donors were activated with CD3/CD28 dynabeads at a 1:1 ratio. At 24 hours post-activation, CD8+ T cells were split evenly and transduced in parallel with TFome CRISPRko gRNA libraries with mCherry or BATF3. At 48 hours post-activation, cells were electroporated with Cas9 protein. Briefly, the cells were collected, spun down at 90xg for 10 minutes, resuspended in 100μL of Lonza P3 Primary Cell buffer with 3.2 μg Cas9 per 10^6^ cells, and electroporated with the pulse code EH115. After electroporation, warm media was immediately added to each cuvette and cells were recovered at 37C for 20 minutes before being transferred into a 6-well plate. On day 3 post transduction, cells were selected with 2 μg/mL of puromycin for 3 days. On day 9 post transduction, cells were stained for CD8, IL7R, and a viability dye. Viable CD8+ T cells in the lower and upper 10% tails of IL7R expression were sorted for subsequent gRNA library construction and sequencing. All replicates were maintained and sorted at a minimum of 75x coverage. Subsequent individual gRNA validations were scaled down to 3.5 x 10^5^ cells per electroporation in an 8-well cuvette strip, but otherwise followed the same protocol and timeline as the CRISPRko screens.

### TFome CRISPRko screen analyses

gRNA enrichment was performed using DESeq2 as explained above. Gene level enrichment was performed using the MAGeCK^87^ test module with --paired and --control sgrna parameters, pairing samples by donors and non-targeting gRNAs as control, respectively.

### Statistics

Statistical details for all experiments can be found in the figure legends. ns = not significant, * < 0.05, ** < 0.01, *** < 0.001, **** < 0.0001.

## Author Contributions

S.R.M., A.S., M.C.B., C.M.A., J.I., and C.A.G. designed experiments. S.R.M., A.S., M.C.B., C.M.A., K.S., and L.H. performed the experiments. S.R.M, A.S., and M.C.B. performed the in vivo tumor experiments. S.R.M. and A.B. analyzed the single cell RNA-sequencing data. T.E.R., S.N., S.A., and C.A.G. supervised the study. S.R.M. and C.A.G. wrote the manuscript with contributions by all authors.

## Data Availability

All data associated with this study are present in the manuscript or its Supplementary Information files. All scRNA-seq, RNA-seq, and ATAC-seq data have deposited in the Gene Expression Omnibus (GEO) under accession number: GSE218988.

## Acknowledgements

We thank all members of the Gersbach lab members for technical assistance and helpful discussions. We thank Wilson Wong for generously providing the HER2 CAR plasmid. Illustrative schematics (Figure 1A-B, 2A, 6A, and S13A) were created using BioRender. This work was supported by National Institutes of Health grants U01AI146356 (C.A.G.) UM1HG013053, UM1HG009428, and RM1HG011123 (T.E.R. and C.A.G.), National Science Foundation grants EFMA-1830957 (C.A.G.), an Allen Distinguished Investigator Award from the Paul G. Allen Frontiers Group to C.A.G, the Open Philanthropy Project, and the Duke-Coulter Translational Partnership.

## Conflict of Interest

S.R.M. and C.A.G. are named inventors on patent applications related to epigenome engineering technologies in primary human T cells. S.R.M. is a consultant for Tune Therapeutics. C.A.G. is a co-founder of Tune Therapeutics and Locus Biosciences, and is an advisor to Sarepta Therapeutics.

