## Supplementary Figures for "Orthogonal CRISPR screens to identify transcriptional and epigenetic regulators of human CD8 T cell function"

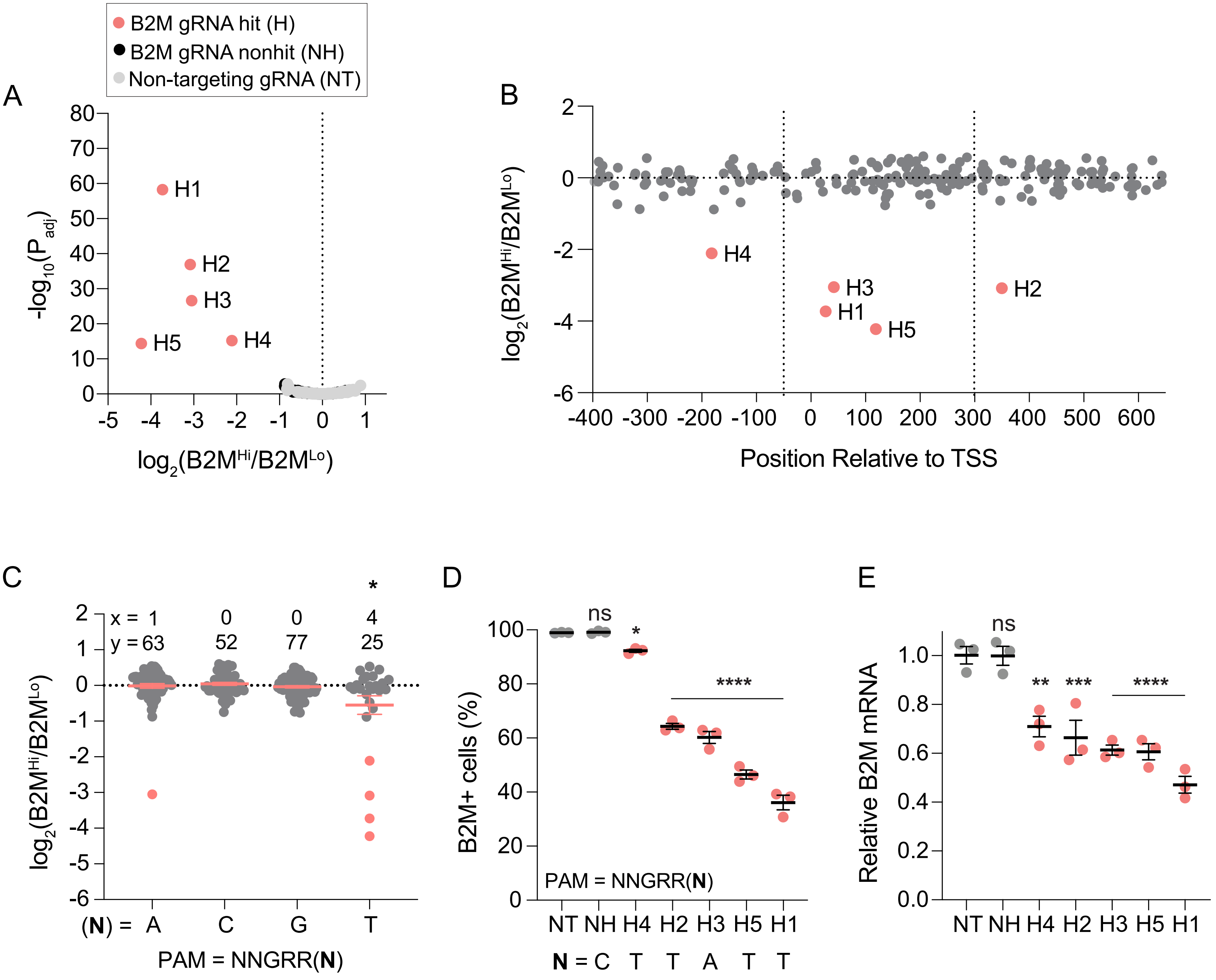
B2M gRNA hit (H)
B2M gRNA nonhit (NH)
Non-targeting gRNA (NT)
A
-log10(Padj)
log2(B2MHi/B2MLo)
H1
H2
H3
H4
H5
B
log2(B2MHi/B2MLo)
Position Relative to TSS
C
log2(B2MHi/B2MLo)
x = 1 0 0 4
y = 63 52 77 25
(N) = A C G T
PAM = NNGRR(N)
D
B2M+ cells (%)
PAM = NNGRR(N)
ns
*
****
NT NH H4 H2 H3 H5 H1
N = C T T A T T
E
Relative B2M mRNA
ns
**
***
****
NT NH H4 H2 H3 H5 H1

**Figure S1**

**Figure S1. Proof-of-principle CRISPRi B2M promoter tiling screen in primary human CD8+ T cells.**

**(A**) Volcano plot of significance ($P_{adj}$) versus fold change in gRNA abundance between B2M-high and B2M-low populations for the B2M CRISPRi screen. Red data points indicate B2M gRNA hits with a $P_{adj} < 10^{-10}$. Black data points indicate non-significant B2M gRNAs. Gray data points indicate the 250 NT gRNAs.

**(B)** B2M gRNA fold change versus gRNA position relative to the transcriptional start site (TSS). Dashed lines represent the previously defined optimal window (-50 to +300 bp of TSS) for CRISPRi.

**(C)** B2M gRNA fold change as a function of the final base pair of the PAM (5'-NNGRRN-3'). x represents the number of gRNA hits and y represents the total number of gRNAs in the library for each PAM variant. A global one-way ANOVA with Dunnett's post hoc test was used to compare the average fold change of gRNAs for each PAM variant to NNGRRT (* < 0.05 denotes that the fold change of gRNAs targeting NNGRRT PAMs was significantly different than all other PAM variants).

**(D)** Validation of B2M gRNA hits. Percentage of B2M positive cells on day 9 post-transduction plotted in rank order based on the mean gRNA activity (n = 3 replicates of CD8+ T cells from pooled PBMC donors, error bars represent SEM). A one-way ANOVA with Dunnett's post hoc test was used to compare the mean percentage of B2M positive cells for each gRNA to NT. The final base pair of the PAM for each gRNA is indicated beneath the gRNA label.

**(E)** Relative B2M mRNA expression of CD8+ cells transduced with indicated gRNA on day 9 post-transduction (n = 3, error bars represent SEM) using RT-qPCR. A one-way ANOVA with Dunnett's post hoc test was used to compare each gRNA to the NT.

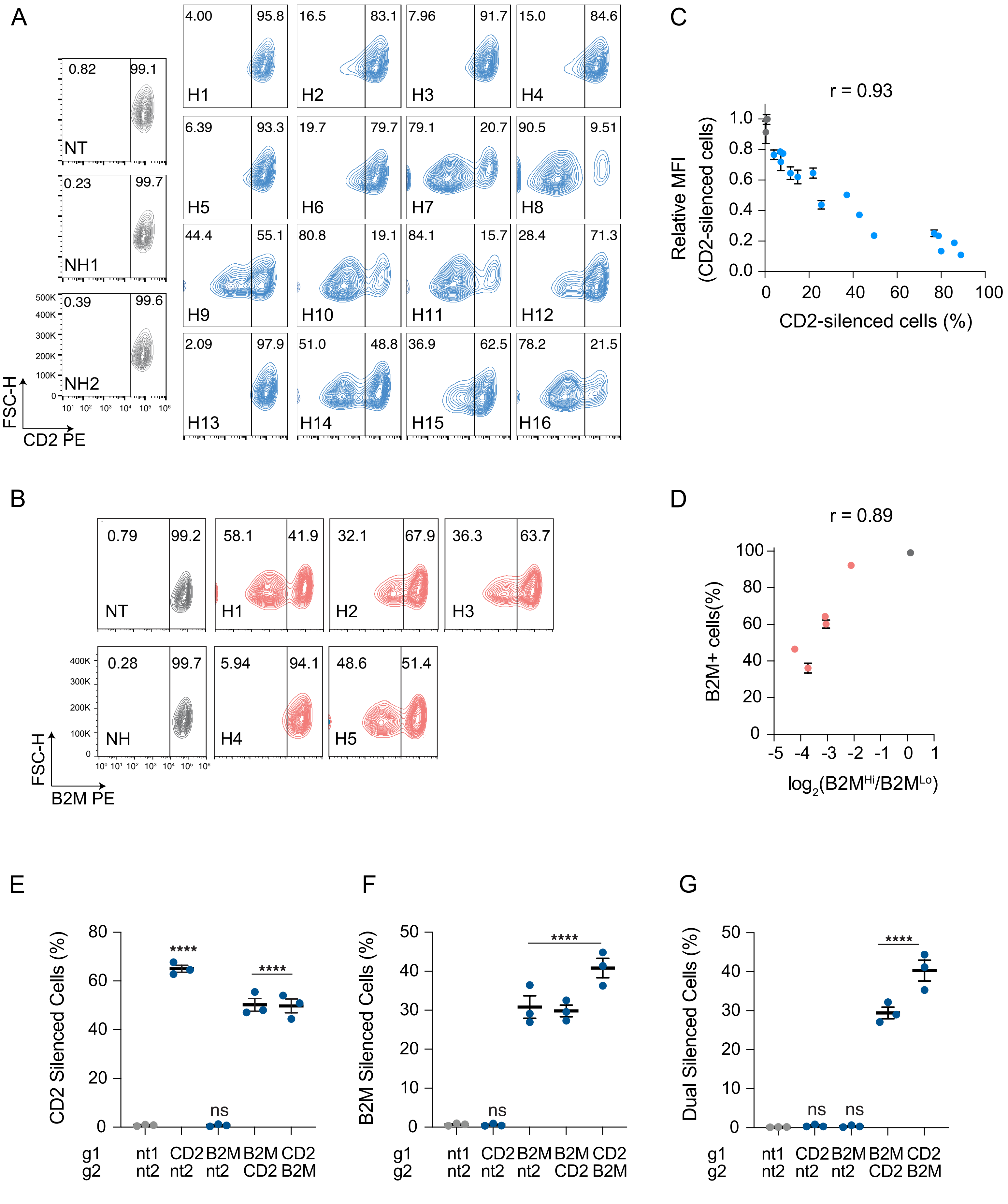
A
NT
NH1
NH2
FSC-H
CD2 PE
H1
H2
H3
H4
H5
H6
H7
H8
H9
H10
H11
H12
H13
H14
H15
H16
C
r = 0.93
Relative MFI (CD2-silenced cells)
CD2-silenced cells (%)
B
NT
NH
H1
H2
H3
H4
H5
FSC-H
B2M PE
D
r = 0.89
B2M+ cells(%)
log2(B2MHi/B2MLo)
E
CD2 Silenced Cells (%)
F
B2M Silenced Cells (%)
G
Dual Silenced Cells (%)
g1
g2
nt1 CD2 B2M B2M CD2
nt2 nt2 nt2 CD2 B2M
ns
****

### Figure S2

**Figure S2. Flow cytometry validation of CD2 and B2M gRNA hits and multiplex gene silencing.**

Representative contour plots of **(A)** CD2 and **(B)** B2M expression in CD8+ T cells across non-targeting (NT), non-hit (NH), and gRNA hits (H) measured on day 9 post transduction.

**(C)** Relationship between relative CD2 mean fluorescence intensity (MFI) of CD2 silenced cells and the percentage of CD2 silenced cells. Pearson's correlation coefficient (r) is indicated above the graph.

**(D)** Relationship between B2M gRNA activity and fold change enrichment in screen. Pearson's correlation coefficient (r) is indicated above the graph.

Average percentage of **(E)** CD2 silenced, **(F)** B2M silenced, and **(G)** dual CD2 and B2M silenced CD8+ T cells on day 10 post transduction with the indicated pairs of non-targeting, CD2, and B2M gRNAs (n = 3 replicates of CD8+ T cells from pooled PBMC donors, error bars represent SEM). g1 is driven by a human U6 promoter and g2 is driven by a mouse U6 promoter. CD2 H8 and B2M H1 gRNAs were used for multiplex gene silencing experiments.

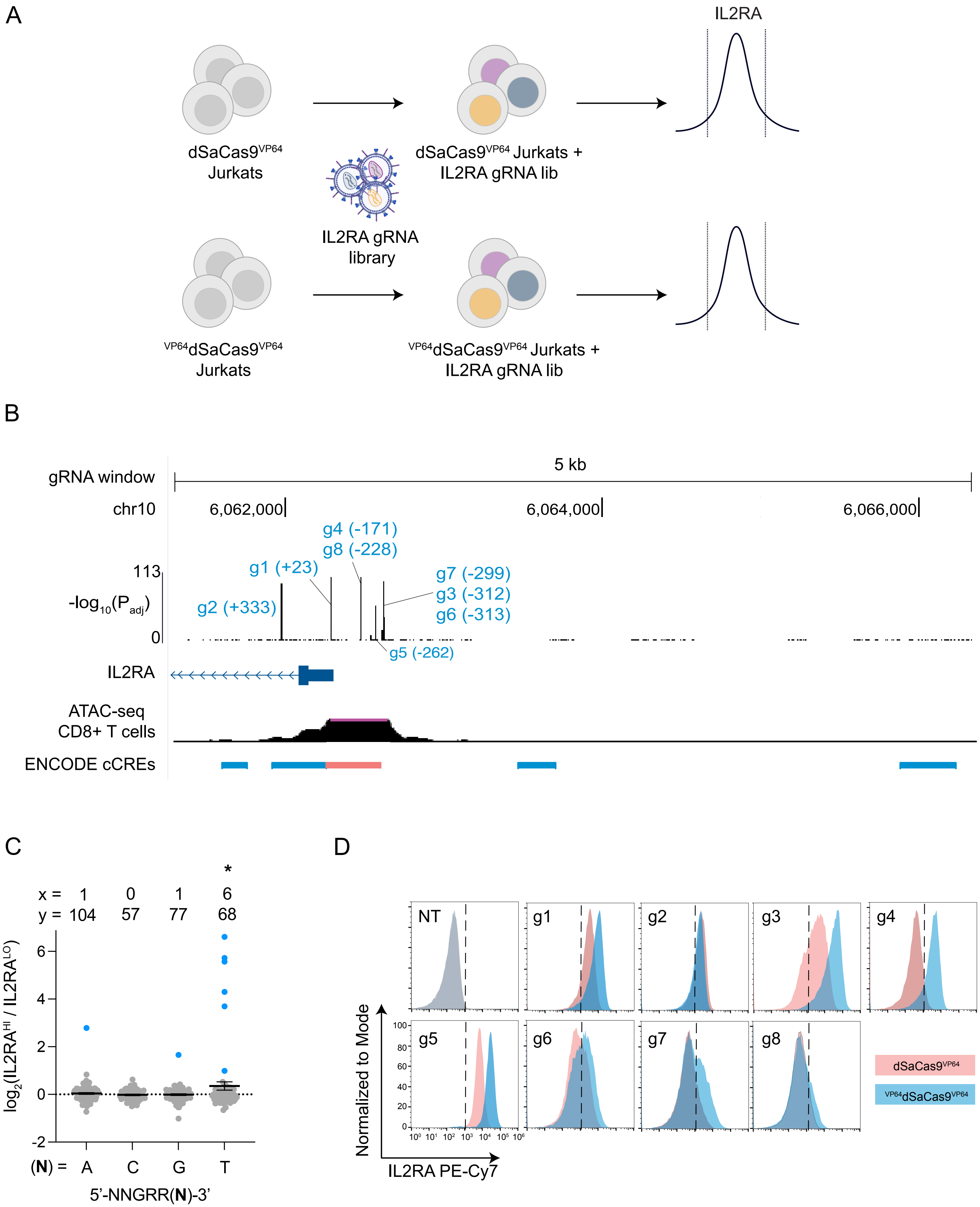

A
IL2RA
dSaCas9VP64
Jurkats
dSaCas9VP64 Jurkats +
IL2RA gRNA lib
IL2RA gRNA
library
VP64dSaCas9VP64
Jurkats
VP64dSaCas9VP64 Jurkats +
IL2RA gRNA lib
B
gRNA window
5 kb
chr10
6,062,000
6,064,000
6,066,000
113
-log10(Padj)
0
g4 (-171)
g8 (-228)
g1 (+23)
g2 (+333)
g7 (-299)
g3 (-312)
g6 (-313)
g5 (-262)
IL2RA
ATAC-seq
CD8+ T cells
ENCODE cCREs
C
*
x = 1 0 1 6
y = 104 57 77 68
log2(IL2RAHI / IL2RALO)
6
4
2
0
-2
(N) = A C G T
5'-NNGRR(N)-3'
D
NT
g1
g2
g3
g4
g5
g6
g7
g8
Normalized to Mode
100
80
60
40
20
0
IL2RA PE-Cy7
dSaCas9VP64
VP64dSaCas9VP64


**Figure S3**

**Figure S3. dSaCas9$^{VP64}$ and $^{VP64}$dSaCas9$^{VP64}$ IL2RA promoter tiling CRISPRa screens in Jurkats.**

**(A)** Schematic of dSaCas9$^{VP64}$ and $^{VP64}$dSaCas9$^{VP64}$ IL2RA promoter tiling CRISPRa screens in Jurkats.

**(B**) UCSC genome browser track of IL2RA locus with statistical significance displayed for each gRNA in $^{VP64}$dSaCas9$^{VP64}$ CRISPRa screen. gRNA hits are annotated and labeled in blue. ATAC-seq and ENCODE candidate cis regulatory elements (cCREs) tracks are overlayed for visualization of chromatin accessibility and annotations. cCREs in red are defined as promoter-like elements and cCREs in yellow are defined as enhancer-like elements.

**(C)** Fold change in gRNA abundance as a function of the final base pair of the PAM (5'-NNGRRN-3') for IL2RA gRNAs. x represents the number of gRNA hits for each PAM and y represents the total number of gRNAs for each PAM. A global one-way ANOVA with Dunnett's post hoc test was used to compare the average fold change of gRNAs for each PAM variant to NNGRRT (* < 0.05 denotes that the fold change of gRNAs targeting NNGRRT PAMs was significantly different than all other PAM variants).

**(D)** Representative overlayed histograms of IL2RA expression for dSaCas9$^{VP64}$ and $^{VP64}$dSaCas9$^{VP64}$ Jurkat lines on day 9 post-transduction across gRNAs.

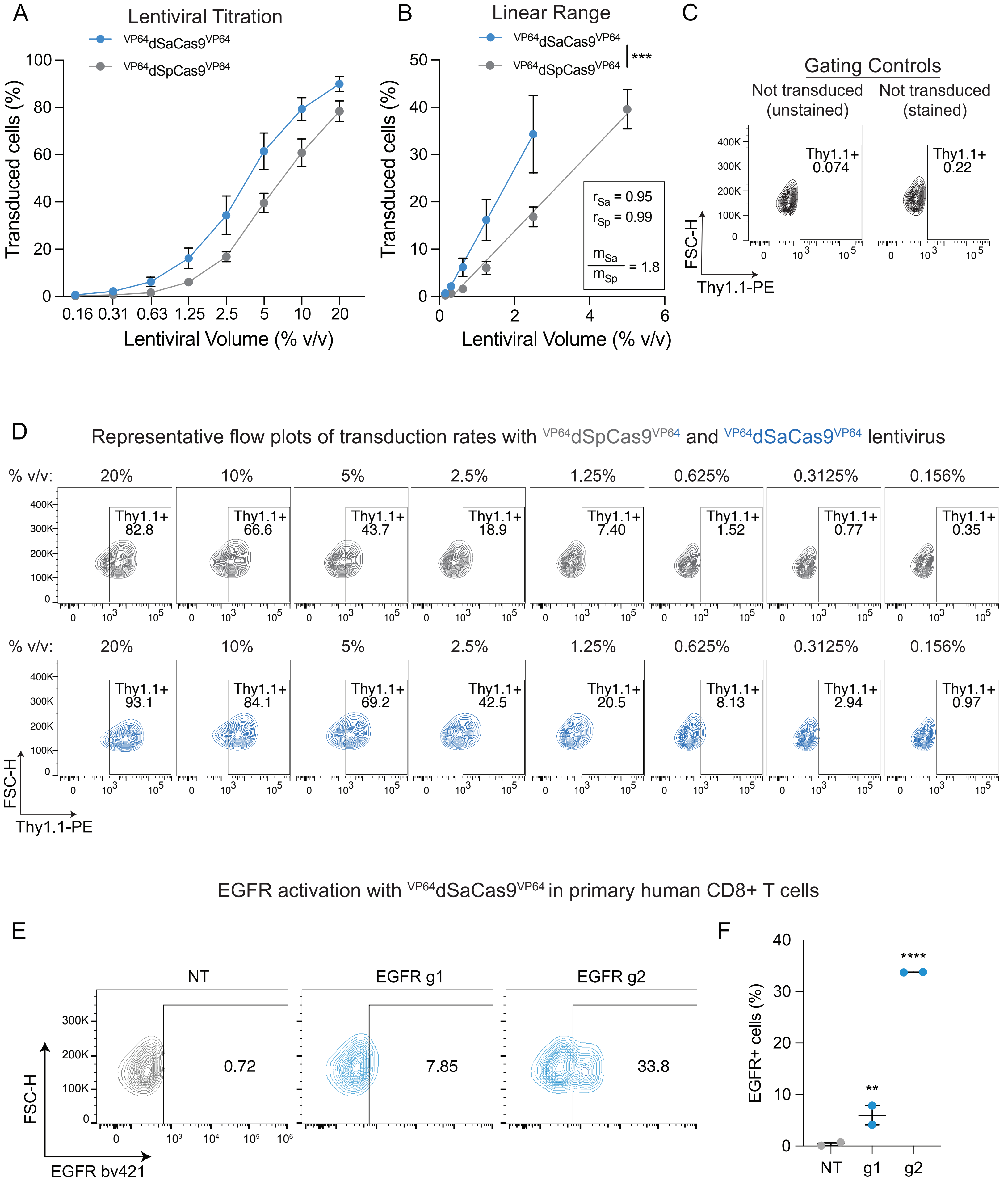

A
Lentiviral Titration
VP64dSaCas9VP64
VP64dSpCas9VP64
Transduced cells (%)
Lentiviral Volume (% v/v)
0.16 0.31 0.63 1.25 2.5 5 10 20
B
Linear Range
***
r_Sa = 0.95
r_Sp = 0.99
m_Sa / m_Sp = 1.8
C
Gating Controls
Not transduced (unstained)
Not transduced (stained)
Thy1.1+ 0.074
Thy1.1+ 0.22
FSC-H
Thy1.1-PE
D
Representative flow plots of transduction rates with VP64dSpCas9VP64 and VP64dSaCas9VP64 lentivirus
% v/v: 20% 10% 5% 2.5% 1.25% 0.625% 0.3125% 0.156%
Thy1.1+ 82.8 | 66.6 | 43.7 | 18.9 | 7.40 | 1.52 | 0.77 | 0.35
Thy1.1+ 93.1 | 84.1 | 69.2 | 42.5 | 20.5 | 8.13 | 2.94 | 0.97
EGFR activation with VP64dSaCas9VP64 in primary human CD8+ T cells
E
NT 0.72
EGFR g1 7.85
EGFR g2 33.8
EGFR bv421
F
EGFR+ cells (%)
NT g1 g2
**
****


**Figure S4**

**Figure S4. Head-to-head comparison of $^{VP64}$dSaCas9$^{VP64}$ and $^{VP64}$dSpCas9$^{VP64}$ lentiviral titers and $^{VP64}$dSaCas9$^{VP64}$ activity in primary human CD8+ T cells.**

**(A)** Transduction rate of primary human CD8+ T cells as a function of lentiviral volume for all-in-one $^{VP64}$dSaCas9$^{VP64}$ and $^{VP64}$dSpCas9$^{VP64}$ plasmids on day 9 post-transduction (n = 2 donors, error bars represent SEM).

**(B)** Linear range of transduction rate as a function of lentiviral volume. Pearson's correlation coefficient (r) and ratio of slopes were calculated using simple linear regression. (n = 2 donors, error bars represent SEM, *** < 0.001 denotes the slopes of the two lines are significantly different).

**(C)** Flow cytometry controls used to set the Thy1.1+ gate for the lentiviral titer experiment.

**(D)** Representative contour plots of Thy1.1 expression in CD8+ T cells transduced with serial titrations of $^{VP64}$dSaCas9$^{VP64}$ and $^{VP64}$dSpCas9$^{VP64}$ lentivirus.

**(E)** Representative contour plots of EGFR expression in primary human CD8+ T cells on day 8 post-transduction with all-in-one lentiviruses encoding for $^{VP64}$dSaCas9$^{VP64}$ and either a non-targeting (NT) or an EGFR gRNA.

**(F)** Summary statistics of EGFR activation (n = 2 replicates of CD8+ T cells from pooled PBMC donors, error bars represent SEM). A global one-way ANOVA with Dunnett's post hoc test was used to compare the effect of EGFR gRNAs to the NT gRNA.

A

TF + epi-modifier gRNA library

| | |
|---|---|
| Genes | 120 |
| Class | Curated TFs + epi-modifiers |
| Targets | Promoters |
| Average gRNAs per TSS | 16 |
| Non-targeting gRNAs | 120 |
| Total | 2099 |

B

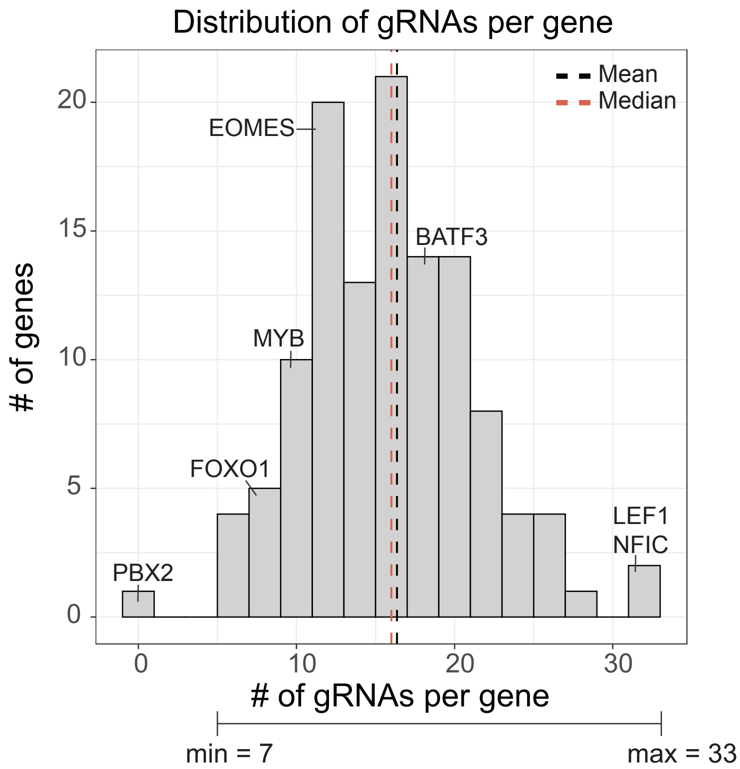


C

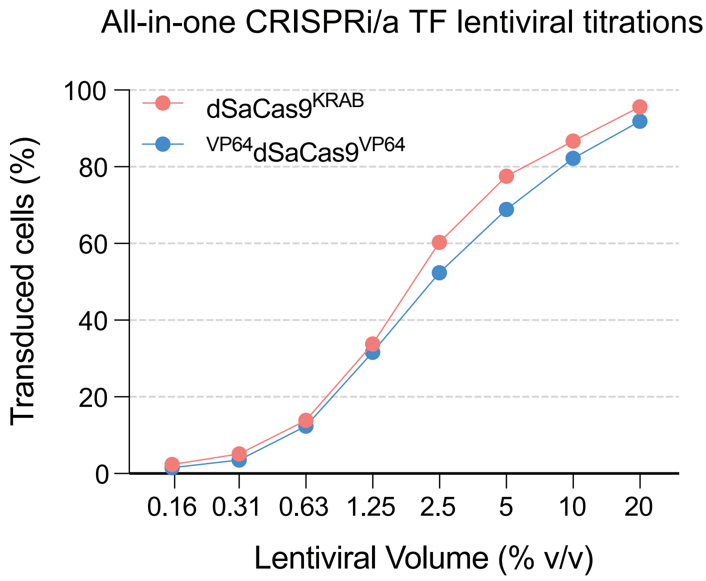


**Figure S5**

**Figure S5. TF and epigenetic modifier gRNA library design.**

**(A)** Details of gRNA library targeting curated list of TFs and epigenetic modifiers

**(B**) Histogram of gRNA representation across 121 candidate genes.

**(C)** Transduction rate of primary human CD8+ T cells as a function of lentiviral volume for all-in-one CRISPRi and CRISPRa TF gRNA plasmids on day 9 post-transduction.

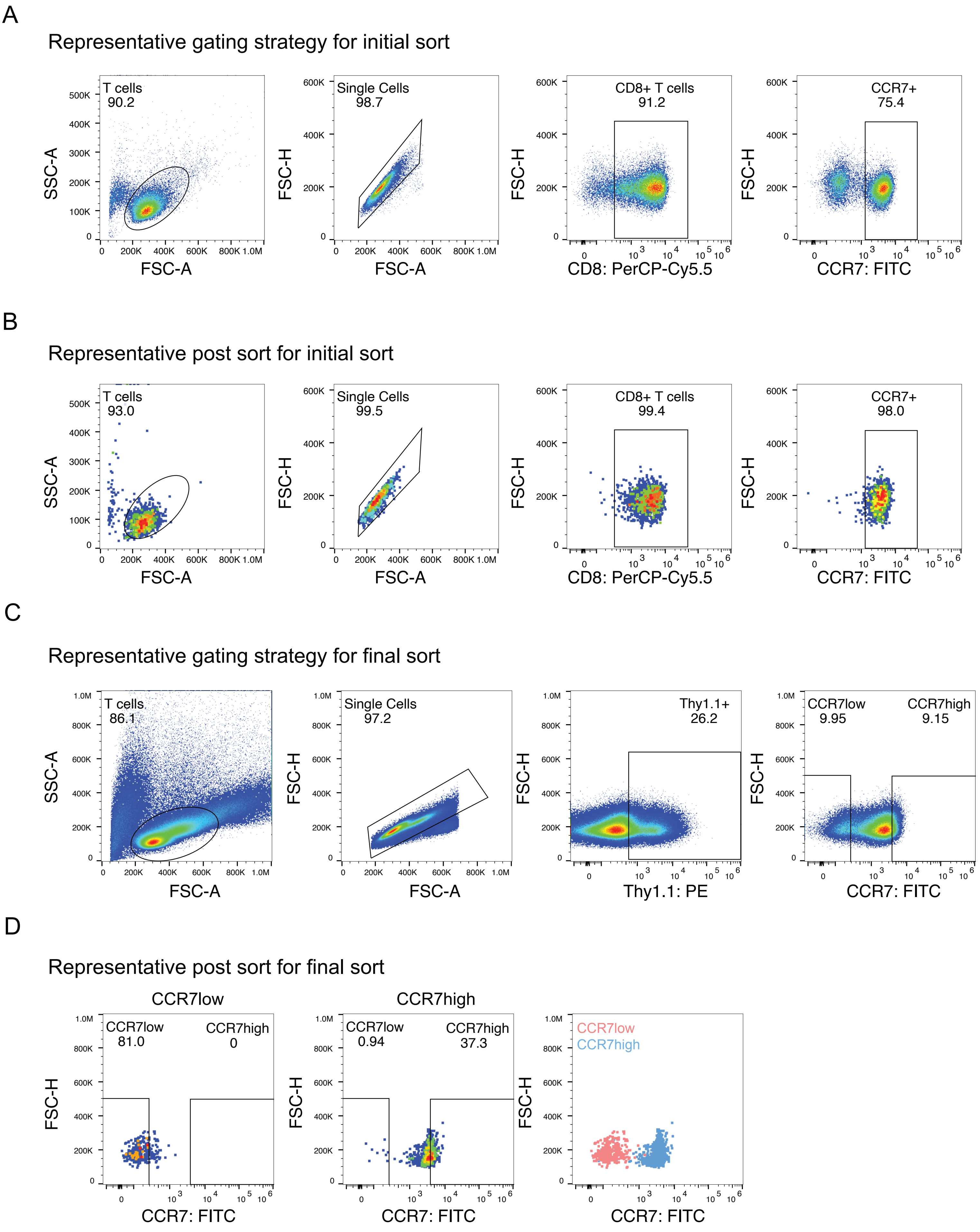
A
Representative gating strategy for initial sort
T cells
90.2
SSC-A
FSC-A
Single Cells
98.7
FSC-H
FSC-A
CD8+ T cells
91.2
FSC-H
CD8: PerCP-Cy5.5
CCR7+
75.4
FSC-H
CCR7: FITC
B
Representative post sort for initial sort
T cells
93.0
SSC-A
FSC-A
Single Cells
99.5
FSC-H
FSC-A
CD8+ T cells
99.4
FSC-H
CD8: PerCP-Cy5.5
CCR7+
98.0
FSC-H
CCR7: FITC
C
Representative gating strategy for final sort
T cells
86.1
SSC-A
FSC-A
Single Cells
97.2
FSC-H
FSC-A
Thy1.1+
26.2
FSC-H
Thy1.1: PE
CCR7low
9.95
CCR7high
9.15
FSC-H
CCR7: FITC
D
Representative post sort for final sort
CCR7low
CCR7low
81.0
CCR7high
0
FSC-H
CCR7: FITC
CCR7high
CCR7low
0.94
CCR7high
37.3
FSC-H
CCR7: FITC
CCR7low
CCR7high
FSC-H
CCR7: FITC

**Figure S6**

**Figure S6. Representative gating and post sorts for initial and final sorts for CRISPRi/a TF screens.**

**(A)** Representative initial gating for CD8+CCR7+ T cell population.

**(B)** Representative post sort of CD8+CCR7+ T cell population.

**(C)** Representative final gating strategy for transduced (Thy1.1+) cells in the lower and upper 10% tails of CCR7 expression on day 9 post-transduction.

**(D)** Representative post sorts of Thy1.1+CCR7-low (left) and Thy1.1+CCR7-high (middle) populations. Overlay of sorted populations (right) shows clear separation of CCR7-low and CCR7-high populations.

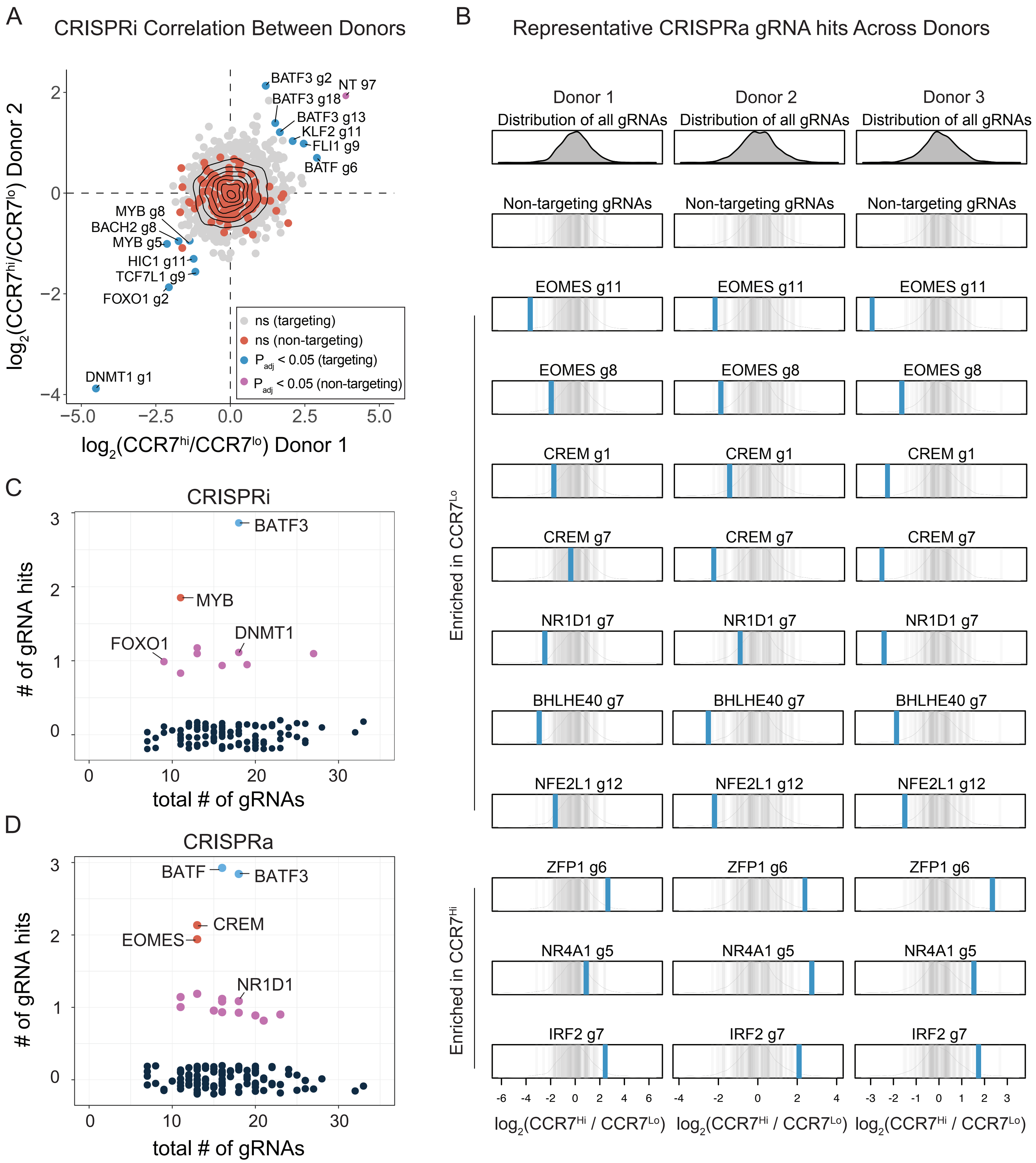
A
CRISPRi Correlation Between Donors
log2(CCR7hi/CCR7lo) Donor 1
log2(CCR7hi/CCR7lo) Donor 2
BATF3 g2
NT 97
BATF3 g18
BATF3 g13
KLF2 g11
FLI1 g9
BATF g6
MYB g8
BACH2 g8
MYB g5
HIC1 g11
TCF7L1 g9
FOXO1 g2
DNMT1 g1
ns (targeting)
ns (non-targeting)
Padj < 0.05 (targeting)
Padj < 0.05 (non-targeting)
B
Representative CRISPRa gRNA hits Across Donors
Donor 1
Donor 2
Donor 3
Distribution of all gRNAs
Non-targeting gRNAs
EOMES g11
EOMES g8
CREM g1
CREM g7
NR1D1 g7
BHLHE40 g7
NFE2L1 g12
ZFP1 g6
NR4A1 g5
IRF2 g7
Enriched in CCR7Lo
Enriched in CCR7Hi
log2(CCR7Hi / CCR7Lo)
C
CRISPRi
BATF3
MYB
FOXO1
DNMT1
### of gRNA hits
total # of gRNAs
D
CRISPRa
BATF
BATF3
CREM
EOMES
NR1D1
### of gRNA hits
total # of gRNAs

**Figure S7**

**Figure S7. Effects of CRISPRi/a gRNA hits across donors**

**(A)** Scatter plot of fold change in gRNA abundances between CCR7-high and CCR7-low populations across two donors in CRISPRi screens.

**(B)** Distribution of fold change in gRNA abundances between CCR7-high and CCR7-low populations for representative gRNA hits in the CRISPRa screen across three donors. Blue vertical lines represent gRNA hits and gray vertical lines represent the distribution of 120 non-targeting gRNAs.

Scatter plots of number of gRNA hits versus total number of gRNAs for each gene in **(C)** CRISPRi and **(D)** CRISPRa TF screens. A slight vertical stagger was implemented for visualization purposes, but there are only discrete values as denoted by the colors (0 gRNA hits = black, 1 gRNA hit = magenta, 2 gRNA hits = red, 3 gRNA hits = blue).

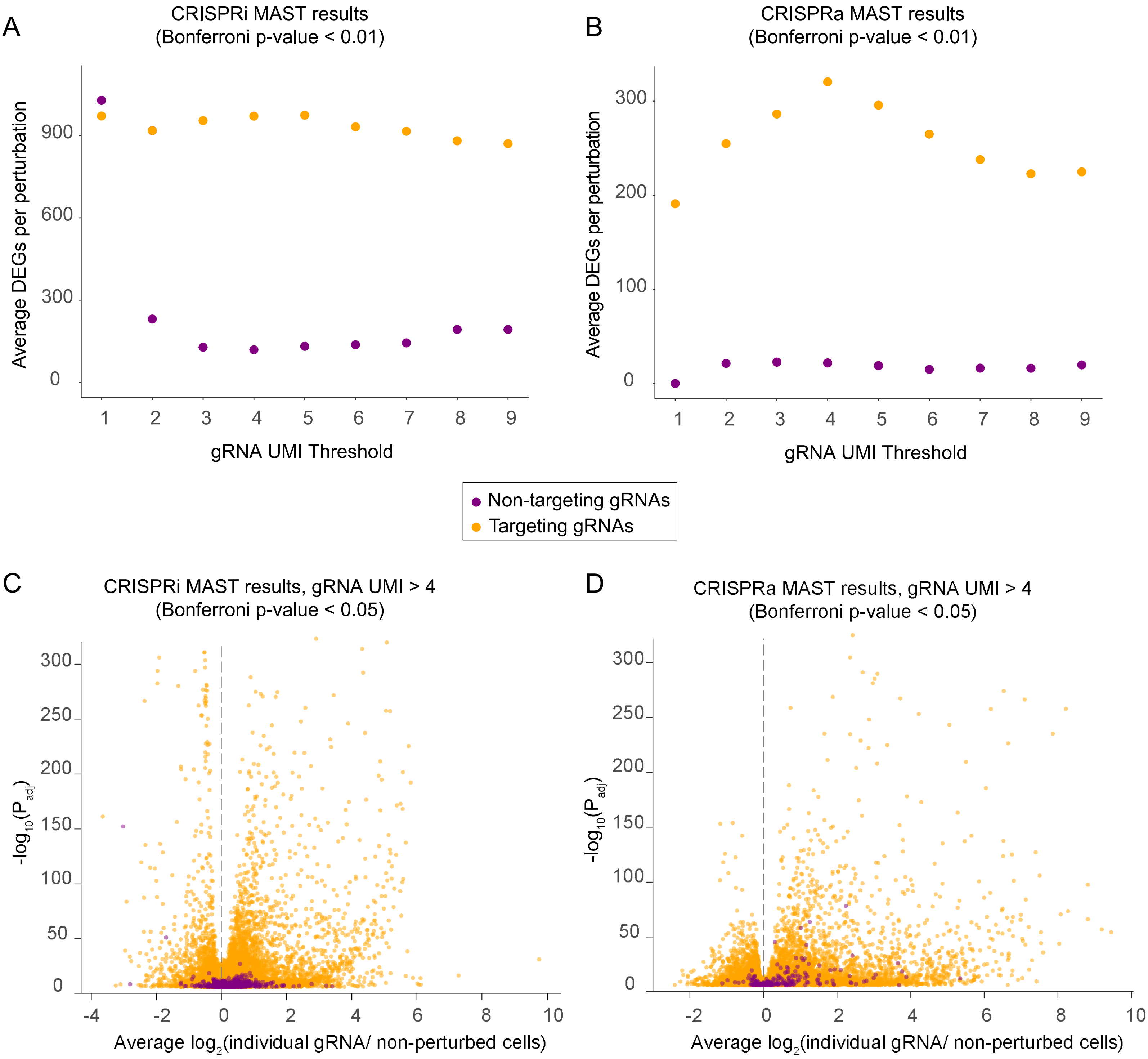

A
CRISPRi MAST results
(Bonferroni p-value < 0.01)
Average DEGs per perturbation
900
600
300
0
1 2 3 4 5 6 7 8 9
gRNA UMI Threshold
B
CRISPRa MAST results
(Bonferroni p-value < 0.01)
Average DEGs per perturbation
300
200
100
0
1 2 3 4 5 6 7 8 9
gRNA UMI Threshold
Non-targeting gRNAs
Targeting gRNAs
C
CRISPRi MAST results, gRNA UMI > 4
(Bonferroni p-value < 0.05)
-log10(Padj)
300
250
200
150
100
50
0
-4 -2 0 2 4 6 8 10
Average log2(individual gRNA/ non-perturbed cells)
D
CRISPRa MAST results, gRNA UMI > 4
(Bonferroni p-value < 0.05)
-log10(Padj)
300
250
200
150
100
50
0
-2 0 2 4 6 8 10
Average log2(individual gRNA/ non-perturbed cells)


**Figure S8**

**Figure S8. Quality control of differential gene expression analyses for CRISPRi and CRISPRa TF scRNA-seq characterization.**

The average number of differentially expressed genes (DEGs) for targeting and non-targeting gRNAs as a function of gRNA UMI threshold used for gRNA assignment to cells in **(A)** CRISPRi and **(B)** CRISPRa scRNA-seq screens.

Volcano plots of **(C)** CRISPRi and **(D)** CRISPRa scRNA-seq screens with the statistical significance ($P_{adj}$) of each significant gRNA-gene pair plotted versus the fold change in gene expression relative to non-perturbed cells.

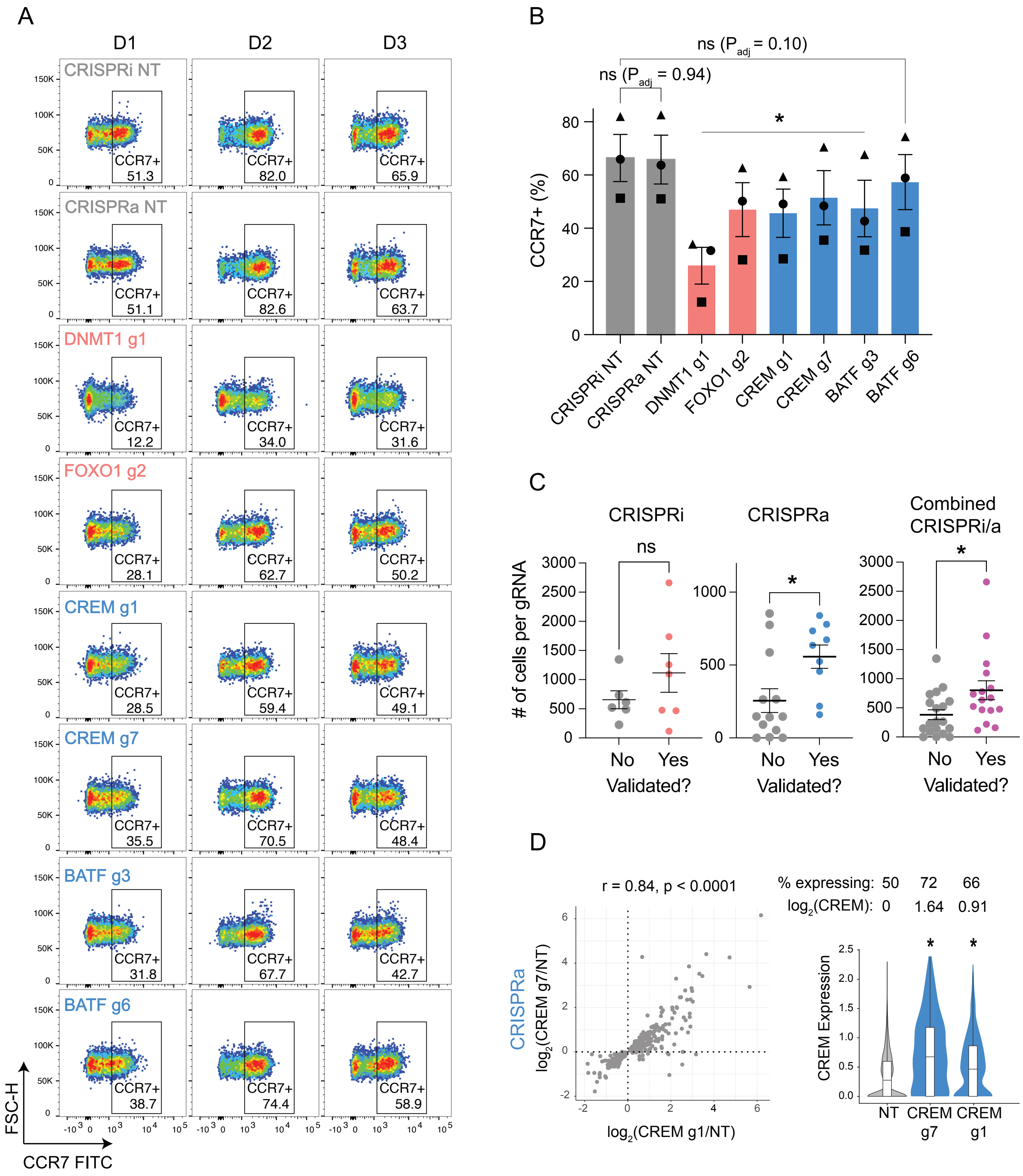
A
D1
D2
D3
CRISPRi NT
CRISPRa NT
DNMT1 g1
FOXO1 g2
CREM g1
CREM g7
BATF g3
BATF g6
CCR7+
51.3
82.0
65.9
51.1
82.6
63.7
12.2
34.0
31.6
28.1
62.7
50.2
28.5
59.4
49.1
35.5
70.5
48.4
31.8
67.7
42.7
38.7
74.4
58.9
FSC-H
CCR7 FITC
B
ns (Padj = 0.10)
ns (Padj = 0.94)
*
CCR7+ (%)
CRISPRi NT
CRISPRa NT
DNMT1 g1
FOXO1 g2
CREM g1
CREM g7
BATF g3
BATF g6
C
CRISPRi
CRISPRa
Combined CRISPRi/a
ns
*
*
### of cells per gRNA
No
Yes
Validated?
D
r = 0.84, p < 0.0001
CRISPRa
log2(CREM g7/NT)
log2(CREM g1/NT)
% expressing: 50 72 66
log2(CREM): 0 1.64 0.91
CREM Expression
NT
CREM g7
CREM g1

### Figure S9

**Figure S9. Individual validation of subset of CRISPRi/a gRNAs on CCR7 expression.**

**(A)** Flow cytometry plots of CCR7 expression in CD8+ T cells across three donors with the indicated CRISPRi/a perturbations.

**(B)** Summary statistics of percent CCR7+ T cells across CRISPRi/a perturbations. A paired, one-way ANOVA test was used to compare the percentage of CCR7+ T cells for each perturbation to the CRISPRi NT treatment group (n = 3 donors, error bars represent SEM, different shapes are used to denote each donor).

**(C)** Number of cells assigned to each gRNA across CRISPRi (left), CRISPRa (middle), and joint CRISPRi/a (right) scRNA-seq datasets with gRNAs stratified based on whether they affected CCR7 expression as predicted by the flow-based screen. A Mann-Whitney test was used to determine statistical significance for each group.

**(D)** Correlation of the union set of DEGs between CREM gRNAs from CRISPRa scRNA-seq characterization (left). Violin plots of CREM expression across cells assigned to indicated gRNA with the fold change in target gene expression relative to NT and the percent of cells expressing the target gene indicated above (right).

A

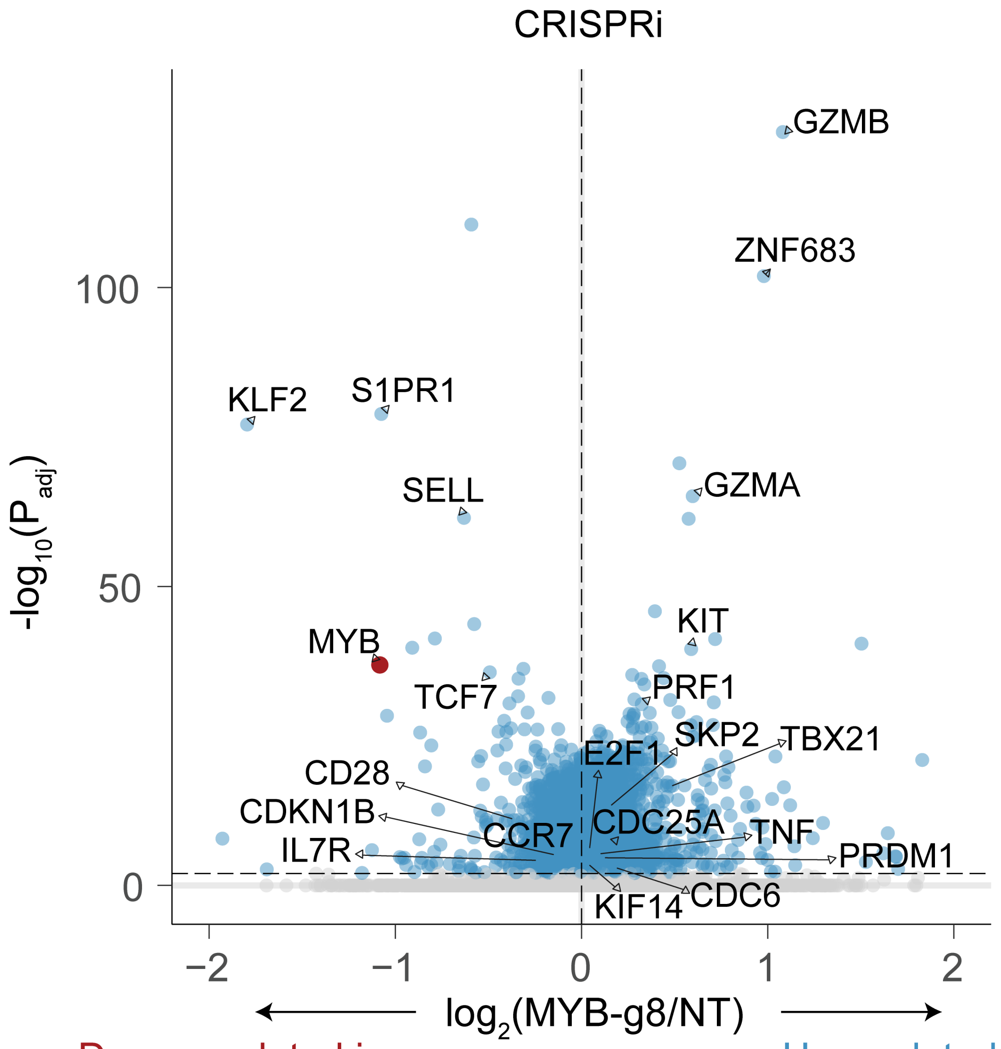


B

Downregulated in MYB-silenced cells

Upregulated in MYB-silenced cells

| Downregulated in MYB-silenced cells | Upregulated in MYB-silenced cells |
| --- | --- |
| Memory TFs and surface markers<br>-MYB, TCF7, KLF2, IL7R | Effector TFs<br>-TBX21, PRDM1, ZNF683 |
| Lymph homing genes<br>-CCR7, SELL, S1PR1 | Effector molecules<br>-GZMB, GZMA, PRF1, TNF, KIT |
| Negative cell cycle regulators<br>-CDKN1B | Positive cell cycle regulators<br>-E2F1, CDC6, SKP2, CDC25A, KIF14 |

**Figure S10**

**Figure S10. MYB silencing drives T cells towards an effector phenotype.**

**(A)** Volcano plot of statistical significance ($P_{adj}$) for each gene versus the fold change in gene expression in MYB CRISPRi-perturbed cells relative to non-perturbed cells. All DEGs ($P_{adj} < 0.01$) are labeled blue apart from MYB, which is labeled dark red. Selected DEGs are annotated.

**(B)** Classification of annotated DEGs based on their functional role.

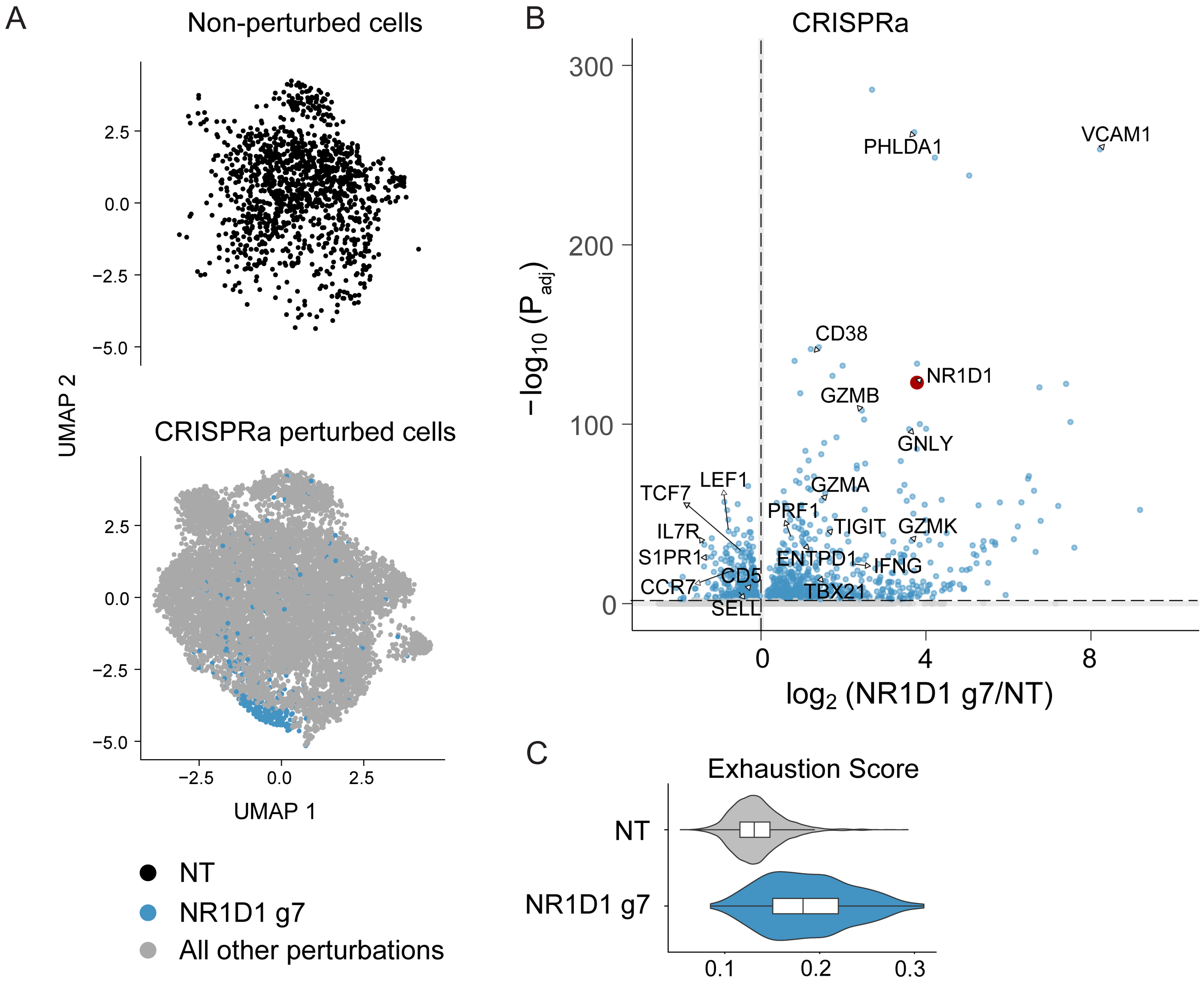

A
Non-perturbed cells
CRISPRa perturbed cells
UMAP 2
UMAP 1
NT
NR1D1 g7
All other perturbations
B
CRISPRa
VCAM1
PHLDA1
CD38
NR1D1
GZMB
GNLY
LEF1
TCF7
GZMA
PRF1
IL7R
TIGIT
GZMK
S1PR1
ENTPD1
IFNG
CCR7
CD5
TBX21
SELL
$-\log_{10}(P_{adj})$
$\log_2$ (NR1D1 g7/NT)
C
Exhaustion Score
NT
NR1D1 g7


**Figure S11**

**Figure S11. NR1D1 synthetically induces exhaustion phenotype.**

**(A)** UMAP plot of CRISPRa scRNA-seq characterization with cells split by perturbation status: non-perturbed (top) and perturbed (bottom). Blue data points indicate cells with a NR1D1 gRNA. Cells were clustered using Seurat's CalcPerturbSig function to mitigate confounding sources of variation such as the donor and phase of cell cycle.

**(B)** Volcano plot of statistical significance ($P_{adj}$) for each gene versus the fold change in gene expression in NR1D1 CRISPRa-perturbed cells relative to non-perturbed cells. All DEGs ($P_{adj} < 0.01$) are labeled blue apart from NR1D1, which is labeled dark red. Selected DEGs are annotated.

**(C)** Violin plot of exhaustion gene signature score across all non-perturbed and NR1D1-perturbed cells in the CRISPRa scRNA-seq screen. UCell gene signature scores are based on the Mann-Whitney U statistic.

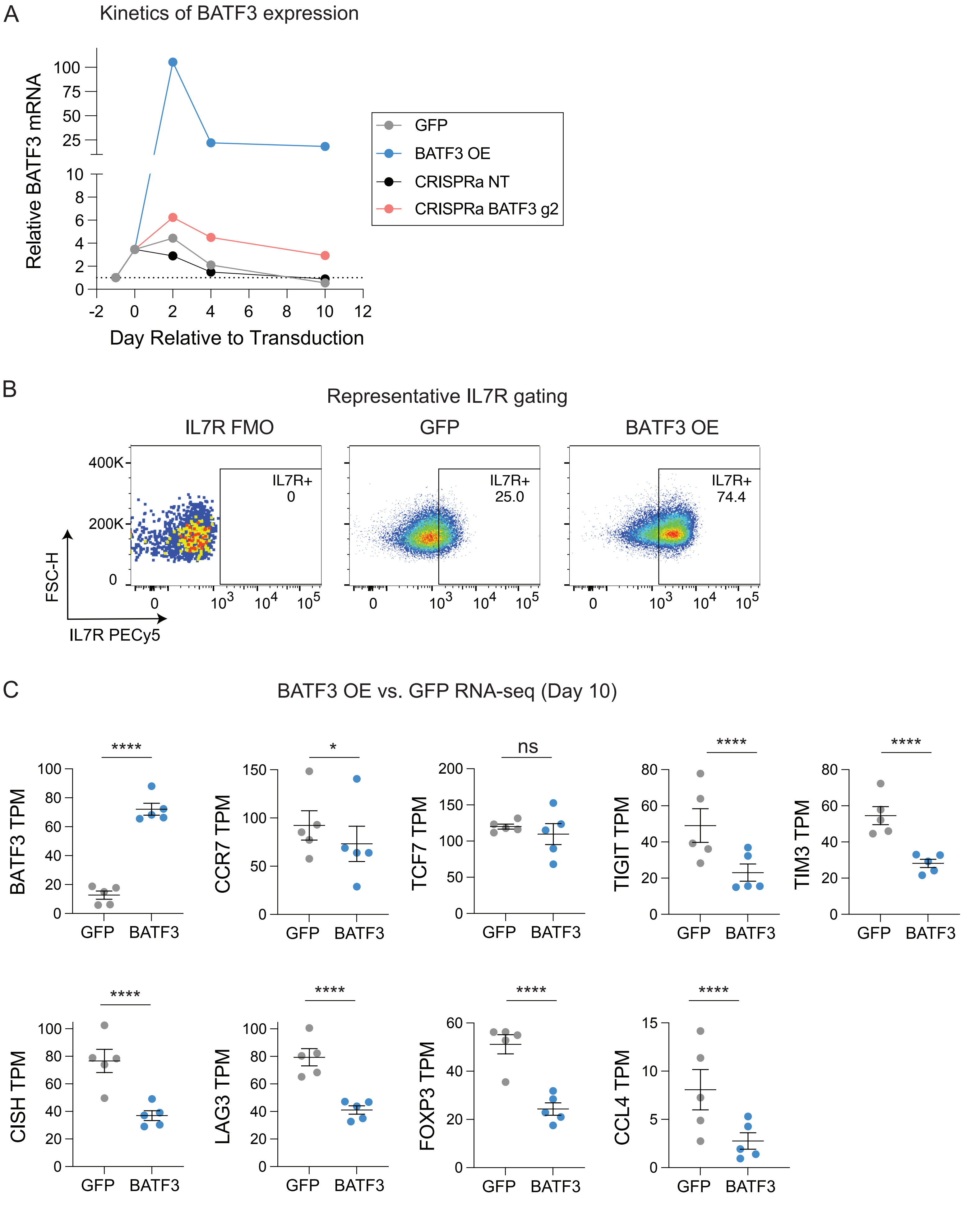
A
Kinetics of BATF3 expression
Relative BATF3 mRNA
Day Relative to Transduction
GFP
BATF3 OE
CRISPRa NT
CRISPRa BATF3 g2
B
Representative IL7R gating
IL7R FMO
GFP
BATF3 OE
IL7R+
0
IL7R+
25.0
IL7R+
74.4
FSC-H
IL7R PECy5
C
BATF3 OE vs. GFP RNA-seq (Day 10)
BATF3 TPM
CCR7 TPM
TCF7 TPM
TIGIT TPM
TIM3 TPM
CISH TPM
LAG3 TPM
FOXP3 TPM
CCL4 TPM
GFP
BATF3
****
*
ns

**Figure S12**

**Figure S12. Kinetics of BATF3 expression and effects of BATF3 OE.**

(**A**) Median BATF3 expression as a function of time relative to BATF3 expression before T cell activation across four treatment groups (n = 3 donors, fold change in BATF3 expression was calculated using $2^{-dCT}$ method relative to baseline BATF3 expression, internal householding control was excluded because T cell stimulation dramatically alters expression of householding genes such as GAPDH and TBP, input mass of RNA into the reverse transcription reaction was the same for all samples).

**(B)** An IL7R fluorescent minus one (FMO, left) control was used to set the IL7R+ gate. Representative IL7R expression of CD8+ T cells from a donor transduced with either GFP (middle) or BATF3 OE (right) on day 8 post-transduction.

**(C)** Transcripts per million (TPM) of selected genes across n = 5 donors with either GFP or BATF3 OE on day 10 post transduction. Statistical significance for each gene was determined using paired DESeq2 analysis between treatment groups.

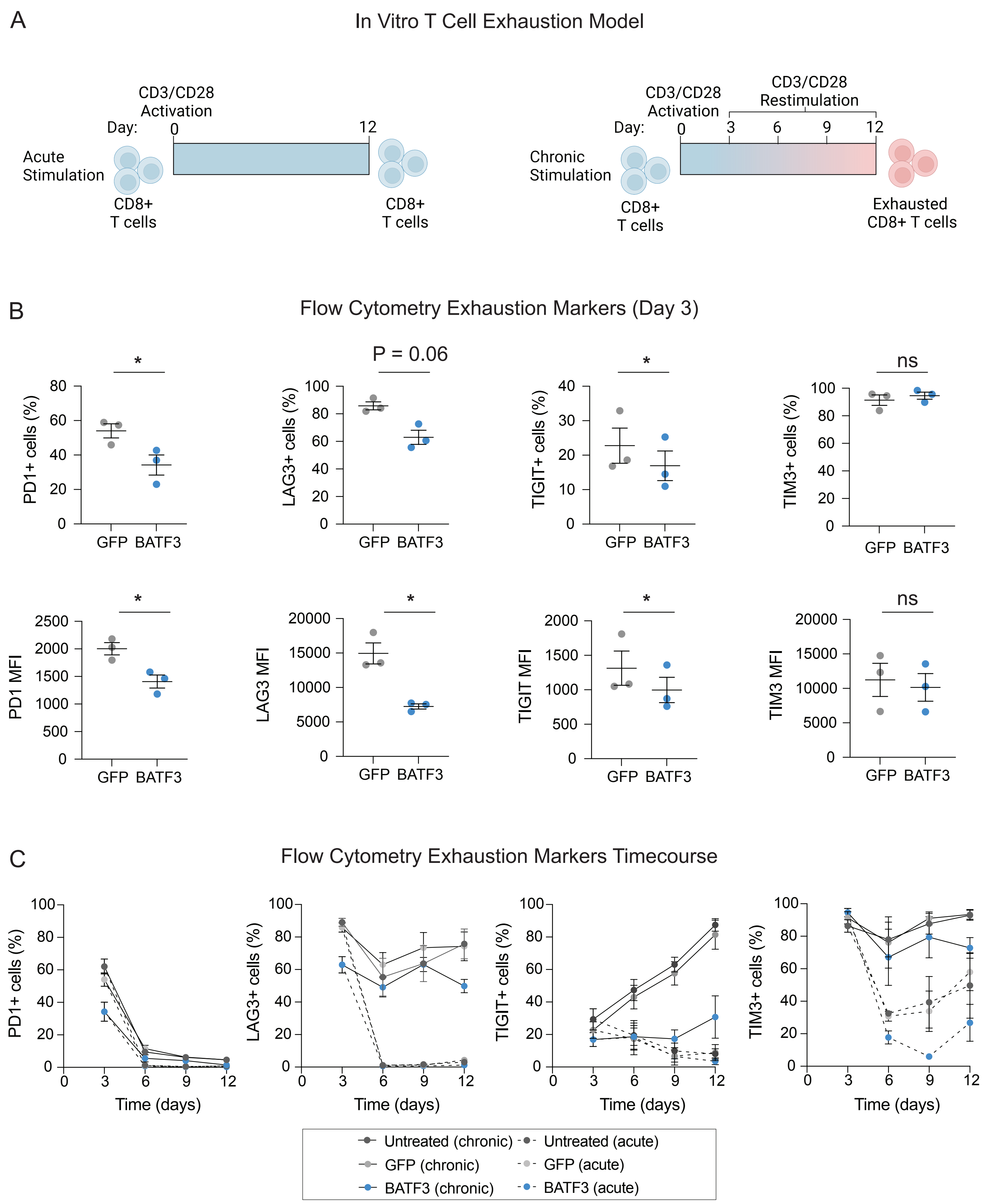
A
In Vitro T Cell Exhaustion Model
CD3/CD28 Activation
Day: 0
12
Acute Stimulation
CD8+ T cells
CD8+ T cells
CD3/CD28 Restimulation
CD3/CD28 Activation
Day: 0
3
6
9
12
Chronic Stimulation
CD8+ T cells
Exhausted CD8+ T cells
B
Flow Cytometry Exhaustion Markers (Day 3)
*
PD1+ cells (%)
P = 0.06
LAG3+ cells (%)
*
TIGIT+ cells (%)
ns
TIM3+ cells (%)
GFP BATF3
*
PD1 MFI
*
LAG3 MFI
*
TIGIT MFI
ns
TIM3 MFI
C
Flow Cytometry Exhaustion Markers Timecourse
PD1+ cells (%)
LAG3+ cells (%)
TIGIT+ cells (%)
TIM3+ cells (%)
Time (days)
Untreated (chronic)
Untreated (acute)
GFP (chronic)
GFP (acute)
BATF3 (chronic)
BATF3 (acute)

**Figure S13**

**Figure S13. BATF3 OE attenuates expression of T cell exhaustion markers in chronically stimulated CD8+ T cells.**

**(A)** Schematic of acute (left) and chronic stimulation (right) with CD3/CD28 dynabeads.

**(B)** Average percentage of positive cells (top panel) and mean fluorescence intensity (MFI, bottom panel) of exhaustion markers: PD1, LAG3, TIGIT, and TIM3 on day 3 post-transduction with GFP or BATF3 OE ($n = 3$ individual donors, error bars represent SEM, paired t tests used to determine statistical significance).

**(C)** Time course of PD1, LAG3, TIGIT, and TIM3 expression post-transduction with GFP or BATF3 OE under acute or chronic stimulation ($n = 3$ individual donors, error bars represent SEM).

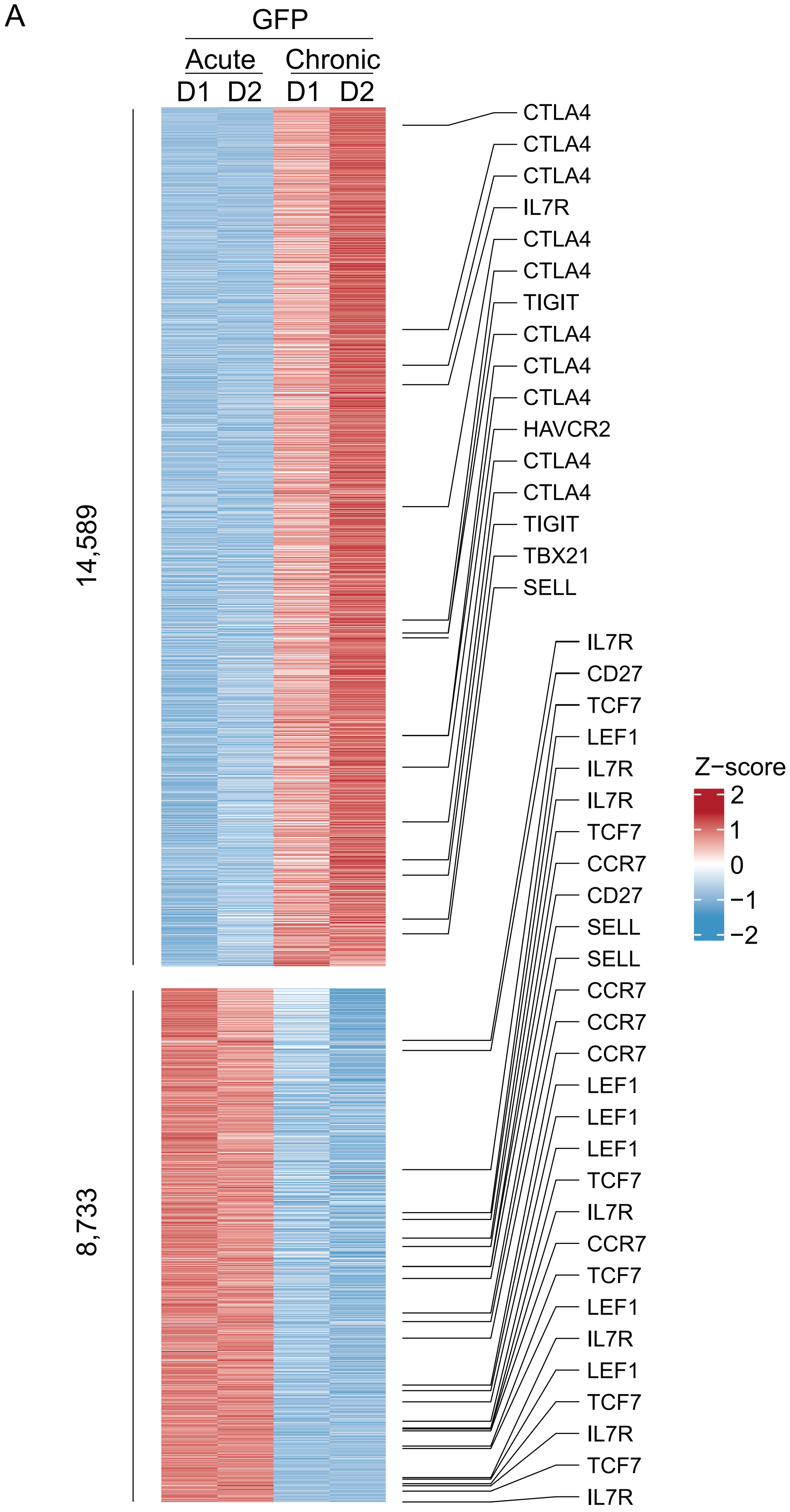
A
GFP
Acute
Chronic
D1
D2
D1
D2
14,589
8,733
CTLA4
CTLA4
CTLA4
IL7R
CTLA4
CTLA4
TIGIT
CTLA4
CTLA4
CTLA4
HAVCR2
CTLA4
CTLA4
TIGIT
TBX21
SELL
IL7R
CD27
TCF7
LEF1
IL7R
IL7R
TCF7
CCR7
CD27
SELL
SELL
CCR7
CCR7
CCR7
LEF1
LEF1
LEF1
TCF7
IL7R
CCR7
TCF7
LEF1
IL7R
LEF1
TCF7
IL7R
TCF7
IL7R
Z−score
2
1
0
−1
−2

**Figure S14**

**Figure S14. Chronic antigen stimulation drives extensive chromatin remodeling in control CD8+ T cells.**

**(A)** Heatmap of differentially accessible regions with selected regions annotated with their nearest gene.

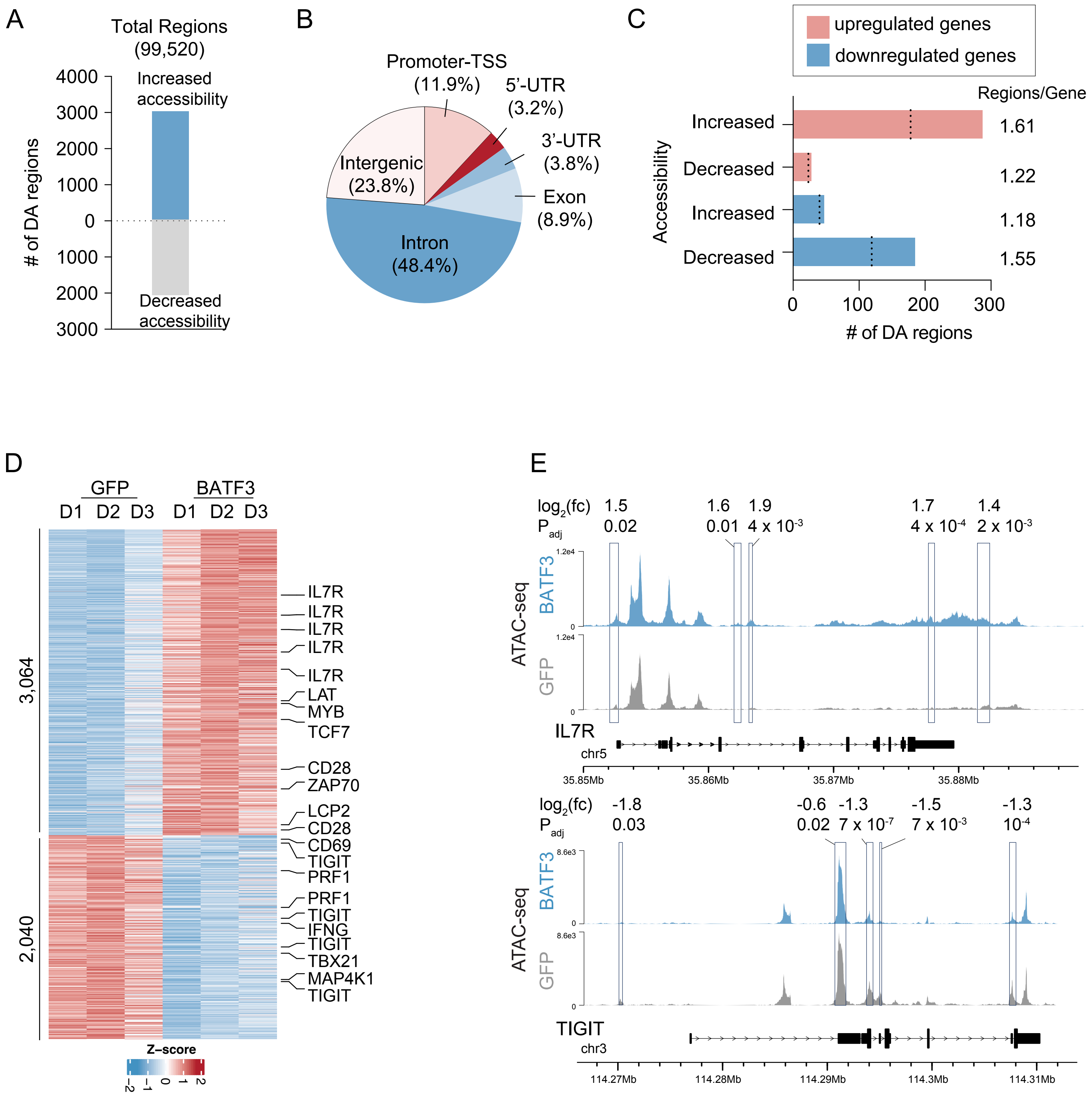

A
Total Regions (99,520)
Increased accessibility
Decreased accessibility
### of DA regions
4000
3000
2000
1000
0
1000
2000
3000
B
Promoter-TSS (11.9%)
5'-UTR (3.2%)
3'-UTR (3.8%)
Exon (8.9%)
Intron (48.4%)
Intergenic (23.8%)
C
upregulated genes
downregulated genes
Regions/Gene
Accessibility
Increased
Decreased
Increased
Decreased
1.61
1.22
1.18
1.55
0
100
200
300
### of DA regions
D
GFP
BATF3
D1 D2 D3 D1 D2 D3
3,064
2,040
IL7R
IL7R
IL7R
IL7R
IL7R
LAT
MYB
TCF7
CD28
ZAP70
LCP2
CD28
CD69
TIGIT
PRF1
PRF1
TIGIT
IFNG
TIGIT
TBX21
MAP4K1
TIGIT
Z-score
E
log2(fc) 1.5 1.6 1.9 1.7 1.4
Padj 0.02 0.01 4 x 10-3 4 x 10-4 2 x 10-3
ATAC-seq
BATF3
GFP
1.2e4
0
IL7R
chr5
35.85Mb
35.86Mb
35.87Mb
35.88Mb
log2(fc) -1.8 -0.6 -1.3 -1.5 -1.3
Padj 0.03 0.02 7 x 10-7 7 x 10-3 10-4
8.6e3
TIGIT
chr3
114.27Mb
114.28Mb
114.29Mb
114.3Mb
114.31Mb


### Figure S15

**Figure S15. BATF3 remodels the chromatin landscape of CD8+ T cells under acute stimulation.**

**(A)** Number of ATAC-seq regions with increased or decreased accessibility in CD8+ T cells (n = 3 individual donors, differential accessible regions defined as $P_{adj} < 0.05$) with BATF3 OE on day 14 post-transduction.

**(B)** Proportion of differentially accessible (DA) regions based on genomic feature classification.

**(C)** Joint analysis of RNA-seq and ATAC-seq datasets. Number of differentially accessible regions near upregulated and downregulated genes. Dashed lines represent the number of unique DEGs associated with DA regions.

**(D)** Heatmap of DA regions with selected regions annotated with their nearest gene.

**(E)** Representative ATAC-seq tracks at *IL7R* and *TIGIT* loci with overlayed rectangles indicating DA regions.

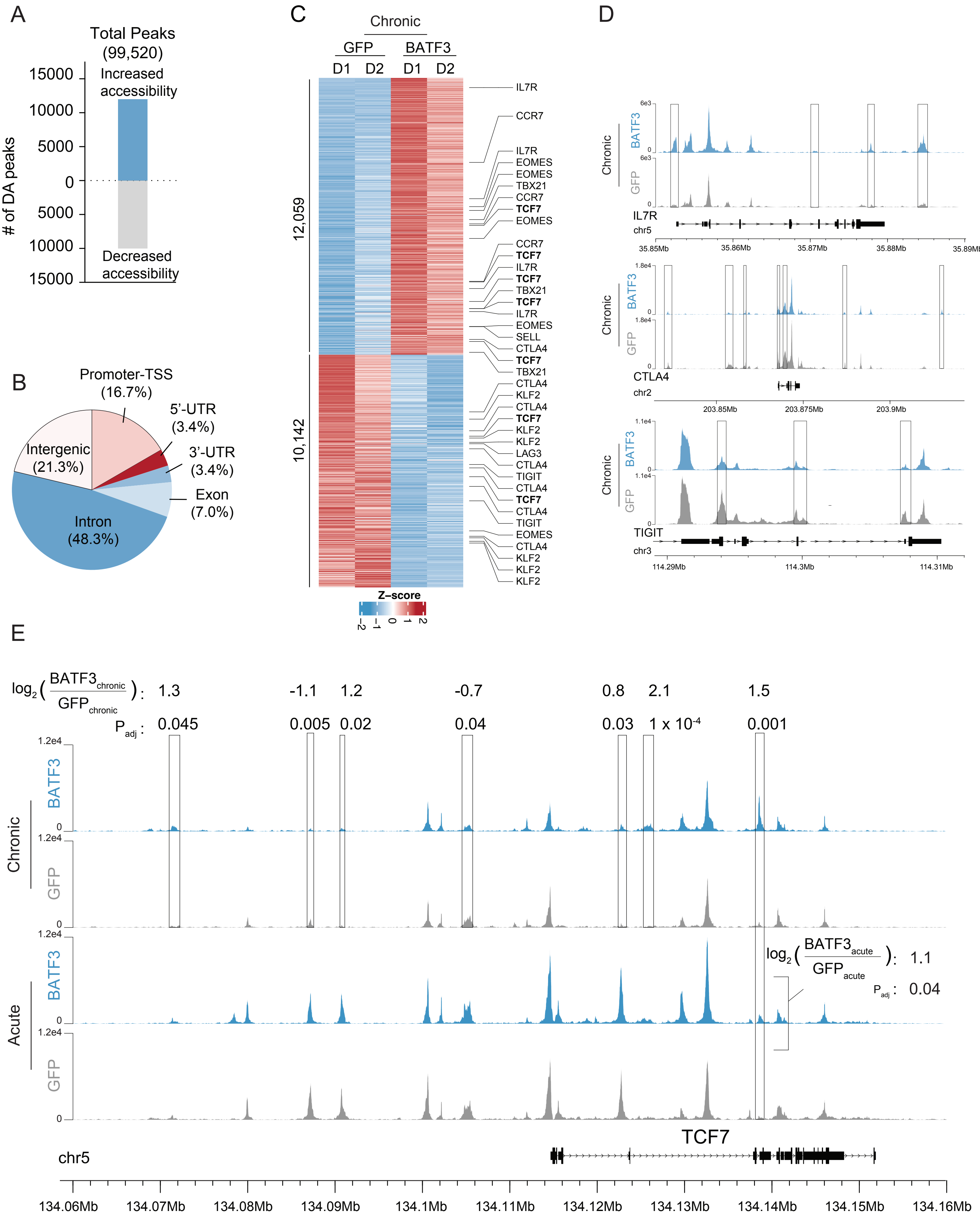
A
Total Peaks (99,520)
Increased accessibility
Decreased accessibility
### of DA peaks
B
Promoter-TSS (16.7%)
5'-UTR (3.4%)
3'-UTR (3.4%)
Exon (7.0%)
Intron (48.3%)
Intergenic (21.3%)
C
Chronic
GFP
BATF3
D1 D2 D1 D2
12,059
10,142
Z-score
IL7R
CCR7
EOMES
TBX21
TCF7
SELL
CTLA4
KLF2
LAG3
TIGIT
D
Chronic
BATF3
GFP
IL7R
chr5
CTLA4
chr2
TIGIT
chr3
E
Chronic
Acute
BATF3
GFP
TCF7
chr5

### Figure S16

**Figure S16. BATF3 remodels the chromatin landscape of CD8+ T cells under chronic stimulation.**

**(A)** Number of ATAC-seq regions with increased or decreased accessibility in HER2-CAR-2A-BATF3 CD8+ T cells compared to HER2-CAR-2A-GFP CD8+ T cells on day 14 post-transduction after repeated rounds of tumor restimulation (n = 2 individual donors, differential accessible regions defined as $P_{adj} < 0.05$).

**(B)** Proportion of differentially accessible (DA) regions based on genomic feature classification.

**(C)** Heatmap of DA regions with selected regions annotated with their nearest gene.

**(D)** Representative ATAC-seq tracks at *IL7R*, *CTLA4*, *TIGIT* loci with overlayed rectangles indicating DA regions.

**(E)** Representative ATAC-seq tracks of the TCF7 locus under acute and chronic stimulation with and without BATF3 OE.

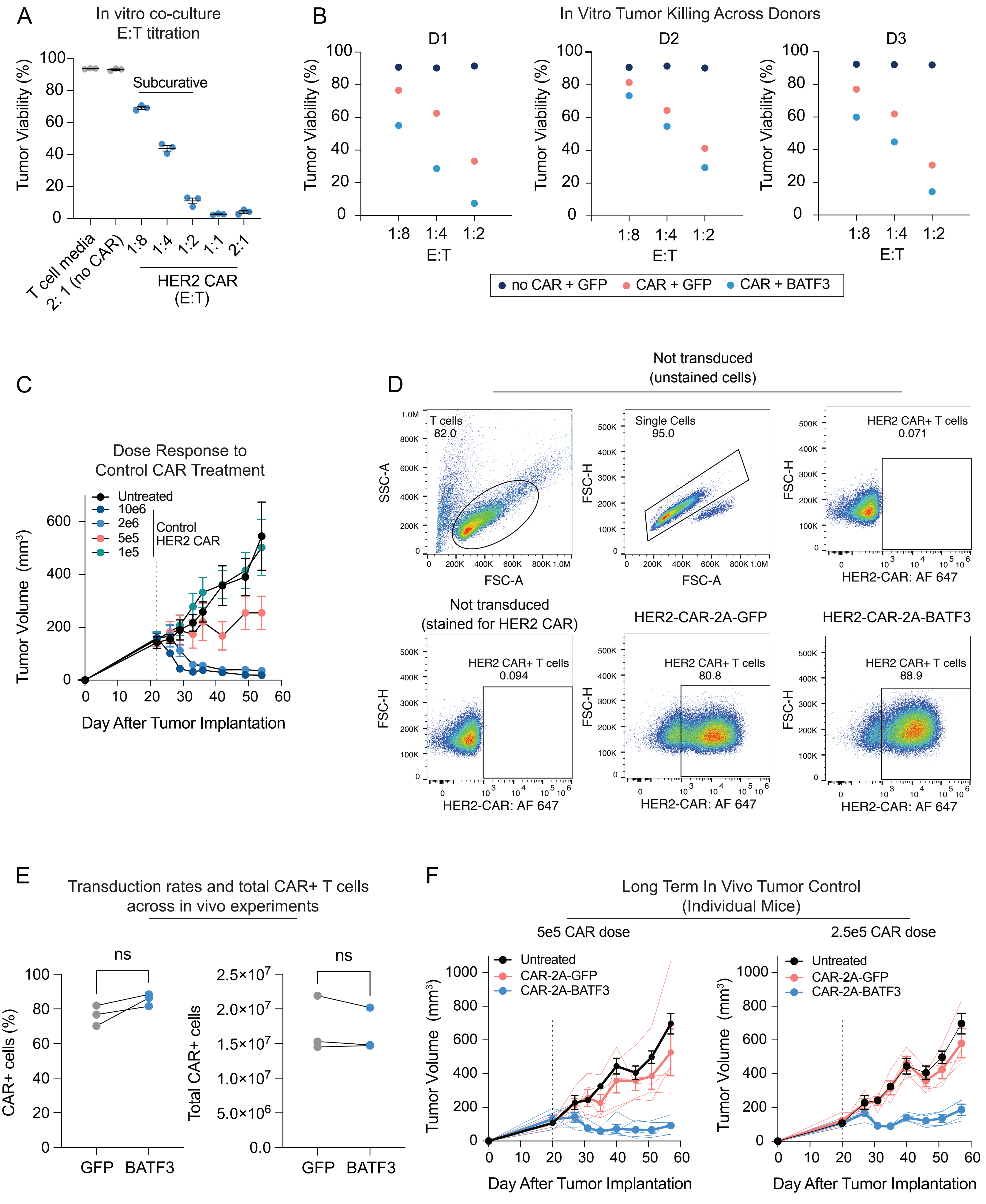

A
In vitro co-culture
E:T titration
Tumor Viability (%)
Subcurative
T cell media
2: 1 (no CAR)
1:8
1:4
1:2
1:1
2:1
HER2 CAR
(E:T)
B
In Vitro Tumor Killing Across Donors
D1
D2
D3
Tumor Viability (%)
E:T
no CAR + GFP
CAR + GFP
CAR + BATF3
C
Dose Response to
Control CAR Treatment
Untreated
10e6
2e6
5e5
1e5
Control
HER2 CAR
Tumor Volume (mm³)
Day After Tumor Implantation
D
Not transduced
(unstained cells)
T cells
82.0
SSC-A
FSC-A
Single Cells
95.0
FSC-H
HER2 CAR+ T cells
0.071
HER2-CAR: AF 647
Not transduced
(stained for HER2 CAR)
HER2 CAR+ T cells
0.094
HER2-CAR-2A-GFP
HER2 CAR+ T cells
80.8
HER2-CAR-2A-BATF3
HER2 CAR+ T cells
88.9
E
Transduction rates and total CAR+ T cells
across in vivo experiments
ns
CAR+ cells (%)
Total CAR+ cells
GFP
BATF3
F
Long Term In Vivo Tumor Control
(Individual Mice)
5e5 CAR dose
2.5e5 CAR dose
Untreated
CAR-2A-GFP
CAR-2A-BATF3
Tumor Volume (mm³)
Day After Tumor Implantation


### Figure S17

**Figure S17. BATF3 OE in human CD8+ T cells enhances in vitro and in vivo tumor control.**

**(A)** Tumor viability after 24 hours of culture in T cell media, co-culture with $CAR^{null}$ T cells, or co-culture with a titration of CAR+ T cell doses ranging from 1:8 to 2:1 E:T (n = 3 donors, error bars represent SEM).

**(B)** Tumor viability after 24 hours of co-culture with GFP $CAR^{null}$, GFP CAR+, and BATF3 OE CAR+ CD8 T cells at indicated effector to target (E:T) cell ratios for each donor.

**(C)** Tumor volume over time as a function of the dose of control HER2 CAR T cells (n = 5 mice per treatment, error bars represent SEM). Mice were intravenously injected with CAR T cells on day 21.

**(D)** Representative flow plots of CAR expression in CD8+ T cells with control and BATF3 OE CAR lentiviral plasmids on day 9 post-transduction (the same day that the mice were intravenously injected with CAR T cells).

**(E)** Summary statistics of transduction rates and total CAR+ T cells with control and BATF3 OE CAR lentiviral plasmids on day 9 post-transduction (n = 3 donors, lines connect donors across treatments)

**(F)** Tumor volumes of individual mice treated with $5 \times 10^5$ (left panel, n = 5 mice per treatment group) or $2.5 \times 10^5$ (right panel, n = 4 mice per treatment group) CAR T cells with or without BATF3 overexpression. Thinner lines represent individual mice and thicker lines represent the average tumor volume across mice in a treatment group.

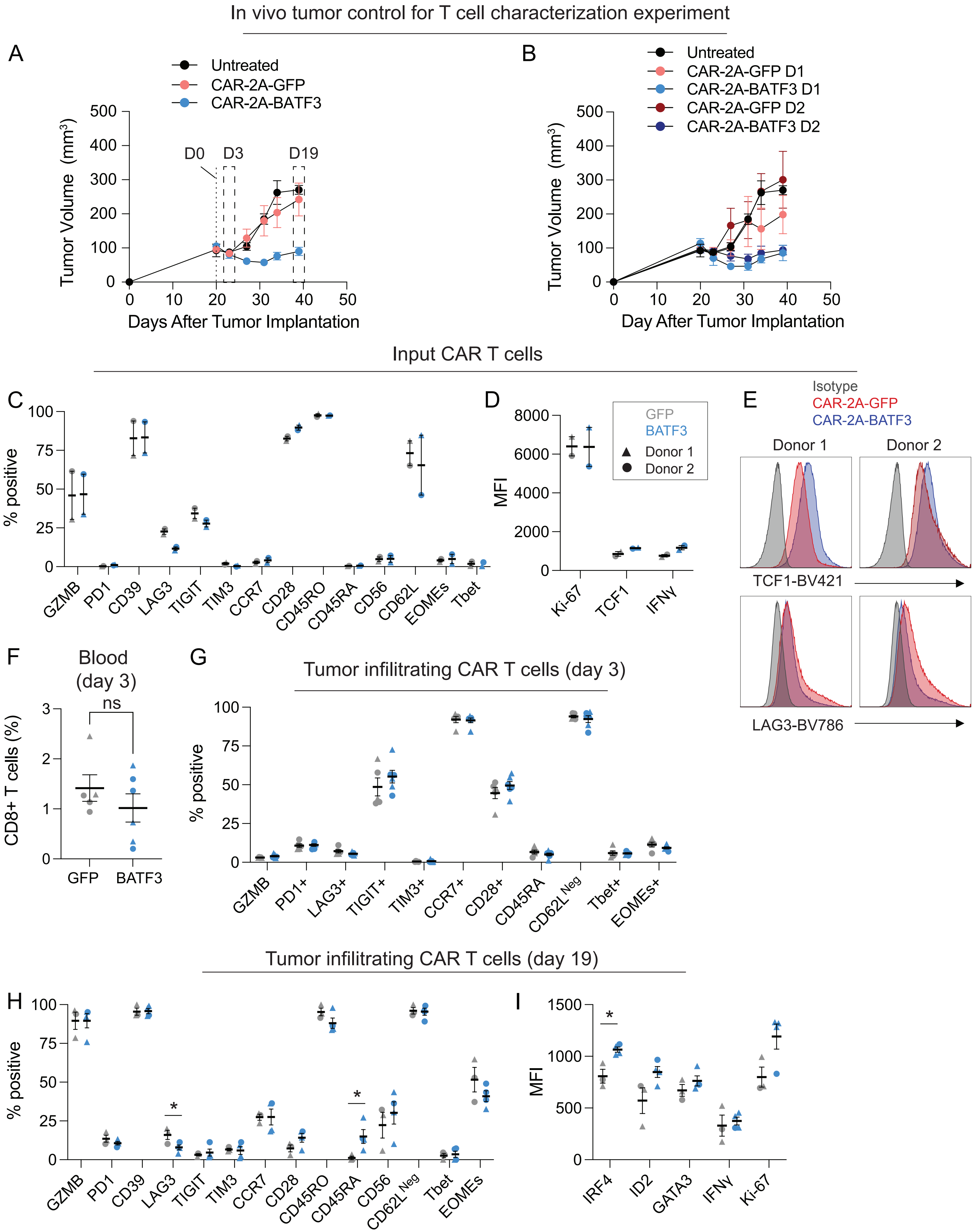
In vivo tumor control for T cell characterization experiment
A
Untreated
CAR-2A-GFP
CAR-2A-BATF3
Tumor Volume (mm³)
D0
D3
D19
Days After Tumor Implantation
B
CAR-2A-GFP D1
CAR-2A-BATF3 D1
CAR-2A-GFP D2
CAR-2A-BATF3 D2
Day After Tumor Implantation
Input CAR T cells
C
% positive
GZMB PD1 CD39 LAG3 TIGIT TIM3 CCR7 CD28 CD45RO CD45RA CD56 CD62L EOMEs Tbet
D
MFI
GFP
BATF3
Donor 1
Donor 2
Ki-67 TCF1 IFNγ
E
Isotype
CAR-2A-GFP
CAR-2A-BATF3
Donor 1
Donor 2
TCF1-BV421
LAG3-BV786
F
Blood (day 3)
ns
CD8+ T cells (%)
GFP BATF3
G
Tumor infiltrating CAR T cells (day 3)
GZMB PD1+ LAG3+ TIGIT+ TIM3+ CCR7+ CD28+ CD45RA CD62L Neg Tbet+ EOMEs+
Tumor infiltrating CAR T cells (day 19)
H
GZMB PD1 CD39 LAG3 TIGIT TIM3 CCR7 CD28 CD45RO CD45RA CD56 CD62L Neg Tbet EOMEs
I
IRF4 ID2 GATA3 IFNγ Ki-67

**Figure S18**

**Figure S18. Characterization of CAR T cells with or without BATF3 overexpression at multiple time points during in vivo tumor control experiment.**

**(A)** Tumor volume over time for untreated mice and mice treated with 5 x $10^5$ CAR T cells with or without BATF3 overexpression (n = 2 donors, 3-4 mice per donor, error bars represent SEM). Flow analysis on input CAR T cells and tumor infiltrating CAR T cells at day 3 and day 19 post-treatment.

**(B)** Same as **(A)** except stratified based on donor.

**(C)** Percentage of positive cells for indicated markers of input CAR T cells across groups

**(D)** MFI for indicated markers of input CAR T cells across groups.

**(E)** Histograms of TCF1 and LAG3 expression for input CAR T cells.

**(F)** Average percentage of CD8+ T cells in peripheral blood on day 3 post-treatment across groups. A Mann-Whitney test was used to compare the percentage of CD8+ cells between the two groups.

**(G)** Percentage of positive cells for indicated markers of tumor infiltrating CAR T cells on day 3 post-treatment across groups.

**(H)** Percentage of positive cells for indicated markers of tumor infiltrating CAR T cells on day 19 post-treatment across groups. An unpaired t test was used to compare expression of each marker between groups.

**(I)** MFI for indicated markers of tumor infiltrating CAR T cells on day 19 post-treatment across groups. An unpaired t test was used to compare expression of each marker between groups.

Error bars represent SEM for all panels in this figure.

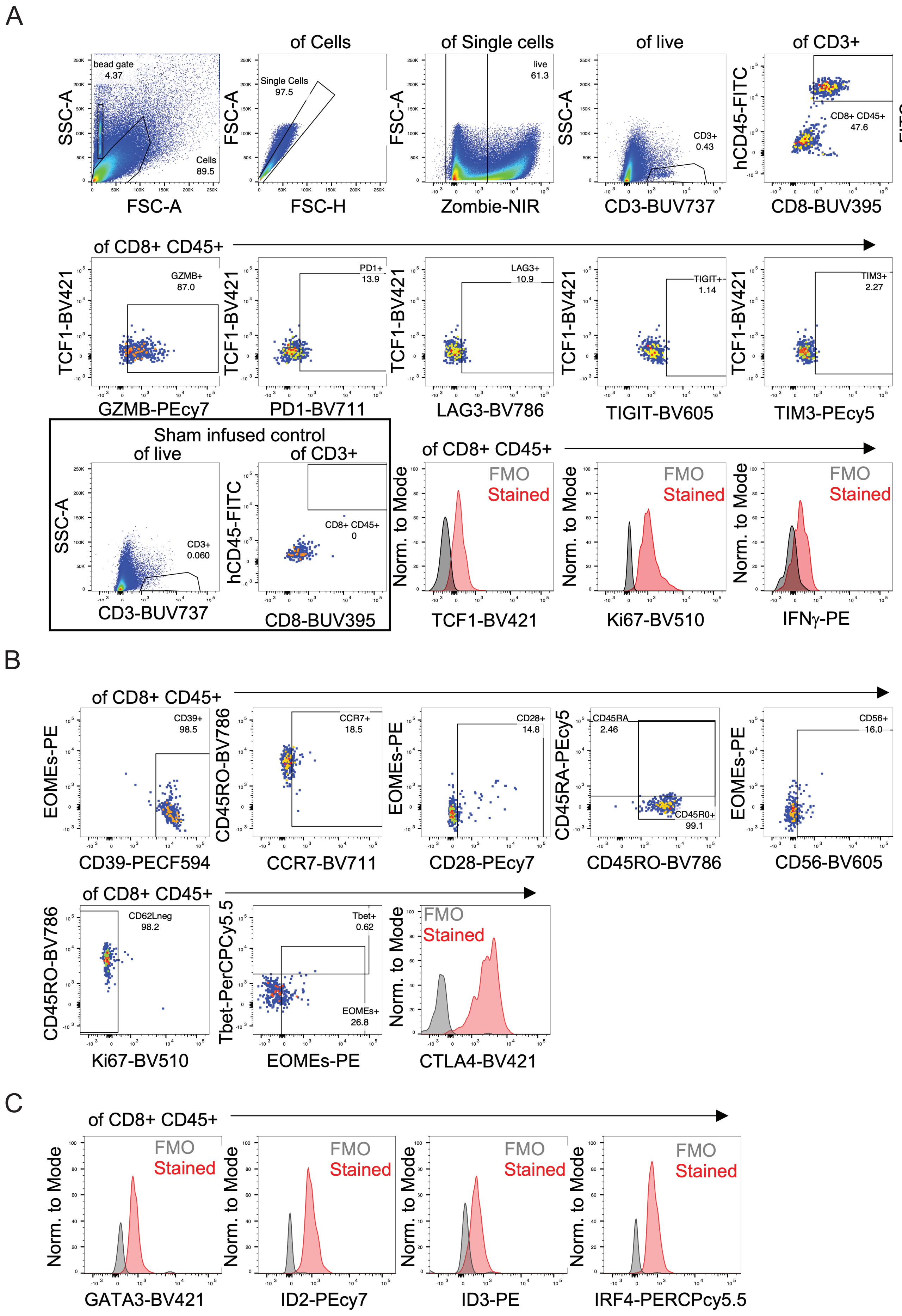
A
of Cells
of Single cells
of live
of CD3+
of bead gate
bead gate
4.37
Cells
89.5
Single Cells
97.5
live
61.3
CD3+
0.43
CD8+ CD45+
47.6
beads
82.2
SSC-A
FSC-A
FSC-H
Zombie-NIR
CD3-BUV737
hCD45-FITC
CD8-BUV395
FITC
BUV395
of CD8+ CD45+
GZMB+
87.0
PD1+
13.9
LAG3+
10.9
TIGIT+
1.14
TIM3+
2.27
TCF1-BV421
GZMB-PEcy7
PD1-BV711
LAG3-BV786
TIGIT-BV605
TIM3-PEcy5
Sham infused control
of live
of CD3+
CD3+
0.060
CD8+ CD45+
0
FMO
Stained
Norm. to Mode
TCF1-BV421
Ki67-BV510
IFNγ-PE
B
of CD8+ CD45+
CD39+
98.5
CCR7+
18.5
CD28+
14.8
CD45RA
2.46
CD45R0+
99.1
CD56+
16.0
EOMEs-PE
CD45RO-BV786
CD45RA-PEcy5
CD39-PECF594
CCR7-BV711
CD28-PEcy7
CD45RO-BV786
CD56-BV605
CD62Lneg
98.2
Tbet+
0.62
EOMEs+
26.8
Tbet-PerCPCy5.5
Ki67-BV510
EOMEs-PE
CTLA4-BV421
C
GATA3-BV421
ID2-PEcy7
ID3-PE
IRF4-PERCPcy5.5

**Figure S19**

**Figure S19. Representative gating strategies for data in figures 5 and S18.**

(**A-C**) Dissociated single cell suspensions from tumors at day 19 post treatment were gated for live singlets, followed by identification of CD3+, CD8+, and human (h)CD45+ to identify CAR T cells as shown. Identified T cells were then stained with three panels [see methods: panels 1 (**A**), 2 (**B**) and 3 (**C**) for indicated targets]. Fluorescence minus one (FMO) controls were used to confirm appropriate compensation and determine positivity for each marker, relevant markers are shown in histograms comparing FMO vs antibody staining.

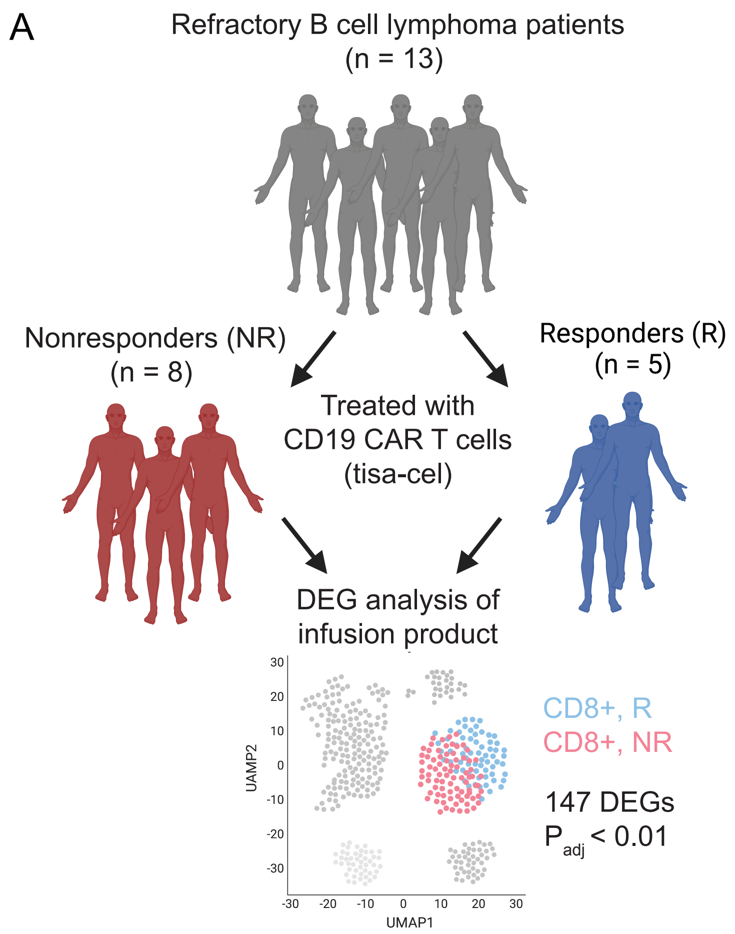
A
Refractory B cell lymphoma patients
(n = 13)
Nonresponders (NR)
(n = 8)
Responders (R)
(n = 5)
Treated with
CD19 CAR T cells
(tisa-cel)
DEG analysis of
infusion product
UAMP2
UMAP1
CD8+, R
CD8+, NR
147 DEGs
$P_{adj} < 0.01$

### Figure S20

**Figure S20. BATF3 programs a transcriptional response associated with positive clinical outcome to CAR T cell therapy.**

**(A)** Schematic of transcriptomic comparison between infused CD8+ CAR T cells of non-responders (NR, red) and responders (R, blue).

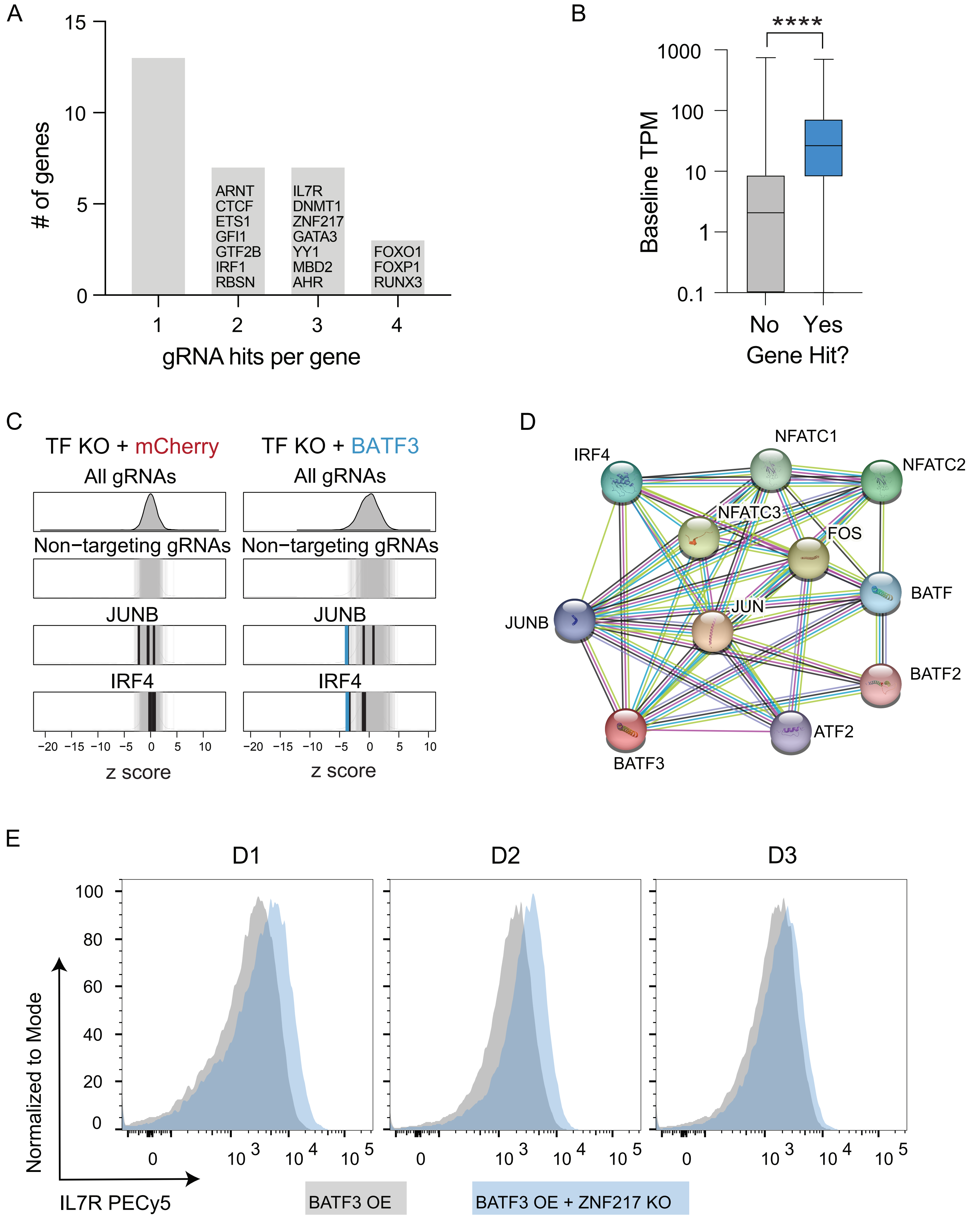
A
### of genes
gRNA hits per gene
ARNT
CTCF
ETS1
GFI1
GTF2B
IRF1
RBSN
IL7R
DNMT1
ZNF217
GATA3
YY1
MBD2
AHR
FOXO1
FOXP1
RUNX3
B
****
Baseline TPM
No
Yes
Gene Hit?
C
TF KO + mCherry
TF KO + BATF3
All gRNAs
Non−targeting gRNAs
JUNB
IRF4
z score
D
IRF4
NFATC1
NFATC2
NFATC3
FOS
BATF
JUNB
JUN
BATF2
BATF3
ATF2
E
D1
D2
D3
Normalized to Mode
IL7R PECy5
BATF3 OE
BATF3 OE + ZNF217 KO

**Figure S21**

**Figure S21. CRISPRko screens reveal co-factors of BATF3 and novel targets for cancer immunotherapy.**

**(A)** Number of gRNA hits ($P_{adj} < 0.01$) per gene in the CRISPRko screen without BATF3 OE. Genes with at least 1 enriched gRNA were included in this plot.

**(B)** Boxplot of baseline expression of genes stratified based on whether they were hits in the CRISPRko screen without BATF3 OE. Genes with an FDR < 0.01 based on mageck gene-level analysis were classified as hits.

**(C)** z scores of gRNAs for JUNB and IRF4 in mCherry (left) and BATF3 (right) screens. Enriched gRNAs ($P_{adj} < 0.01$) are labeled in blue and non-targeting gRNAs are labeled in gray.

**(D)** Predicted functional protein association network of BATF3 using STRING.

**(E)** Effect of ZNF217 knockout on IL7R expression in CD8+ T cells across three donors with BATF3 OE.
